# Harnessing *in vivo* Synthesis of Bioactive Multiarylmethanes in *Escherichia coli* via Oxygen-Mediated Free Radical Reaction Induced by Simple Phenols

**DOI:** 10.1101/2024.07.29.605594

**Authors:** Donglou Wang, Jiangbo He, Yonghong Chen, Boran Liu, Zhuang Wu, Xuerong Pan, Xuemei Niu

**Affiliations:** State Key Laboratory for Conservation and Utilization of Bio-Resources & Key Laboratory for Microbial Resources of the Ministry of Education, School of Life Sciences, Yunnan University, Kunming 650091, P. R. China; Kunming Key laboratory of Respiratory disease, Kunming University, Kunming, 650214, P. R. China

**Keywords:** Multiarylmethanes, Arthrocolins, *in vivo* Synthesis, *Escherichia coli*, Oxygen-mediated free radical reaction, Antitumor

## Abstract

**Background:** Xanthenes and multi-aryl carbon core containing compounds represent different types of complex and condensed architectures that have impressive wide range of pharmacological, industrial and synthetic applications. Moreover, indoles as building blocks were only found in naturally occurring metabolites with di-aryl carbon cores and in chemically synthesized tri-aryl carbon core containing compounds. Up to date, rare xanthenes with indole bearing multicaryl carbon core have been reported in natural or synthetic products. The underlying mechanism of fluores-cein-like arthrocolins with tetra-arylmethyl core were synthesized in an engineered *Escherichia coli* fed with toluquinol remained unclear.

**Results:** In this study, the Keio collection of single gene knockout strains of 3901 mutants of *E. coli* BW25113, together with 14 distinct *E. coli* strains, was applied to explore the origins of endogenous building blocks and the biogenesis for arthrocolin assemblage. Deficiency in bacterial respiratory and aromatic compound degradation genes *ubiX*, *cydB*, *sucA* and *ssuE* inhibited the mutant growth fed with toluquinol. Metabolomics of the cultures of 3897 mutants revealed that only disruption of *tnaA* involving in transforming tryptophan to indole, resulted in absence of arthrocolins. Further media optimization, thermal cell killing and cell free analysis indicated that a non-enzyme reaction was involved in the arthrocolin biosynthesis in *E. coli*. Evaluation of redox potentials and free radicals suggested that an oxygen-mediated free radical reaction was responsible for arthrocolins formation in *E. coli*. Regulation of oxygen combined with distinct phenol derivatives as inducer, 31 arylmethyl core containing metabolites including 13 new and 8 biological active, were isolated and characterized. Among them, novel arthrocolins with *p*-hydroxylbenzene ring from tyrosine were achieved through large scale of aerobic fermentation and elucidated x-ray diffraction analysis. Moreover, most of the known compounds in this study were for the first time synthesized in a microbe instead of chemical synthesis. Through feeding the rat with toluquinol after colonizing the intestines of rat with *E. coli*, arthrocolins also appeared in the rat blood.

**Conclusion:** Our findings provide a mechanistic insight into *in vivo* synthesis of complex and condensed ar-throcolins induced by simple phenols and exploits a quinol based method to generate endoge-nous aromatic building blocks, as well as a methylidene unit, for the bacteria-facilitated synthesis of multiarylmethanes.

---

Fluorescein-like compounds with multiphenylmethyl cores represent a unique family of complex and dense architectures that have attracted tremendous attention in organic synthesis for their importance in medicinal chemistry, materials science and potential drug candidates.^1–3^ In particular, the indole containing methyl cores are prevalent subunits in most of biologically active molecules, such as, cassigarol B, letrozole, vorozole, paraphenyl-substituted di-indolylmethanes, and triphenylmethylamides.^3,4^ Nonetheless, only di-aryl methyl cores bearing indole have been reported in natural metabolites and most of tri-aryl methyl cores bearing indole have been synthesized chemically.^5–9^ Recently, we reported an engineered *E. coli* E_BL21_-FPP*-Ao276* that harbored a fungal gene *Ao276* encoding toluquinol prenyltransferase,^10–12^ when fed with toluquinol (**1**) or hydroxyquinone (**2**), could produce an unprecedented type of 9-phenyl-9H-xanthene-2,7-diols with multi-aryl carbon cores, arthrocolins (**3**–**9**) that shared the similar key features of fluorescein dyes (Fig. 1).^10^ Arthrocolins A–D (**3**–**6**) were the first semi-natural multiaryl methanes bearing indole and fluorescein-like compounds produced in microorganism. Importantly, arthrocolins A–C (**3**–**5**) displayed potential inhibitory activities towards paclitaxel-resistant cell A549/Taxol. However, the underlying chemical mechanism for arthrocolins synthesis in *E. coli* still remained largely unclear.

**Figure 1.**
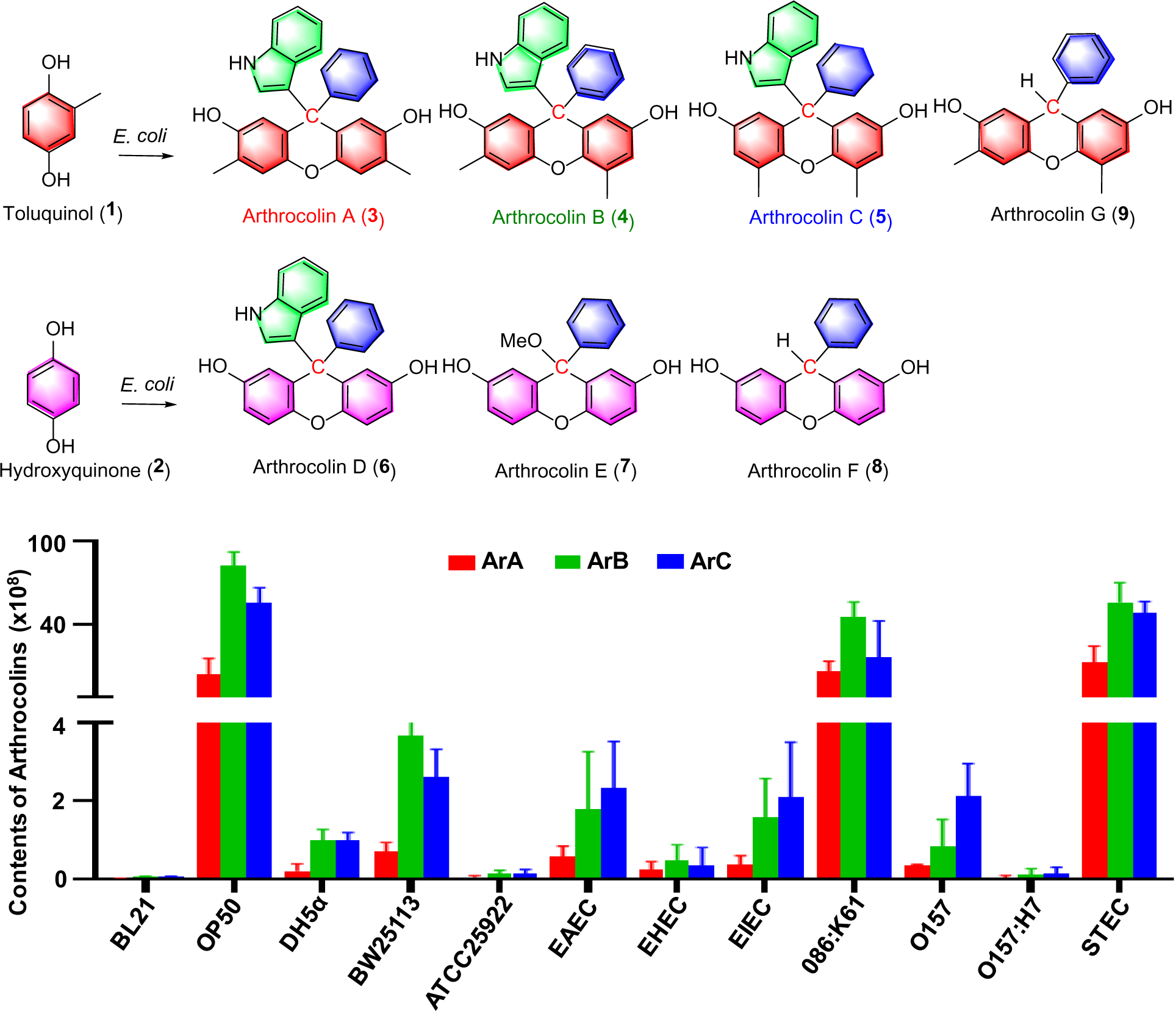
Top) Fluorescein-like multiarylmethanes, arthrocolins (**3**–**9**) produced in an engineering *E. coli* E_BL21_-FPP*-276* fed with toluquinol (**1**) and hydroquinone (**2**), respectively. Down) The contents of arthrocolins A– C (**3**–**5**) in 12 *E. coli* strains fed with toluquinol.

Here, we first cultivated and analyzed the fermentation cultures of twelve distinct *E. coli* strains including BW25113 and OP50, fed with toluquinol and found out that all the strains could yield arthrocolins on either Luria-Bertan (LB) and/or nutrient broth II (NB) media (Figs. 1 and S1). However, the arthrocolin contents in these strains were 100 times less than those in the engineering *E. coli* E_BL21_-FPP*-276*. To evaluate the underlying mechanism for arthrocolin synthesis in *E. coli*, we fermented and compared the metabolic profiles of 3901 mutants of *E. coli* BW25113 in the Keio collection of single-gene knockout strain library. Fed with toluquinol, 2591 mutants exhibited significantly decreased contents of arthrocolins, 1306 mutants displayed significantly increased contents, one mutant Δ*tnaA* couldn’t afford arthrocolins and four mutants Δ*cydB*, Δ*ubiX*, Δ*sucA* and Δ*ssuE* couldn’t grow on LB medium (Table S1). Gene *tnaA* encodes tryptophanase that degrades tryptophan (Trp) to indole in *E. coli*, indicating the indole unit in arthrocolins is derived from Trp. Additions of indole in Δ*tnaA* fed with toluquinol, recovered the arthrocolin production (Figs. S2A–2B). Moreover, additions of indole in WT strain strongly improve the arthrocolin contents, indicating that indole derived from Trp is a building block for the arthrocolin synthesis (Figs. S2A and 2C). Lack of four genes, encoding cytochrome bd ubiquinol oxidase subunit II (CydB), flavin prenyltransferase (UbiX), FMN reductase (SsuE), and 2-oxoglutarate dehydrogenase (SucA), inhibited the bacterial growth on toluquinol, suggesting that the four proteins could metabolize toluquinol similar to the fungal toluquinol prenyltransferase Ao276.^11,12^

Importantly, CydB was reportedly involved in aromatic compound degradation.^13,14^ Supplement experiment exhibited that all the aromatic amino acids, phenylalanine (Phe), tyrosine (Tyr), and Trp increased the arthrocolin contents in *E. coli* fed with toqluquinol. Among them, Phe was the far best inducer and Trp second (Fig. S3). Further supplement experiment revealed that benzal-dehyde was much stronger than benzyl acid in boosting the arthrocolin contents in *E. coli* fed with toluquinol, while benzyl alcohol made no significant effect (Fig. S4). These results suggest-ed that benzaldehyde mainly derived from Phe was the key aromatic unit linking two toluquinols. Comparison of the levels of aromatic amino acids in the twelve *E. coli* strains fed with toluquinol revealed that the strain OP50 with the highest yields of arthrocolins had the highest levels of both benzyl acid and indole, confirming that Phe and Trp were the precursors for arthrocolin synthesis in *E. coli* (Fig. S5).

KEGG pathway analysis indicated that the 2591 genes whose deficiency caused the reductions in arthrocolin contents were mainly enriched in carbon metabolism, glycolysis/gluconeogenesis, starch and sucrose metabolism and pentose phosphate pathway (Fig. S6). Further additions of glycerol, glucose, maltose, galactose, fructose, D-Sorbitol, or lactose indeed inhibited *E. coli* to synthesize arthrocolins (Fig. S7). Moreover, glucose also inhibited arthrocolin production in *E. coli* fed with toluquinol and indole (Figs. S8A–8B). Detailed experiments revealed that glucose treatment strongly inhibited indole levels but increased Trp levels in *E. coli* fed with toluquinol (Fig. S8C–D). Benzaldehyde couldn’t be detected in this study most likely due to its unstable nature. However, an increase in levels Phe and Tyr suggested that glucose also inhibited transformation of Phe to benzaldehyde (Figs. S8E–F).

Meanwhile, we found that toluquinol addition decreased pH values in *E. coli* and an increase in pH values increased the arthrocolin contents (Figs. 2A and S9). We first hypothesized that redox reduction potentials in *E. coli* might be involved in arthrocolins production^15^ because the addition of glucose strongly inhibited redox reduction potentials in *E. coli* WT all the times (Fig S10). However, comparison of redox reduction potentials of those 12 different *E. coli* strains fed with toluquinol revealed that the arthrocolin contents were not related to redox reduction potentials (Fig. 2B). Another evidence came from the evaluation of redox reduction potentials in the top ten productive mutants in Keio collection for arthrocolins (Fig. S11).

**Figure 2.**
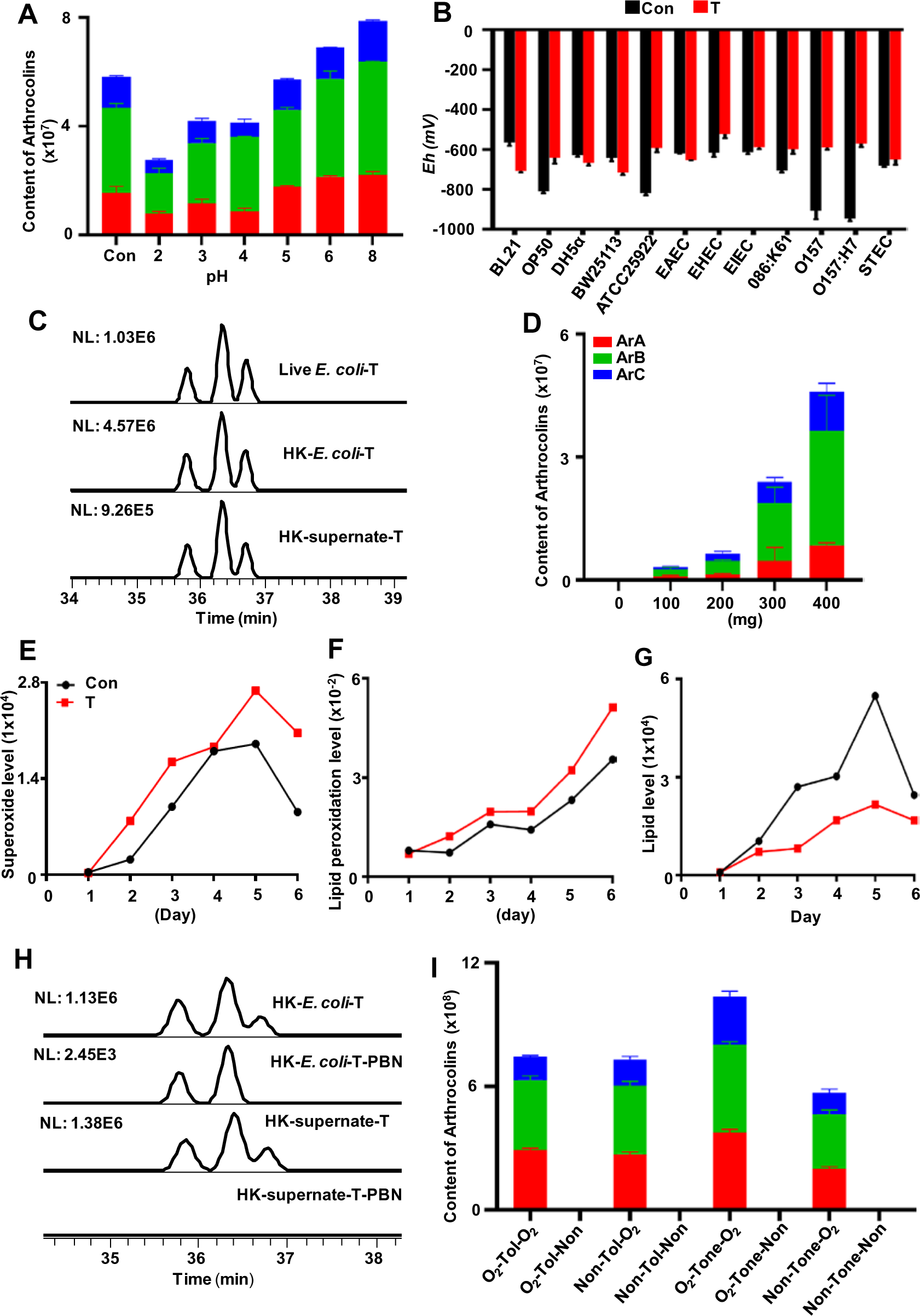
The synthesis of arthrocolins A–C in *E. coli* fed with toluquinol depended on oxygen-mediated free radical reaction. A) pH values were positively related to the contents of arthro-colins in *E. coli* fed with toluquinol. Ars A–C: arthrocolins A–C; T: toluquinol. Solvent used as control (Con). B) Comparison of redox potentials of twelve distinct *E. coli* strains indicated that the synthesis of arthrocolins in *E. coli* fed with toluquinol was not related to redox potentials. C) The bacteria, cultural broths and supernates of *E. coli* fed with toluquinol under heat killing treatment yielded arthrocolins. HK: heat killing treatment. D) The contents of arthrocolins de-pended on bacterial weights of *E. coli* under heat killing treatment. E–G) The effects of tolu-quinol on the levels of superoxide anions (E), lipid peroxidation (F) and lipids (G) in *E. coli*. H) A free radical capturing agent α-phenyl-tert-butylnitrone (PBN) inhibited the synthesis of A–C in *E. coli* WT fed with toluquinol. I) Comparison of the arthrocolin contents in *E. coli* WT treated with toluquinol and toluquinone under aerobic and anaerobic conditions. Tol: toluquinol; Tone: toluquinone; O_2_: aerobic conditions; Non: anaerobic conditions.

We assumed that the arthrocolin synthesis in *E. coli* fed with toluquinol might depend on non-enzyme reactions. *E. coli* OP50 strain cultivated on LB at 37°C for one-day was collected and put at 75°C for 90 min, and then fed with toluquinol at 30°C for 5h. Interestingly, these strains under heat treatment still had the ability to yield arthrocolins (Figs. 2C and S12). Even more, the supernatant from the fermentation broth with the bacteria removed also yielded arthrocolins after toluquinol addition (Figs. 2C and S12). Moreover, arthrocolin contents in *E. coli* fed with toluquinol increased with the bacterial weights (Figs. 2D and S13). These results suggested that the formation of arthrocolins in *E. coli* was independent on enzymes but positively related to bacterial contents.

Our recent study indicated that toluquinol addition could induce C5 or C6-methylsulfonyl substitution of toluquinol via free radical reaction in four dominant gut microbes, including *Enterococcus faecali* and *E. faecium*, *Stretpococcus thermophiles* and *Lactococcus lactis*.^16^ Interestingly, we found that toluquinol could induce significantly increased levels of superoxide anion and lipid peroxidation and largely decreased the lipid levels in *E. coli*, though ROS levels remained unchanged (Figs 2E–G and S14). Lipid peroxidation is the process in which radical extracts a hydrogen atom from the lipid leaving behind a lipid radical and setting off an oxygen-mediated chain reaction, which is the ultimate trigger of ferroptosis, a non-apoptotic cell death of iron-fueled Fenton reaction.^17–19^ Furthermore, toluquinol could increase Fe^2+^/Fe^3+^ ratios in and out of bacterial cells, in particular, the levels of Fe^2+^ and total free iron in all the six *E. coli* broths (Fig S15). Then, a free radical capturing agent α-phenyl-tert-butylnitrone (PBN)^20–22^ was applied to the heat-killed *E. coli* WT supernatants fed with toluquinol (Figs 2H and S16A). Indeed, no arthrocolins could be detected. Further experiment with mutant Δ*tnaA* fed with toluquinol and indole also indicated that PBN strongly inhibited arthrocolin synthesis, suggesting that free radical reaction was involved in arthrocolin synthesis in *E. coli* (Figs 2H and S16B).

Previous studies indicated that CydB, UbiX and SsuE were bacterial respiratory enzymes catalyzing oxidation–reduction reactions with using O_2_ as the final electron acceptor and SucA played a key role in bacterial aerobic growth.^23–25^ We assumed that O_2_ might be involved in arthrocolin synthesis. In addition, toluquinol is easily oxidized to toluquinone under O_2_ treatment. Therefore, both aerobic and anaerobic conditions were applied for the *E. coli* culture and the supplement of toluquinol or toluquinone, respectively. As expected, arthrocolins could be produced only when both compound addition and then bacterial cultivation were performed under aerobic condition (Fig 2I). Notably, toluquinone could induce the *E. coli* culture to produce more arthrocolin contents than toluquinol (Fig 2I). Further evaluation of the levels of toluquinol and toluquinone in *E. coli* fed with toluquinol indicated that no toluquinone was detected under glucose treatment, which inhibited arthrocolin synthesis in *E. coli* (Fig. S17). All the results indicate that O_2_ is an essential factor in arthrocolin synthesis in *E. coli* fed with toluquinol. Importantly, toluquinone, not toluquinol, is the key precursor for arthrocolin synthesis, consistent with the fact that quinones are attributed to the formation of hydroxyl radicals in the presence of iron and turn to radicals via Fenton chemistry.^16^

Based on the above conclusions, we tried to manipulate *E. coli* OP50 fed with toluquinol for synthesis of new arthrocolins since no arthrocolins derived from Tyr, one of three aromatic amino acids, were obtained. *E. coli* was cultivated in 70 L large fermenter with high speed stirring and toluquinol supplement was carried out sufficient aerobic conditions. Eventually, metabolic profiles and detailed chemical investigation led to isolation of 19 targeted metabolites (**3**–**5** and **9**–**26**, Figs. 1 and 3). These structures, including eleven new (Arthrocolins H–R, **10**–**17**, **20**, **24**, **26**), were characterized according to their HRMS, and 1D and 2D NMR data (Tables S2–S8 and Figs. S18–67). Among them, **11**–**12** were a pair of enantiomers which were separated with chiral columns. The absolute structure of **11** was established with X-ray diffraction analysis. Notably, most metabolites were multiarylmethanes of distinct types. Among them, compounds **10**–**12** were three new arthrocolins that harbored a building block, *p*-hydroxybenzene derived from Tyr, instead of benzene derived from Phe in all the known arthrocolins (**3**–**9**), suggesting that toluquinol also induced the formation of p-hydroxybenzene derived from Tyr as a building block in the bacterium. Compounds **10**–**16** were composed of toluquinol as a building block, while **17**–**26** not. Ten metabolites (**14**–**15** and **19**–**26**) harbored two indole units and four compounds (**21**–**23**, **26**) consisted of three indole units, suggesting non-selectivity and contact probability of free radical reaction induced in *E. coli* fed with toluquinol.

**Figure 3.**
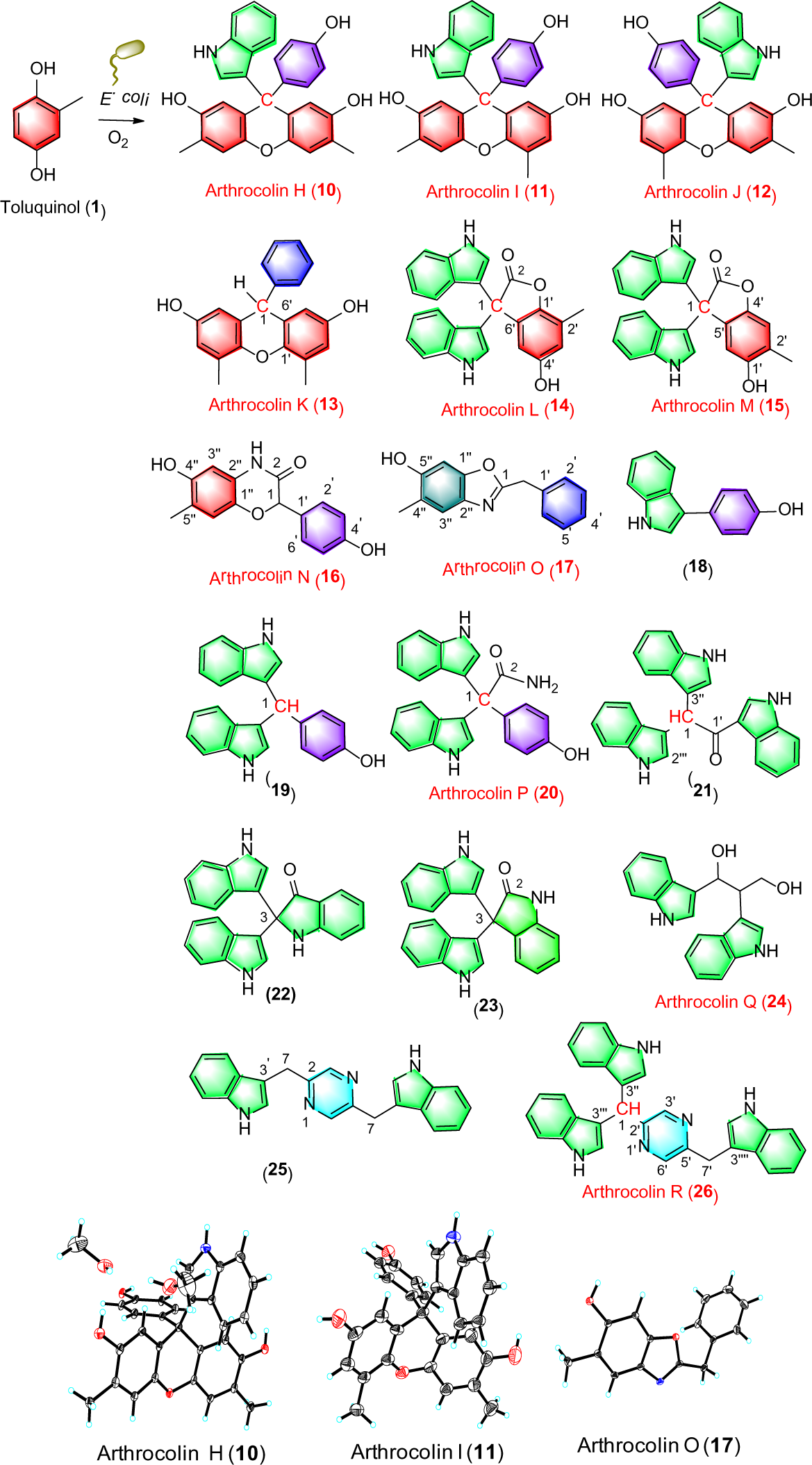
Chemical structures of metabolites **10**–**26** from *E. coli* fed with toluquinol (**1**), and X-ray diffraction analysis of compounds **10**–**11** and **17**.

Because metabolism of xenobiotics is a major source of free radical reaction,^26^ a number of phenol derivatives (**27**–**56**) were applied to evaluate their effects on the assembling capability of *E. coli* for multiarylmethanes (Figs. S68–S72). Among them, resorcinol (**27**) was the major building block for chemical synthesis of xanthenes in artificial chemical reactor.^3^ Metabolic pro-files and detailed chemical investigation indicated that the extra peaks in *E. coli* fed with resorcinol (**27**) were assigned to three aromatic metabolites **57**–**59** (Fig. 4), which were characterized according to their HRMS and 1D and 2D NMR data. The NMR spectra of **57**–**59** (Table S9 and Figs. S73–S77) all displayed the characteristic signals for a 6-substituted resorcinol (**27**). Metabolite **57** was a methane substituted with two identical resorcinol units whose signals completely overlapped due to the symmetry of the molecule. Compounds **58** and **59** were two methanones with a 4-hydroxyphenyl unit in **58** and an indole unit in **59**, respectively. Among them, **59** is a new compound. Obviously, resorcinol (**27**) also induced metabolisms of the three endogenous aromatic amino acids in *E. coli*, resulting in production of aryl building blocks, Tyr-derived 4-hydroxyphenyl and Trp-derived indole units. Importantly, few examples of the methylidene be-tween two identical moieties via C-C-C bonds, have been described in natural products except a recent study that reported dimeric aspidofractinine alkaloids, pleiokomenines A and B isolated from the stem bark of *Pleiocarpa mutica*, linked by a methylidene bridge.^27^ The findings suggested that resorcinol (**27**) could improve *in vivo* synthesis of methylidene.

**Figure 4.**
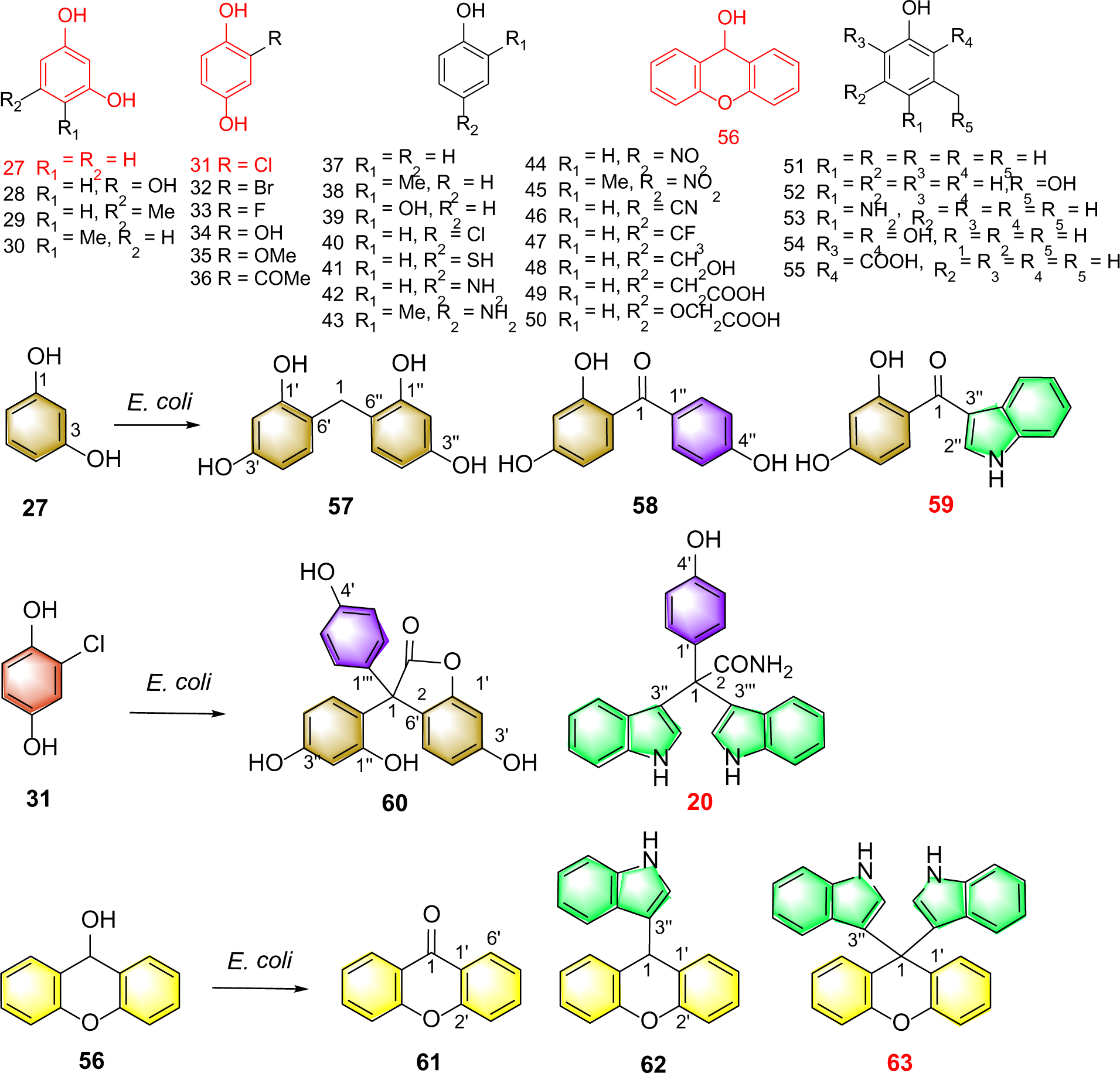
Evaluation of phenyl derivatives on the in vivo synthesis of multiarylmethanes and chemical structures of metabolites **57**–**63** from *E. coli* fed with resorcinol (**27**), 2-chlorobenzene-1,4-diol (**31**), and 9*H*-xanthen-9-ol (**56**), respectively.

Though 2-chloro-hydroquinone (**31**) failed to induce arthrocolin synthesis in *E. coli*, two multiarylmethanes (**60** and **20**) also with Tyr-derived 4-hydroxyphenyl or/and Trp-derived indole units, including compound **20**, were isolated and identified according to their 1D and 2D NMR data (Table S5 and S10 and Figs. S53–S57). Moreover, compound **60** harbored two resorcinol units, which might be derived from 2-chloro-hydroquinone (**31**) most likely via free radical-mediated dechlorination reaction. Compound **56**, 9*H*-xanthen-9-ol, induced formation of three major target metabolites (**61**–**63**) in *E. coli* (Table S10 and Figs. 78–82), which were character-ized as three xanthene derivatives, including a new metabolite 3,3’-(9*H*-xanthen-9,9-diyl)-bis(1*H*-indole) (**63**). Previous study revealed that **56** could spontaneously transform to **61** under air condition.^28^ The existence of two indolyl moieties in **63** might be derived from one indolyl addition to the ketone in **61** followed with the second indolyl substitution. All the results sug-gested that *E. coli* should be an unexploited biofactory for biosynthesis of indole bearing com-plex metabolites, in particular, condensed bis(indolyl)methanes. *E. coli* bacteria naturally live in the intestines of humans and many animals. Previous studies reported that toluquinol was a po-tential antitumor agent.^29,30^ To explore whether toluquinol could transform into arthrocolins in the intestine of animals, toluquinol was orally fed to rat for seven days, and then blood, heart, liver, spleen, kidney, stomach, small intestine of the rat were collected for HPLC-MS analysis. However, no arthrocolins were detected. Further analysis indicated that the rat had very low amounts of *E. coli* in their feces. Thus, *E. coli* strain was orally fed to rat for two weeks until *E. coli* strain was detected in the rat feces. Then toluquinol was orally fed to rat for seven days. Eventually, the major arthrocolins A–C (**3**–**5**), were detected in the blood of the rat while not in the heart, liver, spleen, kidney, stomach, small intestine (Fig. 5). All the results suggest that the synthesis of complex arylmethanes derived from simple phenols in gut microbe in animal via oxygen-mediated free radical reaction is practical.

**Figure 5.**
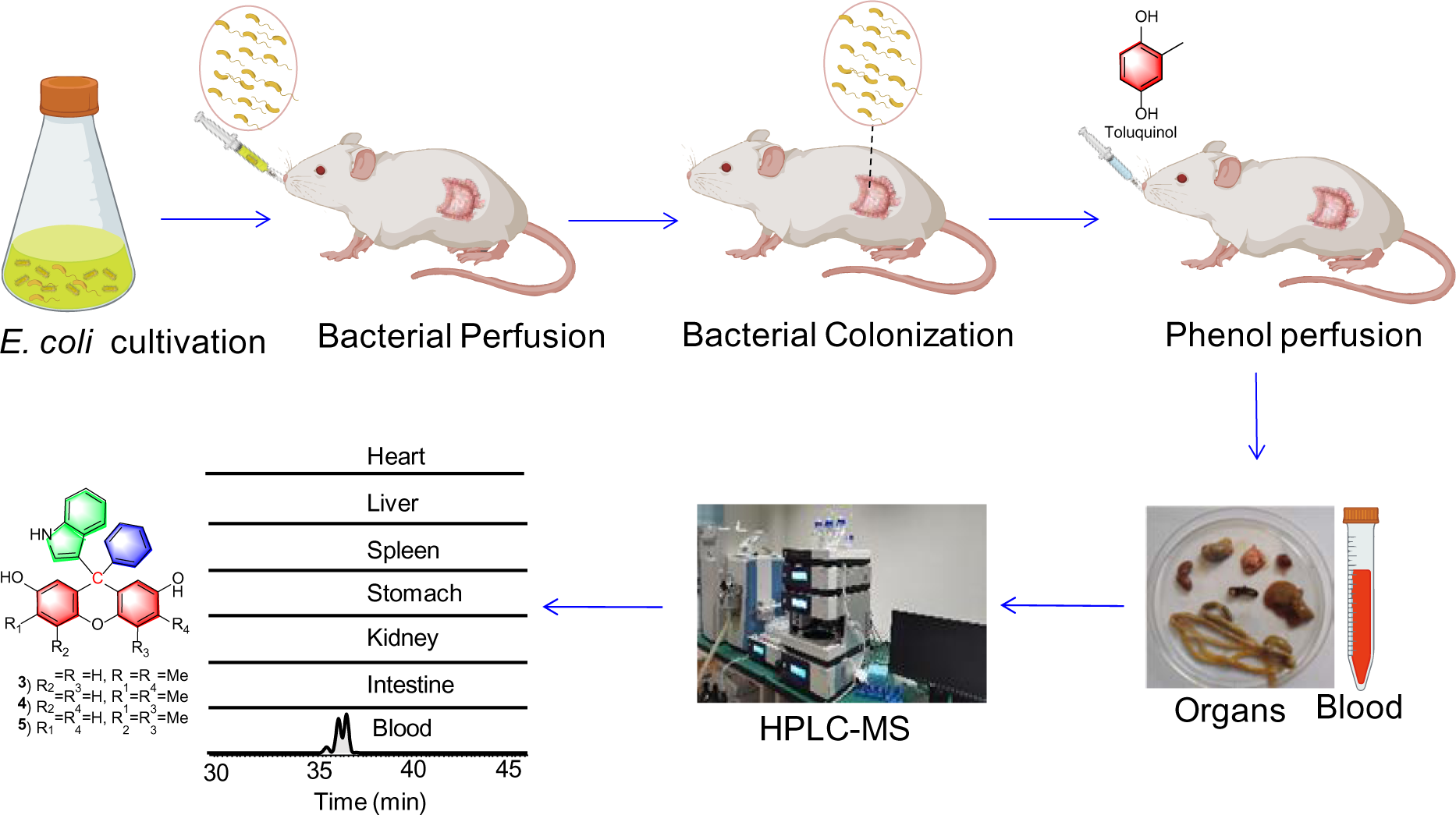
Synthesis of arthrocolins in rat fed with toluquinol after colonizing the intestines of rat with *E. coli*.

We first evaluated the effects of fifteen compounds, **10**–**15** and **18**–**26**, on four cancer cell lines, including human breast carcinoma cell MDA-MD-231, human colorectal carcinoma cells HCT116, human hepatocarcinoma cells SMMC-7721 and glioma cells U251. Six compounds, **15**, **18**–**23** displayed anticancer activities. Among them, compounds **15**, **19**, **21** and **23** showed strong anticancer activity against all the four cancer cell lines with IC_50_ values ranging from 6.5 to 10 μM (Table S11). However, arthrocolins **10**–**13**, with one extra hydroxyl group at phenyl ring showed no anticancer activity. We also evaluated anticancer activities of **20**, **60** and **62**–**63** against ten cancer cell lines, including human non-small cell lung cancer cell A549, human amelanotic melanoma cell A375, human osteosarcoma cell (HOS), human glioma cells U87, human gastric cancer cell SGC-7901, human acute lymphocyte leukemia T lymphocyte lineage (CEM-SS), human renal cell carcinoma cell 786-0, as well as MDA-MD-231, human colorectal carcinoma cells HCT116, and human hepatocarcinoma cells SMMC-7721 (Table S12). Compound **62** inhibited all ten cancer cells with IC_50_ values ranging from 2.2 to 9.1 μM and **63** inhibited seven cells with IC_50_ values ranging from 3.8 to 9.8 μM. Interestingly, **20** and **60** strongly promoted cancer cell viabilities in six of ten different tumor cell lines. In particular, A549 treated with compound **60** displayed concentration-dependent growth.

Here, our study investigated on the underlying mechanism of the synthesis of “unnatural” bio-active multi-aryl core containing metabolites in bacterium *E. coli*. We exploited a quinol based method via oxygen-mediated free radicals to directly generate aromatic building blocks, as well as methylidene unit, for the bacteria-facilitated synthesis of multiarylmethanes. Most of the known compounds in this study were for the first time synthesized in a microbe instead of chemical synthesis. These findings have valuable implications for the manipulation of oxygen-mediated free radical reactions in organisms to make use of endogenous building blocks and synthesize biological active metabolites with complex and condensed structures.

## Methods and Materials

### Cultivation of 12 *E. coli* strains fed with toluquinol

The fermentation steps of the *Escherichia coli* were as follows. The 12.5 mL overnight culture of *E. coli* strains were inoculated into 250 mL of Luria-Bertan (LB) or nutrient broth II (NB) media in a 500 mL flat-bottom flask. The cultures were incubated at 37°C for 6h (200 rpm., OD_600_ = 0.6–0.8) and then added with 5 mg toluquinol. After growth for 72 h at 28 °C (200 rpm.), the cultures were concentrated *in vacuo* to 50 mL and were extracted overnight with ethyl acetate (1:1 v/v). The organic fractions were evaporated to dryness to give residues. The dried organic residues were then dissolved in 500 μL of methanol, filtered through 0.22 μm membranes, and further analyzed using UPLC-MS.

### Cultivation of 3901 *E. coli* strains fed with toluquinol

The fermentation steps of the *Escherichia coli* were as follows. The 50 μL of *E. coli* Knockout strains were inoculated into 5 mL of LB medium containing 100 μg/mL Ampicillin and 1mM/L toluquinol in a 7 mL 96 deep hole plate. After growth for 72 h at 37 °C (200 rpm.). Then were extracted overnight with ethyl acetate (1:1 v/v). The organic fractions were evaporated to dryness to give residues. The dried organic residues were then dissolved in 100 μL of methanol, filtered through 0.22 μm membranes, and further analyzed using UPLC-MS.

### Cultivation of *E. coli* under different conditions

#### Different pH conditions

The pH of the medium is adjusted to 2-8 using sodium phosphate dibasic-citric acid buffer, the LB liquid medium with pH = 7 was set as control group, and the other pH values were set as ex-perimental group. The 12.5 mL overnight culture of *E. coli* strains were inoculated into 250 mL of Luria-Bertan (LB) media in a 500 mL flat-bottom flask. The cultures were incubated at 37°C for 6h (200 rpm., OD_600_ = 0.6–0.8) and then added with 5 mg toluquinol. After growth for 72 h at 28 °C (200 rpm.), the cultures were concentrated *in vacuo* to 50 mL and were extracted overnight with ethyl acetate (1:1 v/v). The organic fractions were evaporated to dryness to give residues. The dried organic residues were then dissolved in 500 μL of methanol, filtered through 0.22 μm membranes, and further analyzed using UPLC-MS.

#### Medium supplemented with precursors and carbon sources

Potential precursors Indoles, L-Phenylalanine, Tyr, Trp, benzyl alcohol, benzaldehyde or benzoic acid and exogenous carbon sources such as glycerol, glucose, maltose, galactose, fructose, D-sorbitol or lactose were added respectively to LB medium. Then 12.5 mL overnight culture of *E. coli* OP50 were inoculated into 250 mL of Luria-Bertan (LB) media in a 500 mL flat-bottom flask. The cultures were incubated at 37°C for 6h (200 r.p.m., OD_600_ = 0.6–0.8) and then added with 5 mg Toluquinol. After growth for 72 h at 28 °C (200 r.p.m.), the cultures were concentrated *in vacuo* to 50 mL and were extracted overnight with ethyl acetate (1:1 v/v). The organic fractions were evaporated to dryness to give residues. The dried organic residues were then dissolved in 500 μL of methanol, filtered through 0.22 μm membranes, and further analyzed using UPLC-MS.

### Heat-killing experiments

The 12.5 mL overnight culture of *E. coli* strains were inoculated into 250 mL of Luria-Bertan (LB) media in a 500 mL flat-bottom flask. The cultures were incubated at 37°C for 1 day (200 rpm.). Bacteria and supernatant were collected by centrifugation at 4000 rpm for 5 min, Bacteria washed with PBS for 3 times, and then resuspended with 5 mL of PBS, then Bacteria and Supernatant put at 75°C for 90 min, and then fed with toluquinol at 30°C for 5h, were extracted overnight with ethyl acetate (1:1 v/v). The organic fractions were evaporated to dryness to give residues. The dried organic residues were then dissolved in 100 μL of methanol, filtered through 0.22 μm membranes, and further analyzed using UPLC-MS.

The bacteria were weighed and resuspended with PBS at a final concentration of 100 mg/mL, then 0,1,2,3,4 mL bacterial suspension were added into 15 mL centrifuge tube, and PBS was added to fill up to 5 mL. The bacterial contents were 0,100,200,300,400 mg respectively. The bacterial solutions were put at 75°C for 90 min, and then fed with toluquinol at 30°C for 5h, were extracted overnight with ethyl acetate (1:1 v/v). The organic fractions were evaporated to dryness to give residues. The dried organic residues were then dissolved in 100 μL of methanol, filtered through 0.22 μm membranes, and further analyzed using UPLC-MS.

### Fermentation of *E. coli* under oxygen influx

The bioreactor used was a 50-L (working volume) stirred tank bioreactor (LP351, Bioengineering AG, Switzerland). Fermentation was carried out in two stages, a biomass accumulation stage (Stage 1) and a polymer biosynthesis stage (Stage 2). The first stage of fermentation was performed at 37°C with an initial agitation rate of 200 rpm and an aeration rate of 5.0 L min^−1^, and pH was maintained at 7.0 by automated addition of concentrated NH_4_OH (17 M) as monitored by an internal pH meter. Dissolved oxygen was maintained at 50% air saturation controlled by agitation and air (compressed atmospheric) sparging cascades (200 to 950 rpm, then 5.0 to 20.0 L min^−1^ air flow). Foaming was controlled by automated addition of Antifoam B (Sigma-Aldrich, 0.05% v/v unless otherwise noted). OD_600_ measurements were made on 1 mL culture aliquots each hour using Multimode microplate reader(Molecular Devices). After approximately 6 h of growth in Stage 1 (OD_600_ > 3), Then the precursor toluquinol was added until the final concentration was 1 mM. Stage 2 was initiated by decreasing the fermentation temperature to 28°C, The other conditions were unchanged, and the fermentation time was 72 h (Scheel *et al.*, 2021).

### Metabolite Profiles

Twenty μL of the solution was injected to the LC-DAD/MS system for analysis. LC-MS analyses were performed on a Q Exactive Focus UPLC-MS (Thermofisher, USA) with a PDA detec-tor and a Obitrap mass detector (Shiseido, 5 μm, 4.6 × 250 mm, CAPCELL PAK C18 column) using positive and negative mode electrospray ionization. The total flow rate was 1 mL/min; mobile phase A was 0.1% formic acid in water; and mobile phase B was 0.1% formic acid in acetonitrile. The column temperature was maintained at 40 °C. The injection volume for the extracts was 25 μL. The liquid chromatography (LC) conditions were manually optimized on the basis of separation patterns with the following gradient: 0−2 min, 10% B; 10 min, 25% B; 30 min, 35% B; 35 min, 90% B; 36 min, 95% B; 40 min, 90% B; 40.1 min, 10% B; and 45 min, 10%B. UV spectra were recorded at 204–400 nm.

### Transcriptional analysis

Total RNAs were extracted from about 1×10^10^ *E. coli OP50* between the control group and tolu-quinol groups using the RNeasy Plus kit (Qiagen; catalog no. 74136) following the manufactur-er’s instructions. The concentration of the extracted RNA samples was determined as described above, and the integrity of the RNA was examined by the RNA integrity number (RIN) using an Agilent 2100 bioanalyzer (Agilent, Santa Clara, USA).

RNA library was validating on the Agilent Technologies 2100 bioanalyzer for quality control. The double stranded PCR products above were heated denatured and circularized by the splint oligo sequence. The single strand circle DNA (ssCir DNA) was formatted as the final library. The final library was amplified with phi29 (Thermo Fisher Scientific, MA, USA) to make DNA nanoball (DNB) which had more than 300 copies of one molecular, DNBs were loaded into the patterned nanoarray and single end 50 bases reads were generated on BGISEQ500 platform (BGI-Shenzhen, China). Three biological replicates were analyzed for each sample. Low quality reads (Phred≤20) and adaptor sequences were filtered out, and the Q20, Q30 and total raw reads of clean date were calculated. Transcript abundances (FPKM) were provided by BGI after se-quencing and differentially expressed genes were calculated using R software. All genes with P-value≤0.05 and log2(fold_change) ≥1were considered significantly differentially expressed and used for volcano plotting, GO terms and KEGG pathway enrichment analysis.

### Physiological bioassays

pH values: A pH meter was used to measure the pH values during fermentation process.

Redox potentials: bacteria were collected by centrifugation at 4000 rpm for 5 min, washed with PBS for 3 times, and then resuspended with 5-10 ml of M9 buffer, after adjusting OD600 consistently, Redox potential detection was performed using a redox potential detector.

ROS and superoxide levels: ab139476 ROS/Superoxide Detection Assay Kit was used for ROS and superoxide levels. Bacteria were collected by centrifugation at 4000 rpm for 5 min, washed with PBS for 3 times, and then resuspended with 1mL of PBS. Add 4μLROS/Superoxide Detection Mix and incubate at room temperature in the dark for 30-60 minutes. bacteria were washed twice with 1× wash buffer, resuspended in 500 μL PBS, and analyzed by flow cytometry. The ROS detection channel is Ex/Em= 490/525 nm). The superoxide detection channel set to Ex/Em =550/620nm.

Lipid contents: Lipid content detection kit (AB242307) was used. Bacteria were collected by centrifugation at 4000 rpm for 5 min, washed with PBS for 3 times,and then resuspended with 200 μL of 1 × fat detection fluorescent dye then Incubate at 37°C for 15min in the dark. Then bacteria were washed twice with PBS and resuspend the bacteria using 500 μL of PBS, analysis was performed using Flow cytometry, and the detection fluorescence wavelength was set to Ex/Em= 380/650nm.

Lipid peroxidation levels: Lipid peroxidation test kit (AB243377) was used for evaluation of lipid peroxidation levels. Bacteria were collected by centrifugation at 4000 rpm for 5 min, washed with PBS for 3 times, and then resuspended with 90 μl of PBS. Add 10 μL of 1× Lipid Peroxidation Sensor and incubate the bacteria in 37 °C incubator for 30 min. Then bacteria were washed twice with PBS and resuspend the bacteria using 500 μL of PBS, and perform the assay using a flow cytometer with flow cytometry detection channels set to Ex = 488 nm, Em = 530 nm (FITC) and 572 nm (PE).

Iron contents: An iron content detection kit (# I291, Dojindo) was used. Bacteria were collected by centrifugation at 4000 rpm for 5 min, washed with PBS for 3 times, and then resuspended with 5-10 ml of PB, after adjusting OD600 consistently, Aspirate 1 mL of suspension, centrifuge to remove the supernatant, and resuspend the bacteria with 500 μL iron assay buffer and homogenized using the homogenizer sitting on ice. Then, the mixture was centrifuged at 12000 rpm for 10 min at 4°C, and 100 μL supernatant was collected for detection. For extracellular iron level, 1 mL filtrate was collected and centrifuged at 12000 rpm for 10 min at 4°C. One hundred μL supernatant was collected for detection. Next iron assay buffer /reducer was added in to the collected supernatant, mixed, and incubated. Finally, the incubated solution with iron probe for 1 h in the dark was immediately measured on microplate reader at OD = 593 nm.

### General methods for targeted compound isolation and structural elucidation

The 50 L cultural broths of the mutants cultured on liquid medium were concentrated *in vacuo* and partitioned with acetyl acetate. The acetyl acetate parts were evaporated to dryness to give residue. The crude extracts of mutants was subjected to purification first with absorption resin D101, then by reversed-phase (RP18) flash chromatography eluting with H_2_O–MeOH (10–90%) with decreasing polarity through a continuous gradient, and subsequently by open-column chromatographies on Sephadex LH-20 with acetone, respectively. Silica gel 60 (Merck, 200-400 mesh) was used for column chromatography. Column chromatography was performed on 200-300 mesh silica gel (Qingdao Marine Chemical Factory, P. R. China). The TLC spots were detected by spraying the TLC plates with 20% (w/v) H_2_SO_4_ and then heating them on a hot plate.

The purified samples were structurally elucidated with NMR and MS data. NMR experiments were carried out on either a Bruker DRX-500 spectrometer with solvent as internal standard. MS were recorded on a VG-Auto-Spec-3000 spectrometer. High-resolution ESIMS data were measured on a Bruker Bio-TOF III electrospray ionization mass spectrometer. Optical rotations were measured on a Horiba-SEAP-300 spectropolarimeter. UV spectral data were obtained on a Shimadzu-210A double-beam spectrophotometer. IR spectra were recorded on a Bruker-Tensor-27 spectrometer with KBr pellets.

### Bioassay of metabolites against tumor-cell lines

Human cancer cell lines were purchased from the Kunming Cell Bank of the Chinese Academy of Science. human breast carcinoma cell MDA-MD-231, human colorectal carcinoma cells HCT116, human hepatocarcinoma cells SMMC-7721, glioma cells U251, human non-small cell lung cancer cell A549, human amelanotic melanoma cell A375, human osteosarcoma cell HOS, human glioma cells U87, human gastric cancer cell SGC-7901, human acute lymphocyte leukemia t lymphocyte lineage CEM-SS, human renal cell carcinoma cell 786-0, as well as MDA-MD-231, human colorectal carcinoma cells HCT116. Cells were all cultured in DMEM (Gibco, Carlsbad, CA, USA) supplemented with 10% fetal bovine serum (BI, Kibbutz Beit Haemek, Israel) and 1% penicillin/streptomycin (Gibco); A549 cells were cultivated in RPMI-1640 medium (Gibco). All cells were cultured at 37 ◦C in a 5% CO2 humidified environment. For the cell viability assays, the cells were plated at a density of 5000 cells/well in 96-well plates and incubated overnight. The cells were then treated with CCK-8 for 72 h and assessed using a microplate reader at 450 nm.

Ten human cancer cell lines (786-0, MD-MB-231, U87, SMCC-7721, HOS, HCT116, A549, SGC-7901, A375, CEM-SS) were selected for the assay. The tested cell lines were seeded in 96-well plates, and then the plates were incubated for 24 h at 37 °C in 5% CO_2_ incubator. Subsequently, the compounds were added at a dosage of 1 – 10 μM. After 72 h, CCK8 was added to the culture medium and the absorbance at 490 nm was measured with a microplate reader (Spectra Max plus 384, MD, USA). Each experiment was carried out in triplicate. The IC_50_ values were calculated with the Graphpad Prism 5.01 software. Taxol was used a positive control.

### Mice bioassay

This study was carried out in strict accordance with the recommendations of the Guide for the Care and Use of Laboratory Animals of the National Institutes of Health. The protocol was approved by the Ethics Committee of Yunnan University of Yunnan University (YNUCARE20210034). Male adult Sprague Dawley (SD) rats weighing 180–220 g, aged 8 weeks, were purchased from the SPF Animal Experiment Center of Kunming Institute of Zoology, Chinese Academy of Science (KIZ, CAS, China). All animals were housed in filter-top cages with autoclaved bedding, environment with temperature of 20–25°C, humidity of 65–69 %, and a 12 h light/dark cycle, and fed with autoclaved food and water ad libitum.

The 50 mL overnight culture of *E. coli* OP50 were collected, washed with PBS for 3 times, and then resuspended with PBS. The bacteria concentrations were adjusted to 10^6^ cells/mL. The rats were perfused with 0.5 mL bacterial suspension by intragastric injection every day for two weeks. Then 10 mg toluquinol was given daily for 7 days. The rats were anesthetized with isoflu-rane and euthanized by exsanguination via cardiac puncture. The blood was collected in EDTA tubes, and plasma samples were obtained by centrifugation (3000x g for 10 min at 4°C) and stored at −80°C until analysis. Different tissues and organs (heart, liver, spleen, kidney, stomach, small intestine) were removed from the rats and placed in pre-weighed tubes. The collected tis-sues were washed with physiological serum at 37 °C and placed between absorbent paper to re-move the rest of the serum. In the case of the intestine, the luminal contents were first removed by applying pressure with the index finger and thumb, and then the lumen was washed by perfus-ing with the physiological serum at 37°C.^31^ It was then placed between absorbent paper to re-move the remains of the washings. In order to avoid compound oxidation, the collected tissues were kept on ice at all times. Finally, the samples were weighed and stored at −80°C until analy-sis. For each group, three animals were used.^32^

The plasma samples were thawed and centrifuged (11,000× *g* at 4 °C at 10 min). Plasma (100 µL) was mixed with cold ACN containing 2% formic acid in a ratio of 1:5 (*v*/*v*) to precipitate proteins, homogenized for 1 min and kept at −20 °C for 20 min. The samples were then centri-fuged (11,000× *g* at 4 °C for 10 min), and finally, 100 µL of each organic phase was transferred to vials for UPLC-MS analysis. Three replicates were evaluated for each sample.

The tissues were homogenized with a Glass homogenizer in a 1:1 ratio (*w*/*v*) with water: ACN (1:1 (*v*/*v*) with 0.1% ascorbic acid). The samples were sonicated for 5 min in an ice bath and shaken for 1 min. Each homogenate was centrifuged for 10 min (11,000× *g* at 4 °C). A volume of 100 µL of the upper layer was blended with cold ACN containing 2% formic acid in a 1:3 ratio (*v*/*v*) in order to precipitate the proteins (Polson *et al*., 2003). The samples were homogenized for 1 min and kept at −20 °C for 20 min before centrifugation (11,000× *g* at 4 °C for 10 min). Finally, 100 µL of supernatant was transferred to vials for UPLC-MS analysis. Three replicates were evaluated for each sample.

## Ethics approval and consent to participate

This study was carried out in strict accordance with the recommendations of the Guide for the Care and Use of Laboratory Animals of the National Institutes of Health. The protocol was ap-proved by the Ethics Committee of Yunnan University of Yunnan University (YNUCARE20210034).

## Consent for publication

Not applicable.

## Availability of data and materials

All supplementary figures (Figures S1-S82) and tables (Tables S1-S12) mentioned in this report were provided in supporting information.

## Authors’ contributions

M.N., D. W., Y. C. and J. H conceived the study and designed scientific objectives. X.M.N. led the project and provided funding support and experimental platform. D. W., Y. C. and J. H analyzed the data. M.N., D. W., Y. C. and J. H prepared the manuscript, and provided the materials and performed the experiments. J.H., Z.W., and B. L. contributed to experiments. M.N. and D.W. wrote the main manuscript text and prepared all the figures. All authors reviewed the manuscript.

## Acknowledgements

We appreciated Keqin Zhang for providing nice suggestion.

## Authors’ information

Donglou Wang, Yonghong Chen, Boran Liu, Zhuang Wu, Xuerong Pan, Xuemei Niu: State Key Laboratory for Conservation and Utilization of Bio-Resources & Key Laboratory for Microbial Resources of the Ministry of Education, School of Life Sciences, Yunnan University, Kunming 650091, P. R. China. Jiangbo He: Kunming Key laboratory of Respiratory disease, Kunming University, Kunming, 650214, P. R. China.

