## Supporting information for "Harnessing *in vivo* Synthesis of Bioactive Multiarylmethanes in *Escherichia coli* via Oxygen-Mediated Free Radical Reaction Induced by Simple Phenols"

**Experimental procedures**

1. Cultivation of 12 *E. coli* different strains fed with toluquinol

2. Cultivation of 3901 *E. coli* strains of the Keio collection fed with toluquinol

3. Cultivation of *E. coli* under different conditions

   3.1 pH

   3.2 Precursors and carbon sources

4. Heat-killing experiments

5. Fermentation of *E. coli* under oxygen influx

6. Mice bioassay

7. Metabolite Profiles

8. Transcriptional analysis

9. Physiological bioassays

   9.1. pH values

   9.2. Redox potentials

   9.3 ROS and superoxide levels

   9.4 Lipid contents:

   9.5. Lipid peroxidation levels

   9.6 Iron contents:

10. General methods for Targeted compound isolation and structural elucidation

   10.1 Metabolites from strain *E. coli* fed with Toluquinol (**1**).

   10.2 Metabolites from strain *E. coli* fed with resorcinol (**27**).

   10.3 Metabolites from strain *E. coli* fed with 2-chloro-benzene-1,4-diol (**31**)

   10.4 Metabolites from strain *E. coli* fed with 9*H*-xanthen-9-ol (**56**).

11. Bioassay of metabolites against ten tumor-cell lines.

**Supplementary Tables**

**Table S1.** Evaluation of arthrocolin synthesis in 3901 *E. coli* mutants.

**Table S2.** The $^{1}$H NMR (500 MHz) and $^{13}$C NMR (125 MHz) data of **10**–**13**.

**Table S3.** The $^{1}$H NMR (500 MHz) and $^{13}$C NMR (125 MHz) data of **14**–**15**.

**Table S4.** The $^{1}$H NMR (500 MHz) and $^{13}$C NMR (125 MHz) data of **16**–**17**.

**Table S5.** The $^{1}$H NMR (500 MHz) and $^{13}$C NMR (125 MHz) data of **19**–**20**.

**Table S6.** The $^{1}$H NMR (500 MHz) and $^{13}$C NMR (125 MHz) data of **21**–**23**.

**Table S7.** The $^{1}$H NMR (500 MHz) and $^{13}$C NMR (125 MHz) data of **24**.

**Table S8.** The $^{1}$H NMR (500 MHz) and $^{13}$C NMR (125 MHz) data of **25**–**26**.

**Table S9.** The $^{1}$H NMR (500 MHz) and $^{13}$C NMR (125 MHz) data of **57**– **59**.

**Table S10.** The $^{1}$H NMR (500 MHz) and $^{13}$C NMR (125 MHz) data of **60**, **62** and **63**.

**Table S11.** The effects of fifteen compounds, **10**–**15** and **18**-**26**, on four cancer cell lines, including human breast carcinoma cell MDA-MD-231, human colorectal carcinoma cells HCT116, human hepatocarcinoma cells SMMC-7721 and glioma cells U251.

**Table 12.** The $IC_{50}$ (μM) values of compounds **62**–**63** against ten cancer cells.

**Supplementary Figures**

Fig.S1. The synthesis of arthrocolins in twelve distinct *E. coli* strains fed with toluquinol on two media, LB and NB.

Fig. S2. A) The effects of indole and tryptophanase gene *tnaA* on the synthesis of arthrocolins in *E. coli*. T: toluquinol, I: indole. B) Comparison of arthrocolin contents in Δ*tnaA* treated with and without indole. C) Comparison of arthrocolin contents in WT treated with and without indole.

Fig S3. The effects of aromatic amino acids on the synthesis of arthrocolins A–C in *E. coli* OP50 fed with toluquinol.

Fig S4. The effects of Bezyl alcohol, benzaldehyde, and benzoic acid on the synthesis of arthrocolins A–C in *E. coli* OP50 fed with toluquinol.

Fig. S5. The levels of benzoic acid, hydroxybenzoic acid, indole and three aromatic amino acids in the in twelve *E. coli* strains fed with toluquinol. Phe: phenylalanine, Tyr: tyrosine, and Trp: tryptophan.

Fig S6. KEGG pathway analysis revealed that the 2591 genes whose deficiency caused the reductions in arthrocolin contents were mainly enriched in carbon metabolism, glycolysis/gluconeogenesis, starch and sucrose metabolism and pentose phosphate pathway.

Fig S7. Monose, together with glycerol, inhibited the synthesis of arthrocolins in *E. coli* fed with toluquinol.

Fig. S8. Glucose inhibited the synthesis of arthrocolins and degradation of aromatic amino acids in *E. coli* fed with toluquinol.

Fig. S9. The effect Glucose inhibited the synthesis of arthrocolins and degradation of aromatic amino acids in *E. coli* fed with toluquinol.

Fig. S10. Glucose increased the redox reduction potentials in *E. coli*.

Fig S11. Comparison of redox reduction potentials and the arthrocolin contents in ten mutants fed with toluquinol indicated that the redox reduction potentials were not related to the arthrocolin contents in *E. coli*..

Fig. S12. Comparison of the arthrocolins levels in the bacteria, cultural broths and supernates of *E. coli* fed with toluquinol under heat killing treatment.

Fig. S13. The arthrocolins levels were related to bacterial weights of *E. coli* fed with toluquinol under heat killing treatment.

Fig. S14. The effects of toluquinol on the levels of superoxide anions (A), lipid peroxidation (B), ROS (C) and lipid (D) in *E. coli*.

Fig. S15. The effects of toluquinol on the levels of ferrous, ferric and total iron and iron ratios in and out of six *E. coli* strains.

Fig. S16. A free radical capturing agent α-phenyl-tert-butylnitrone (PBN) inhibited the synthesis of arthrocolins in *E. coli* WT (A) fed with toluquinol and $\Delta$*tnaA.* (B) treated with indole and toluquinol.

Fig. S17. Glucose inhibited transformation of toluquinol to toluqunone.

Fig. S18. The $^{1}$H NMR spectrum of **10.**
Fig. S19. The $^{13}$C NMR spectrum of **10.**
Fig. S20. The HSQC spectrum of **10.**
Fig. S21. The $^{1}$H-$^{1}$HCOSY spectrum of **10.**
Fig. S22 The HMBC spectrum of **10.**

Fig. S23. The $^{1}$H NMR spectrum of **11 and 12.**
Fig. S24. The $^{13}$C NMR spectrum of **11 and 12.**
Fig. S25. The HSQC spectrum of **10**–**11 and 12.**
Fig. S26. The $^{1}$H-$^{1}$HCOSY spectrum of **11 and 12.**
Fig. S27. The HMBC spectrum of **11 and 12.**

Fig. S28. The $^{1}$H NMR spectrum of **13.**
Fig. S29.. The $^{13}$C NMR spectrum of **13.**
Fig. S30 The HSQC spectrum of **13.**
Fig. S31. The $^{1}$H-$^{1}$HCOSY spectrum of **13.**
Fig. S32. The HMBC spectrum of **13.**

Fig. S33. The $^{1}$H NMR spectrum of **14.**
Fig. S34. The $^{13}$C NMR spectrum of **14.**
Fig. S35. The HSQC spectrum of **14.**
Fig. S36. The $^{1}$H-$^{1}$HCOSY spectrum of **14.**
Fig. S37. The HMBC spectrum of **14.**

Fig. S38. The $^{1}$H NMR spectrum of **15.**
Fig. S39. The $^{13}$C NMR spectrum of **15.**
Fig. S40. The HSQC spectrum of **15.**
Fig. S41. The $^{1}$H-$^{1}$HCOSY spectrum of **15.**
Fig. S42. The HMBC spectrum of **15.**

Fig. S43. The $^{1}$H NMR spectrum of **16.**
Fig. S44. The $^{13}$C NMR spectrum of **16.**
Fig. S45. The HSQC spectrum of **16.**
Fig. S46. The $^{1}$H-$^{1}$HCOSY spectrum of **16.**
Fig. S47. The HMBC spectrum of **16.**

Fig. S48. The $^{1}$H NMR spectrum of **17.**
Fig. S49. The $^{13}$C NMR spectrum of **17.**
Fig. S50. The HSQC spectrum of **17.**
Fig. S51. The $^{1}$H-$^{1}$HCOSY spectrum of **17.**
Fig. S52. The HMBC spectrum of **17.**

Fig. S53. The $^{1}$H NMR spectrum of **20.**
Fig. S54. The $^{13}$C NMR spectrum of **20.**
Fig. S55. The HSQC spectrum of **20.**
Fig. S56. The $^{1}$H-$^{1}$HCOSY spectrum of **20.**
Fig. S57. The HMBC spectrum of **20.**

Fig. S58. The $^{1}$H NMR spectrum of **24.**
Fig. S59. The $^{13}$C NMR spectrum of **24.**
Fig. S60. The HSQC spectrum of **24.**
Fig. S61. The $^{1}$H-$^{1}$HCOSY spectrum of **24.**
Fig. S62. The HMBC spectrum of **24.**

Fig. S63. The $^{1}$H NMR spectrum of **26.**
Fig. S64. The $^{13}$C NMR spectrum of **26.**
Fig. S65. The HSQC spectrum of **26.**
Fig. S66. The $^{1}$H-$^{1}$HCOSY spectrum of **26.**
Fig. S67. The HMBC spectrum of **26.**

Fig. S68 The LC-MS analysis of *E. coli* fed with compounds **1-2** and **27-31**.
Fig. S69 The LC-MS analysis of *E. coli* fed with compounds **32-38**.
Fig. S70 The LC-MS analysis of *E. coli* fed with compounds **39-45**.
Fig. S71 The LC-MS analysis of *E. coli* fed with compounds **46-52**.
Fig. S72 The LC-MS analysis of *E. coli* fed with compounds **53-56**.

Fig. S73. The $^{1}$H NMR spectrum of **59.**
Fig. S74. The $^{13}$C NMR spectrum of **59.**
Fig. S75. The HSQC spectrum of **59.**
Fig. S76. The $^{1}$H-$^{1}$HCOSY spectrum of **59.**
Fig. S77. The HMBC spectrum of **59.**

Fig. S78. The $^{1}$H NMR spectrum of **63.**
Fig. S79. The $^{13}$C NMR spectrum of **63.**
Fig. S80. The HSQC spectrum of **63.**
Fig. S81 The $^{1}$H-$^{1}$HCOSY spectrum of **63.**
Fig. S82. The HMBC spectrum of **63.**

**1. Metabolites from *E. coli* strain $E_{BL21}$-*276* fed with toluquinol (1).**

**Compound 10:** $C_{29}H_{23}NO_4$; Negative HRESI-MS *m/z*: 432.1595 [M-H]$^-$, calculated as 432.1594 for $C_{29}H_{23}NO_4$; IR (KBr) $v_{max}$: 3427, 3057, 2926, 2859, 1613, 1509, 1488,1457, 1405, 1381, 1335, 1246, 1181, 1104, 1008, 932, 871, 787, 744, 648, 613, 581, 555, 528, 486, 456, 427 cm$^{-1}$; H NMR and $^{13}$C NMR, see Table S2. Crystal data for **10** (CCDC NO. 2349837)**:** $C_{29}H_{23}NO_4•2(CH_4O)$, $M$ = 513.57, $a$ = 9.0191(4) Å, $b$ = 20.9712(8) Å, $c$ = 13.7202(6) Å, $\alpha$ = 90°, $\beta$ = 96.485(2)°, $\gamma$ = 90°, $V$ = 2578.45(19) Å$^3$, $T$ = 100.(2) K, space group $P$121/$c$1, $Z$ = 4, $\mu$(Cu Kα) = 0.745 mm$^{-1}$, 74199 reflections measured, 5079 independent reflections ($R_{int}$ = 0.2421). The final $R_1$ values were 0.1083 ($I > 2\sigma(I)$). The final $wR(F^2)$ values were 0.3302 ($I > 2\sigma(I)$). The final $R_1$ values were 0.1357 (all data). The final $wR(F^2)$ values were 0.3469 (all data). The goodness of fit on $F^2$ was 1.319.

**Compound 11:** $C_{29}H_{23}NO_4$; $[\alpha]_D^{22.0}$ –4.45° ($c$ = 0.15, MeOH); Negative HRESI-MS *m/z*: 432.1595 [M-H]$^-$, calculated as 432.1594 for $C_{29}H_{23}NO_4$; IR (KBr) $v_{max}$: 3427, 3057, 2925, 2856, 1708, 1611, 1509, 1458,1411, 1380, 1334, 1244, 1178, 1149, 1101, 1036, 1010, 918, 863, 844, 817, 775, 750, 630, 614, 582, 521, 484, 427 cm$^{-1}$;$^1$H NMR and $^{13}$C NMR, see Table S2. Crystal data for **11** (CCDC NO. 2349838): $C_{29}H_{23}NO_4$, $M$ = 449.48, $a$ = 13.9126(10) Å, $b$ = 14.7966(11) Å, $c$ = 13.9592(14) Å, $\alpha$ = 90°, $\beta$ = 104.623(5)°, $\gamma$ = 90°, $V$ = 2780.5(4) Å$^3$, $T$ = 100.(2) K, space group $P$121/$c$1, $Z$ = 4, $\mu$(Cu Kα) = 0.577 mm$^{-1}$, 42430 reflections measured, 5284 independent reflections ($R_{int}$ = 0.3475). The final $R_1$ values were 0.1258 ($I > 2\sigma(I)$). The final $wR(F^2)$ values were 0.3260 ($I > 2\sigma(I)$). The final $R_1$ values were 0.2029 (all data). The final $wR(F^2)$ values were 0.3828 (all data). The goodness of fit on $F^2$ was 1.096.

**Compound 12:** $C_{29}H_{23}NO_4$; $[\alpha]_D^{22.0}$ 18.95° ($c$ = 0.12, MeOH); Negative HRESI-MS *m/z*: 432.1595 [M-H]$^-$, calculated as 432.1594 for $C_{29}H_{23}NO_4$. $^1$H NMR and $^{13}$C NMR, see Table S2.

**Compound 13**: $C_{21}H_{18}O_3$; UV (MeOH) $\lambda_{max}$ (log $\varepsilon$): 197 (4.05), 255 (3.63), 307 (3.58) nm; Negative HRESI-MS: *m/z*: 317.1180 [M-H]$^-$, calculated 317.1178 as for $C_{21}H_{17}O_3$ [M-H]$^-$. $^1$H NMR and $^{13}$C NMR, see Table S2.

**Compound 14**: $C_{25}H_{18}N_2O_3$; UV (MeOH) $\lambda_{max}$ (log $\varepsilon$): 195.5 (3.95), 218 (4.02), 280 (3.31) nm; Positive HRESI-MS: *m/z*: 417.1218 [M+Na]$^+$, calculated 417.1215 as for $C_{25}H_{18}N_2O_3Na$ [M+Na]$^+$. $^1$H NMR and $^{13}$C NMR, see Table S3.

**Compound 15**: $C_{25}H_{18}N_2O_3$; UV (MeOH) $\lambda_{max}$ (log $\varepsilon$): 195 (3.93), 217 (4.03), 281 (3.32) nm; Positive HRESI-MS: *m/z*: 417.1218 [M+Na]$^+$, calculated 417.1215 as for $C_{25}H_{18}N_2O_3Na$ [M+Na]$^+$. $^1$H NMR and $^{13}$C NMR, see Table S3.

**Compound 16:** colorless solid, UV (MeOH) $\lambda_{max}$ (log $\varepsilon$): 206 (4.06), 275 (3.89), 296.5 (3.88) nm; $C_{15}H_{13}NO_4$; $[\alpha]_D^{22.0}$ 6.17° ($c$ =0.12, MeOH); Positive HRESI-MS: *m/z*:

294.0746 [M+Na]$^+$, calculated as 294.0742 for $C_{15}H_{13}NO_4Na$. $^1$H NMR and $^{13}$C NMR, see Table S4.

**Compound 17:** colorless solid, $C_{15}H_{13}NO_2$; UV (MeOH) $\lambda_{max}$ (log $\varepsilon$): 204.5 (4.25), 246(3.97), 289.5 (3.98) nm; $C_{15}H_{13}NO_4$; $[\alpha]_D^{22.0}$ 10.22° (*c* =0.09, MeOH); Positive HRESI-MS: *m*/*z*: 262.0848 [M+Na]$^+$, calculated as 262.0844 for $C_{15}H_{13}NO_2Na$. $^1$H NMR and $^{13}$C NMR, see Table S4. Crystal data for **17** (CCDC NO. 23498379): $C_{15}H_{13}NO_2$, $M$ = 239.26, $a$ = 6.0106(3) Å, $b$ = 18.5798(9) Å, $c$ = 11.3684(5) Å, $\alpha$ = 90°, $\beta$ = 96.0690(10)°, $\gamma$ = 90°, $V$ = 1262.46(10) Å$^3$, $T$ = 102.(2) K, space group $P121/n1$, $Z$ = 4, $\mu$(Cu Kα) = 0.677 mm$^{-1}$, 19889 reflections measured, 2480 independent reflections ($R_{int}$ = 0.0696). The final $R_1$ values were 0.0477 ($I > 2\sigma(I)$). The final $wR(F^2)$ values were 0.1237 ($I > 2\sigma(I)$). The final $R_1$ values were 0.0496 (all data). The final $wR(F^2)$ values were 0.1259 (all data). The goodness of fit on $F^2$ was 1.046.

**Compound 18:** colorless solid, $C_{14}H_{11}NO$; Positive ESI-MS *m*/*z*: 232 [M+Na]$^+$.$^1$H NMR and $^{13}$C NMR, see Table S5.

**Compound 19:** colorless solid, $C_{23}H_{18}N_2O$; Positive ESI-MS *m*/*z*: 361 [M+Na]$^+$.$^1$H NMR and $^{13}$C NMR, see Table S5.

**Compound 20:** colorless solid, $C_{24}H_{19}N_3O_2$; UV (MeOH) $\lambda_{max}$ (log $\varepsilon$): 302 (3.83), 290 (3.93), 203 (4.82) nm; IR (KBr) $\nu_{max}$: 3440, 2925, 2854, 1629, 1562, 1494,1383, 1303, 1259, 1239, 1176, 1069, 1012, 979, 887, 848, 773, 699, 671, 519 cm$^{-1}$; Positive HRESI-MS *m*/*z*: 404.1379 [M+Na]$^+$, calculated as 404.1375 for $C_{24}H_{19}N_3O_2Na$. $^1$H NMR and $^{13}$C NMR, see Table S5.

**Compound 21:** colorless solid, $C_{26}H_{19}N_3O$; Positive ESI-MS *m*/*z*: 412 [M+Na]$^+$.$^1$H NMR and $^{13}$C NMR, see Table S6.

**Compound 22:** colorless solid, $C_{24}H_{17}N_3O$; Positive ESI-MS *m*/*z*: 386 [M+Na]$^+$.$^1$H NMR and $^{13}$C NMR, see Table S6.

**Compound 23:** colorless solid, $C_{24}H_{17}N_3O$; Positive ESI-MS *m*/*z*: 386 [M+Na]$^+$.$^1$H NMR and $^{13}$C NMR, see Table S6.

**Compound 24:** colorless solid, $C_{29}H_{18}N_2O_2$; $[\alpha]_D^{22.0}$ 34.15° (*c* = 0.15, MeOH);UV (MeOH) $\lambda_{max}$ (log $\varepsilon$): 196 (3.87), 223 (3.91), 282 (3.26) nm; Positive HRESI-MS *m*/*z*: 329.1269 [M+Na]$^+$, calculated as 329.1266 for $C_{29}H_{18}N_2O_2Na$. $^1$H NMR and $^{13}$C NMR, see Table S7;

**Compound 25:** colorless solid, $C_{22}H_{18}N_4$; Positive ESI-MS *m*/*z*: 361 [M+Na]$^+$.$^1$H NMR and $^{13}$C NMR, see Table S8.

**Compound 26:** colorless solid, $C_{30}H_{23}N_5$; UV (MeOH) $\lambda_{max}$ (log $\varepsilon$): 196 (3.83), 219 (3.93), 274 (3.28) nm; Positive HRESI-MS *m*/*z*: 476.1854 [M+Na]$^+$, calculated as 476.1851 for $C_{30}H_{23}N_5Na$. $^1$H NMR and $^{13}$C NMR, see Table S8.

**Table S2.** The $^{1}H$ NMR (500 MHz) and $^{13}C$ NMR (125 MHz) data of **10**–**13**.

| | 10 | 11 and 12 | 13 |
|---|---|---|---|
| 1 | 49.57 s | 49.84 s | 46.8 d, 5.03 s |
| Indole | | | |
| 2 | 127.98 d, 6.44 s | 128.03 d, 6.44 s | |
| 3 | 122.14 s | 122.02 s | |
| 4 | 127.39 s | 127.41 s | |
| 5 | 139.32 s | 139.30 s | |
| 6 | 112.09 d, 7.29 d (8.1) | 112.16 d, 7.32 d (7.4) | |
| 7 | 122.23 d, 6.97 t (8.1) | 122.23 d, 7.00 t (7.4) | |
| 8 | 119.35 d, 6.70 t (8.1) | 119.36 d, 6.73 overlap | |
| 9 | 123.11 d, 6.74 d (8.1) | 123.13 d, 6.75 overlap | |
| Phenyl | | | |
| 1' | 138.85 s | 138.70 s | 148.4 s |
| 2', 6' | 115.20 d, 6.60 d (8.2) | 115.21 d, 6.62 d (8.2) | 129.3 d, 7.14 d, 8.3 |
| 3', 5' | 130.64 d, 6.82 d (8.2) | 130.64 d, 6.83 d (8.2) | 129.7 d, 7.23 t, 8.3 |
| 4' | 156.50 s | 156.54 s | 127.6 d, 7.13, 8.2 |
| 5-substituted Toluquinol | | | |
| 1'' | 151.36 s | 151.44 s | |
| 2'' | 124.89 s | 124.93 s | |
| 3'' | 118.62 d, 6.83 s | 118.76 d, 6.91 s | |
| 4'' | 147.88 s | 147.95 s | |
| 5'' | 128.92 s | 128.94 s | |
| 6'' | 116.15 d, 6.37 s | 116.11 d, 6.37 s | |
| 7'' | 16.03 q, 2.17 s | 16.11 q, 2.21 s | |
| 6-substituted Toluquinol | | | |
| 1''' | | 146.56 s | 144.7s |
| 2''' | | 126.95 s | 127.8 s |
| 3''' | | 116.55 d, 6.57 d (3.0) | 117.3 d, 6.53 d, 2.8 |
| 4''' | | 152.45 s | 153.2 s |
| 5''' | | 114.39 d, 6.27 d (3.0) | 113.6 d, 6.30 d, 2.8 |
| 6''' | | 131.54 s | 124.7 s |
| 7''' | | 16.05 q, 2.35 s | 16.4 q, 2.34 s |

**Table S3.** The $^{1}H$ NMR (500 MHz) and $^{13}C$ NMR (125 MHz) data of **14**–**15**.

| No | **14** (EC-19) | **15 (EC-20)** |
|---|---|---|
| | $CD_3OD$ | $CD_3OD$ |
| 1 | 53.64 s | 53.31 s |
| 2 | 179.52 s | 179.84 s |
| 1' | 145.73 s | 153.61 s |
| 2' | 122.83 s | 126.80 s |
| 3', | 117.71 d, 6.67 s | 113.43 d, 7.06 s |
| 4' | 155.41 s | 147.06 s |
| 5' | 110.94 d, 6.60 s | 131.98 s |
| 6' | 134.55 s | 112.66 d, 6.77 s |
| 7' | 15.36 q, 2.39 s | 16.79 q, 2.27 s |
| 2'', 2''' | 125.91 d, 6.93 s | 125.88 d, 6.93 s |
| 3'', 3''' | 114.93 s | 115.09 s |
| 4'', 4''' | 121.80 d, 7.28 d (8.0) | 121.82 d, 7.30 d (8.0) |
| 5'', 5''' | 120.04 d, 6.88 t (8.0) | 120.02 d, 6.88 t (8.0) |
| 6'', 6''' | 122.83 d, 7.09 t (8.0) | 122.82 d, 7.09 t (8.0) |
| 7'', 7''' | 112.68 d, 7.34 d (8.0) | 112.66 d, 7.38 d (8.0) |
| 8'', 8''' | 126.94 s | 126.94 s |
| 9'', 9''' | 138.98 s | 138.97 s |

**Table S4.** The $^{1}H$ NMR (500 MHz) and $^{13}C$ NMR (125 MHz) data of **16**–**17**.

| No | **16 (ECB-30)** | **17 (ECB-25)** |
|---|---|---|
| | $CD_3OD$ | $CD_3OD$ |
| 1 | 79.78 d, 5.44 s | 165.65 s |
| 2 | 167.94 s | 36.71 t, 4.19 s 2H |
| 1' | 128.03 s | 136.69 s |
| 2', 6' | 130.02 d, 7.16 d 8.8, 2H | 130.07 d, 7.33 overlap, 2H |
| 3', 5' | 116.42 d, 6.73 d 8.8, 2H | 129.07 d, 7.33 overlap, 2H |
| 4' | 159.36 s | 128.41 d, 7.26 m |
| 1'' | 142.83 s | 151.53 s |
| 2'' | 119.90 s | 134.28 s |
| 3'' | 105.06 d, 6.35 s | 120.65 d, 7.30 s |
| 4'' | 153.19 s | 124.09 s |
| 5'' | 120.01 s | 155.41 s |
| 6'' | 118.61 d, 6.60 s | 97.53 d, 6.90 s |
| 7'' | 15.73 q, 2.10 s | 16.86 q, 2.24 s |

**Table S5.** The $^{1}H$ NMR (500 MHz) and $^{13}C$ NMR (125 MHz) data of **18**–**20**.

| No | **18** (EC-25) | **19** (EC-52) | **20** (EC-29/17) |
|---|---|---|---|
| | $CD_3OD$ | $CD_3OD$ | $CD_3OD$ |
| 1 | | 40.84 d | 57.41 s |
| | | 5.75 s | |
| 2 | | | 179.68 s |
| 1' | 130.71 s | 137.33 s | 135.32 s |
| 2',6' | 129.79 d, 6.98 d (2H, 8.7) | 130.73 d, 7.14 d (2H, 8.7) | 131.71 d, 7.23 d (2H, 9.0) |
| 3', 5' | 116.78 d, 6.60 d (2H, 8.7) | 115.73 d, 6.70 d (2H, 8.7) | 115.50 d, 6.68 d (2H, 9.0) |
| 4' | 156.71 s | 156.39 s | 157.05 s |
| 2'', 2''' | 131.92 d, 7.46 s | 124.65 d, 6.63 s | 126.68 d, 6.95 s |
| 3'', 3''' | 105.02 s | 120.61 s | 120.03 s |
| 4'', 4''' | 120.08 d, 7.47 d (7.8) | 120.53 d, 7.28 d (7.5) | 122.54 d, 7.06 d (8.0) |
| 5'', 5''' | 121.10 d, 7.04 t (7.8) | 119.27 d, 6.88 t (7.5) | 119.74 d, 6.80 t (8.0) |
| 6'', 6''' | 123.38 d, 7.15 t (7.8) | 122.09 d, 7.05 t (7.5) | 122.42 d, 7.04 t (8.0) |
| 7'', 7''' | 112.90 d, 7.41 d (7.8) | 112.05 d, 7.33 d (7.5) | 112.44 d, 7.34 d (8.0) |
| 8'', 8''' | 129.01 s | 128.45 s | 128.37 s |
| 9'', 9''' | 138.59 s | 138.60 s | 138.78 s |

**Table S6.** The $^{1}H$ NMR (500 MHz) and $^{13}C$ NMR (125 MHz) data of **21**–**23**.

| No | **21 (EC-32)** | **22 (EC-26)** | **23 (EC-18/35)** |
|---|---|---|---|
| | $CD_3OD$ | $CD_3OD$ | $CD_3OD$ |
| 1 | 45.26 d, 6.40 s | | |
| 1' | 198.13 s | | |
| 2' | 134.81 d, 8.38 s | 205.20 s | 182.90 s |
| 3' | 117.76 s | 69.93 s | 55.12 s |
| 4' | 128.08 d, 8,29 d (8.0) | 125.87 d, 7.61 d (8.0) | 126.62 d, 7.29 d (8.0) |
| 5' | 123.14 d, 7.17 t (8.0) | 118.92 d, 6.81 t (8.0) | 123.44 d, 7.00 t (8.0) |
| 6' | 124.23 d, 7.20 t (8.0) | 139.18 d, 7.55 t (8.0) | 129.16 d, 7.28 t (8.0) |
| 7' | 112.85 d, 7.42 d (8.0) | 113.35d, 7.01 d (8.0) | 111.13 d, 7.08 d (8.0) |
| 8' | 127.66 s | 119.74 s | 136.91 s |
| 9' | 138.42 s | 162.87 s | 142.46 s |
| 2'', 2''' | 125.11 d, 7.03 s | 125.38 d, 7.11 s | 125.74 d, 6.93 s |
| 3'', 3''' | 115.80 s | 115.35 s | 115.80 s |
| 4'', 4''' | 120.03 d, 7.57 d (8.0) | 121.76 d, 7.38 d (8.0) | 122.24 d, 7.30 d (8.0) |
| 5'', 5''' | 119.77 d, 6.95 t (8.0) | 119.79 d, 6.88 t (8.0) | 119.69 d, 6.84 t (8.0) |
| 6'', 6''' | 122.38 d, 7.07 t (8.0) | 122.49 d, 7.08 t (8.0) | 122.51 d, 7.06 t (8.0) |
| 7'', 7''' | 112.30 d, 7.33 d (8.0) | 121.76 d, 7.38 d (8.0) | 112.49 d, 7.35 t (8.0) |
| 8'', 8''' | 128.33 s | 127.21 | 127.44 s |
| 9'', 9''' | 138.41 s | 138.84 s | 138.93 s |

**Table S7.** The $^{1}H$ NMR (500 MHz) and $^{13}C$ NMR (125 MHz) data of **24**.

| No | **24(EC-39)** | | | | |
|---|---|---|---|---|---|
| | $CD_3OD$ | | | | |
| 1 | 38.13 d | 4.70 d 6.7 | | | |
| 2 | 76.39 d | 4.52 m | | | |
| 3 | 66.44 t | 3.65 dd 4.8, 11.0 | | | |
| | | 3.53 dd 7.4, 11.0 | | | |
| 2' | 123.90 d | 7.17 s | 2'' | 124.11 d | 7.31 s |
| 3' | 118.02 s | | 3'' | 118.02 s | |
| 4' | 120.22 d | 7.58 d (8.0) | 4'' | 120.03 d | 7.57 d (8.0) |
| 5' | 119.43 d | 6.96 t (8.0) | 5'' | 119.37 d | 6.94 t (8.0) |
| 6' | 122.15 d | 7.04 t (8.0) | 6' | 122.05 d | 7.06 t (8.0) |
| 7' | 112.16 d | 7.32 d (8.0) | 7'' | 112.03 d | 7.33 d (8.0) |
| 8' | 128.46 s | | 8'' | 129.31 s | |
| 9' | 138.00 s | | 9'' | 138.11 s | |

**Table S8.** The $^{1}H$ NMR (500 MHz) and $^{13}C$ NMR (125 MHz) data of **25**–**26**.

| No | **26 (EC-6)** | | | 25 (EC-40) | |
|---|---|---|---|---|---|
| 1 | 41.98 d | 6.02 s | | | |
| 2' | 158.41 s | | 2, 5 | 153.75 s | |
| 3' | 144.74 d | 8.46 s | 3, 6 | 142.98 d | 8.45 s, , 2H |
| 5' | 156.09 s | | | | |
| 6’ | 144.62 d | 8.41 s | | | |
| 7’ | 32.43 t | 4.29 s, 2H | 7 | 30.84 t | 4.17 s, 2H |
| 2'', 2''' | 125.02 d | 6.77 s, 2H | 2' | 123.43 d | 7.20 s, 2H |
| 3'', 3''' | 117.45 s | | 3' | 117.45 s | |
| 4'', 4''' | 120.08 d | 7.25 d 8.0, 2H | 4' | 118.40 d | 7.45 d 7.7, 2H |
| 5'', 5''' | 119.97 d | 6.89 t 8.0, 2H | 5' | 118.40 d | 6.94 t 7.7, 2H |
| 6'', 6''' | 122.70 d | 7.10 t 8.0, 2H | 6' | 121.00 d | 7.05 t 7.7, 2H |
| 7'', 7''' | 112.54 d | 7.35 d 8.0, 2H | 7' | 111.40 d | 7.33 d 7.7, 2H |
| 8'', 8''' | 128.15 s | | 8' | 126.81 s | |
| 9'', 9''' | 138.61 s | | 9' | 136.23 s | |
| 2'''' | 124.49 d | 7.11 s | | | |
| 3''''' | 112.76 s | | | | |
| 4'''' | 119.38 d | 7.37 d 8.0 | | | |
| 5'''' | 120.08 d | 6.95 t 8.0 | | | |
| 6'''' | 122.72 d | 7.09 t 8.0 | | | |
| 7'''' | 112.54 d | 7.34 d 8.0 | | | |
| 8'''' | 128.50 s | | | | |
| 9'''' | 138.47 s | | | | |

### 2 Metabolites from *E. coli* strain $E_{BL21}$-*276* fed with resorcinol (27).

**57:** colorless solid, $C_{13}H_{12}O_4$; $^{1}H$ NMR and $^{13}C$ NMR, see Table S9; Negative HRESI-MS *m/z*: 231.0649 $[M-H]^-$, calculated as 231.0657 for $C_{13}H_{11}O_4$.

**58:** colorless solid, $C_{13}H_{10}O_4$; $^{1}H$ NMR and $^{13}C$ NMR, see Table S9; Negative HRESI-MS *m/z*: 229.0495 $[M-H]^-$, calculated as 229.0501 for $C_{13}H_9O_4$.

**59:** colorless solid, $C_{15}H_{11}NO_3$; UV (MeOH) $\lambda_{max}$ (log $\varepsilon$): 302 (3.83), 290 (3.93), 203 (4.82) nm; IR (KBr) $\nu_{max}$: 3440, 2925, 2854, 1629, 1562, 1494,1383, 1303, 1259, 1239, 1176, 1069, 1012, 979, 887, 848, 773, 699, 671, 519 $cm^{-1}$; $^{1}H$ NMR and $^{13}C$ NMR, see Table S9; Negative HRESI-MS *m/z*: 252.0653 $[M-H]^-$, calculated as 252.0661 for $C_{15}H_{11}NO_3$.

Table S9. The [1]H and [13]C NMR spectroscopic data of metabolites **57**-**59** with data measured in acetone-$d_6$ at 500 MHz for $^1H$ NMR and 125 MHz for $^{13}C$ NMR with reference to the $CD_3OD$ solvent signals, $\delta$ in ppm and $J$ in Hz.

| | **57** (R-2) | **58** (R-5) | **59** (R-4) |
|---|---|---|---|
| 1 | 29.76 t, 3.67 s (2H) | 200.42 s | 194.92 s |
| 1' | 156.47 s | 167.11 s | 166.09 s |
| 2' | 103.22 d, 6.28 d (2.4, 2H) | 104.06 d, 6.34 d (2.2) | 104.10 d, 6.35 d (2.4) |
| 3' | 157.66 s | 166.33 s | 165.32 s |
| 4' | 107.97 d, 6.21 dd (2.4, 8.2, 2H) | 108.84 d, 6.35 dd (2.2, 9.0) | 108.60 d, 6.40 dd (2.4, 8.8) |
| 5' | 131.79 d, 6.83 d (8.2, 2H) | 126.62 d, 7.50 (9.0) | 135.26 d, 7.83 d (8.8) |
| 6' | 120.47 s | | 115.50 s |
| 1" | | 131.09 s | |
| 2" | | 132.68 d, 7.56 d (8.3, 2H) | 134.31 d, 7.89 (s) |
| 3" | | 116.06 s, 6.89 d (8.3, 2H) | 116.55 s |
| 4" | | 162.52 s | 122.87 d, 8.12 d (8.0) |
| 5" | | | 122.94 d, 7.21 t (8.0) |
| 6" | | | 124.48 d, 7.25 t (8.0) |
| 7" | | | 112.98 d, 7.49 d (8.0) |
| 8" | | | 128.10 s |
| 9" | | | 138.34 s |

**3 Metabolites from *E. coli* strain $E_{BL21}$-*276* fed with 2-chloro-benzene-1,4-diol (31).**

**60(Cl-4):** colorless solid, $C_{20}H_{14}O_6$; UV (MeOH) $\lambda_{max}$ (log $\varepsilon$): 302 (3.83), 290 (3.93), 203 (4.82) nm; IR (KBr) $v_{max}$: 3440, 2925, 2854, 1629, 1562, 1494,1383, 1303, 1259, 1239, 1176, 1069, 1012, 979, 887, 848, 773, 699, 671, 519 $cm^{-1}$; Positive HRESI-MS $m/z$: 373.06830 $[M+Na]^+$; Negative HRESI-MS $m/z$: 349.07129 $[M-H]^-$, calculated as 439.0712 for $C_{20}H_{14}O_6$. $^1H$ NMR and $^{13}C$ NMR, see Table S10;

**4 Metabolites from *E. coli* strain $E_{BL21}$-*276* fed with 9*H*-xanthen-9-ol (56).**

**61** Pale yellow solid, $C_{13}H_8O_2$.

**62:** $C_{21}H_{15}NO$; Pale purple solid; Negative HRESI-MS $m/z$: 296.10752 $[M-H]^-$, calculated as 296.1075 for $C_{21}H_{14}NO$.

**63:** $C_{29}H_{20}N_2O$; Pale purple solid; UV (MeOH) $\lambda_{max}$ (log $\varepsilon$): 302 (3.83), 290 (3.93), 203 (4.82) nm; IR (KBr) $v_{max}$: 3416, 3129,3054, 1617, 1597, 1472,1444, 1238, 1213, 1151, 1097, 1013, 963, 870, 795, 763, 742, 605 $cm^{-1}$; Negative HRESI-MS $m/z$: 411.15065 $[M-H]^-$, calculated as 411.1497 for $C_{29}H_{20}N_2O$. $^1H$ NMR and $^{13}C$ NMR, see Table S10.

Table S10. The $^{1}H$ and $^{13}C$ NMR spectroscopic data of metabolites **60** and **63** with data measured in acetone-$d_6$ at 500 MHz for $^{1}H$ NMR and 125 MHz for $^{13}C$ NMR with reference to the solvent signals, $\delta$ in ppm and $J$ in Hz.

| **60**(Cl-4) | | **60** (Cl-4) | | **63** | |
|---|---|---|---|---|---|
| $CD_3OD$ | | DMSO-$d_6$ | | $CD_3OD$ | |
| 1 | 57.99 s | 1 | 55.72 s | 1 | 44.35 s |
| 2 | 181.22 s | 2 | 178.06 s | | |
| 1' | 130.41 s | 1' | 127.83 s | 1' | 125.69 s |
| 2',6' | 131.55 d, 7.23 d | 2',6' | 129.84 d, 7.11 d (2H, 8.6) | 2' | 129.32 d, 7.18 d (8.8) |
| 3',5' | 116.14 d, 6.77 d | 3',5' | 115.15 d, 6.76 d (2H, 8.6) | 3' | 122.57 d, 6.97 t (8.8) |
| 4' | 159.36 s | 4' | 157.82 s | 4' | 127.54 d, 7.26 t (8.8) |
| 1'' | 122.93 s | 1'' | 121.13 s | 5' | 116.02 d, 7.20 d (8.8) |
| 2'' | 158.52 s | 2'' | 153.62 s | 6' | 152.62 s |
| 3'' | | 3'' | 97.86 d<br>6.59 s | 2'' | 125.90 d, 6.63 s |
| 4'' | 159.36 s | 4'' | 157.82 s | 3'' | 119.79 s |
| 5'' | | 5'' | 110.53 d, 6.55 d (8.0) | 4'' | 121.45 d, 7.13 d (8.0) |
| 6'' | | 6'' | 126.07 d, 6.80 d (8.0) | 5'' | 118.41 d, 6.80 t (8.0) |
| 1''' | | 1''' | 120.27 s | 6'' | 121.10 d, 7.05 t (8.0) |
| 2''' | | 2''' | 155.15 s | 7'' | 111.50 d, 7.42 d (8.0) |
| 3''' | | 3''' | 102.42 s<br>6.16 s | 8'' | 129.81 s |
| 4''' | | 4''' | 157.04 s | 9'' | 137.70 s |
| 5''' | | 5''' | 105.79 d, 6.12 d (8.0) | | |
| 6''' | | 6''' | 129.50 d, 6.40 d (8.0) | | |

**Table S11.** The effects of fifteen compounds, **10**–**15** and **18**-**26**, on four cancer cell lines, including human breast carcinoma cell MDA-MD-231, human colorectal carcinoma cells HCT116, human hepatocarcinoma cells SMMC-7721 and glioma cells U251.

| NO | **10** | **11** | **12** | **13** | **14** | **15** | **18** | **19** | **20** | **21** | **22** | **23** | **24** | **25** | **26** |
|---|---|---|---|---|---|---|---|---|---|---|---|---|---|---|---|
| MDA-MD-231 | - | - | - | - | - | 6.8 | - | 8.4 | - | 8.8 | 8.9 | 7.2 | - | - | - |
| U251 | - | - | - | - | - | 8.3 | - | 8.6 | - | 8.8 | 9.4 | 8.1 | - | - | - |
| HCT116 | - | - | - | - | - | 8.2 | - | 8.4 | - | 7.6 | - | 7.2 | - | - | - |
| SMMC-7721 | - | - | - | - | - | 8.9 | 8.8 | 8.4 | - | 7.8 | 9.0 | 9.6 | - | - | - |

**Table 12.** The $IC_{50}$ (μM) values of compounds **62**–**63** against ten cancer cells.

| No | 786-0 | A375 | A549 | CEM-SS | HCT116 | HOS | MDA-MB-231 | SGC-7901 | SMCC-7721 | U87 |
|---|---|---|---|---|---|---|---|---|---|---|
| **60** | - | - | - | - | - | - | - | - | - | - |
| **20** | - | - | - | - | - | - | - | - | - | - |
| **62** | 9.1 | 4.8 | 2.2 | 6.8 | 5.4 | 5.3 | 7.9 | 6.1 | 5.7 | 5.7 |
| **63** | - | 6.8 | 9.8 | - | 6.0 | 3.8 | 9.8 | 6.4 | 9.7 | - |

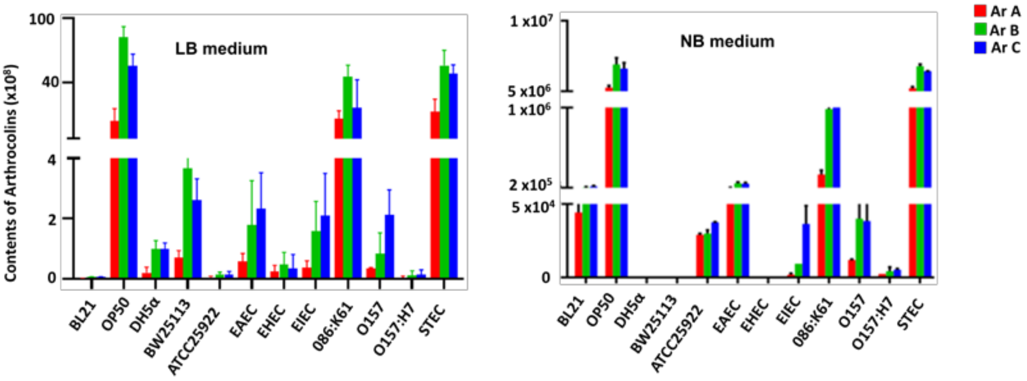


Figure S1. The synthesis of arthrocolins in twelve distinct *E. coli* strains fed with toluquinol on two media, LB and NB. Ar: arthrocolin.

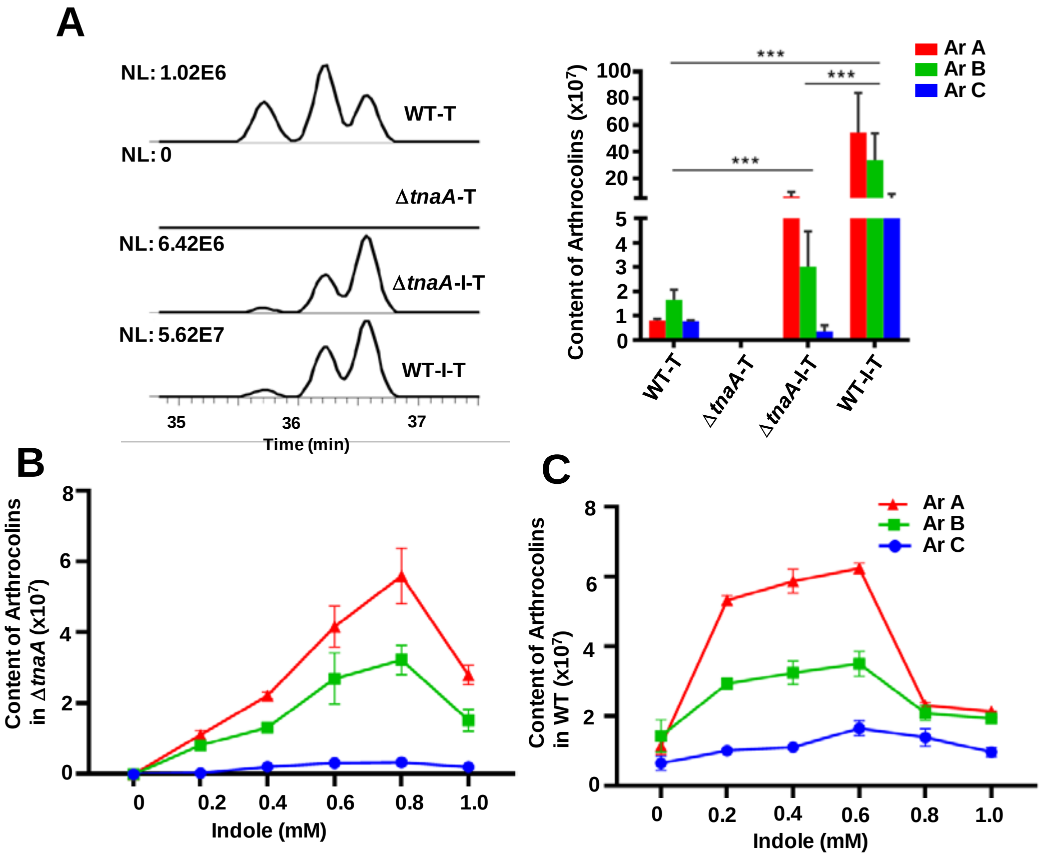


Figure S2. A) The effects of indole and tryptophanase gene *tnaA* on the synthesis of arthrocolins in *E. coli*. T: toluquinol, I: indole. B) Comparison of arthrocolin contents in Δ*tnaA* treated with and without indole. C) Comparison of arthrocolin contents in WT treated with and without indole. Ar: arthrocolins.

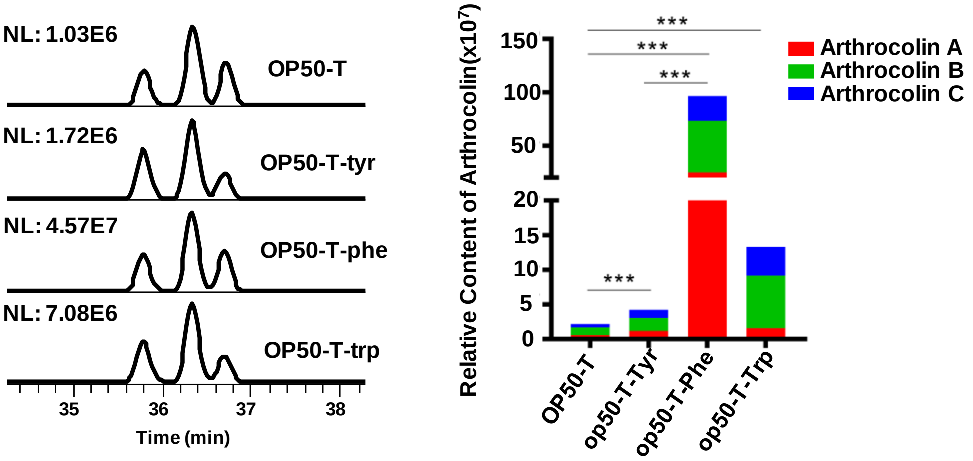


Figure S3. The effects of aromatic amino acids on the synthesis of arthrocolins A–C in *E. coli* OP50 fed with toluquinol. T: toluquinol, Con: solvent, Phe: phenylalanine, Tyr: tyrosine, and Trp: tryptophan.

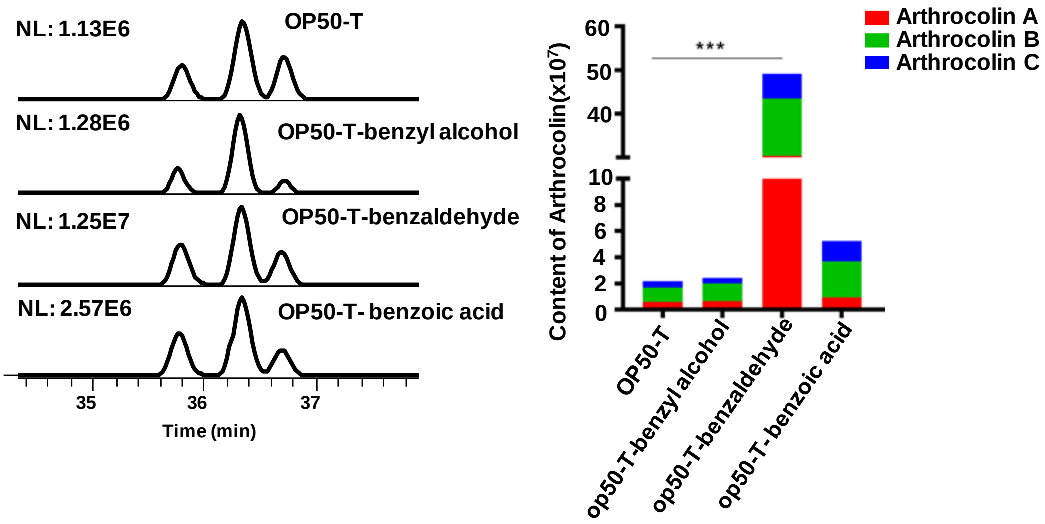


Figure S4. The effects of bezyl alcohol, benzaldehyde, and benzoic acid on the synthesis of arthrocolins A–C in *E. coli* OP50 fed with toluquinol. T: toluquinol.

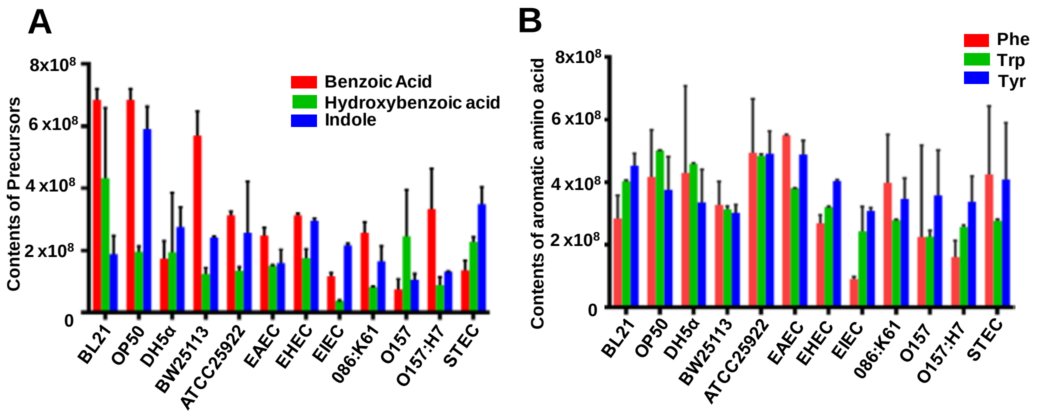


Figure S5. The levels of benzoic acid, hydroxybenzoic acid, indole and three aromatic amino acids in the in twelve *E. coli* strains fed with toluquinol. Phe: phenylalanine, Tyr: tyrosine, and Trp: tryptophan.

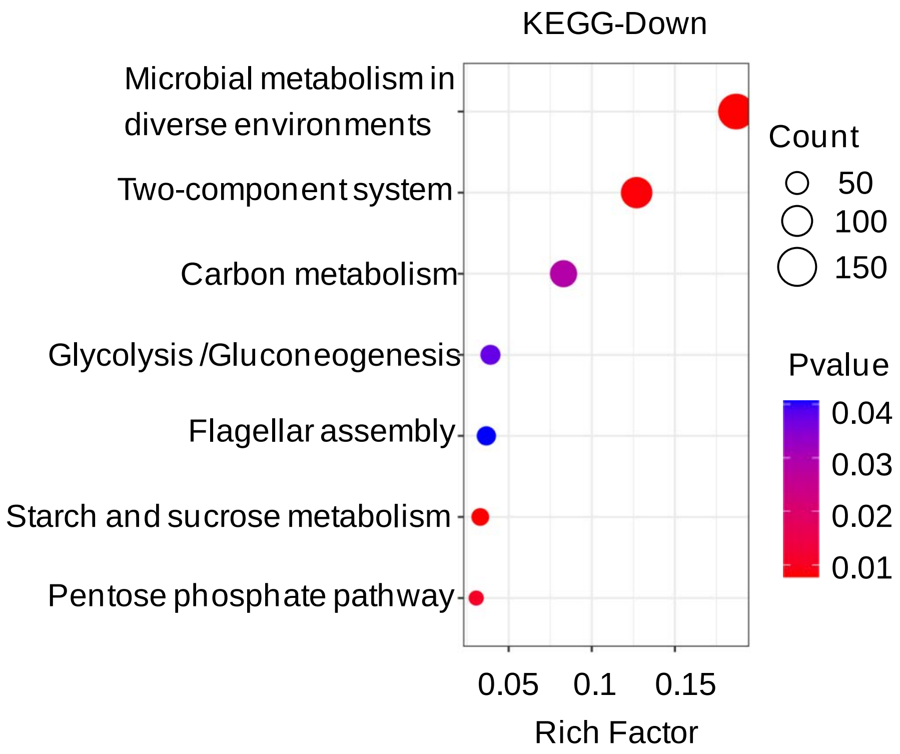


Figure S6. KEGG pathway analysis revealed that the 2591 genes whose deficiency caused the reductions in arthrocolin contents were mainly enriched in carbon metabolism, glycolysis/gluconeogenesis, starch and sucrose metabolism and pentose phosphate pathway.

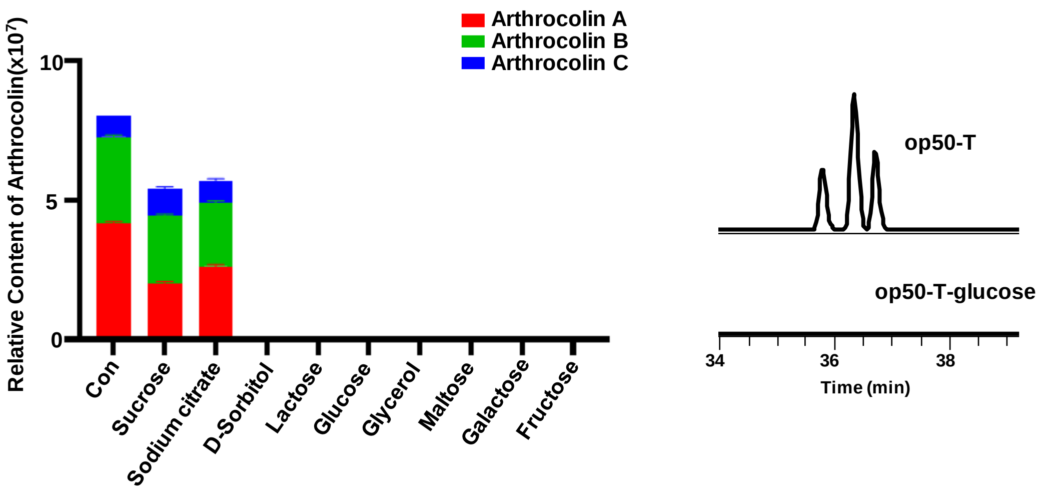


Figure S7. Monose, together with glycerol, inhibited the synthesis of arthrocolins in *E. coli* fed with toluquinol. T: toluquinol.

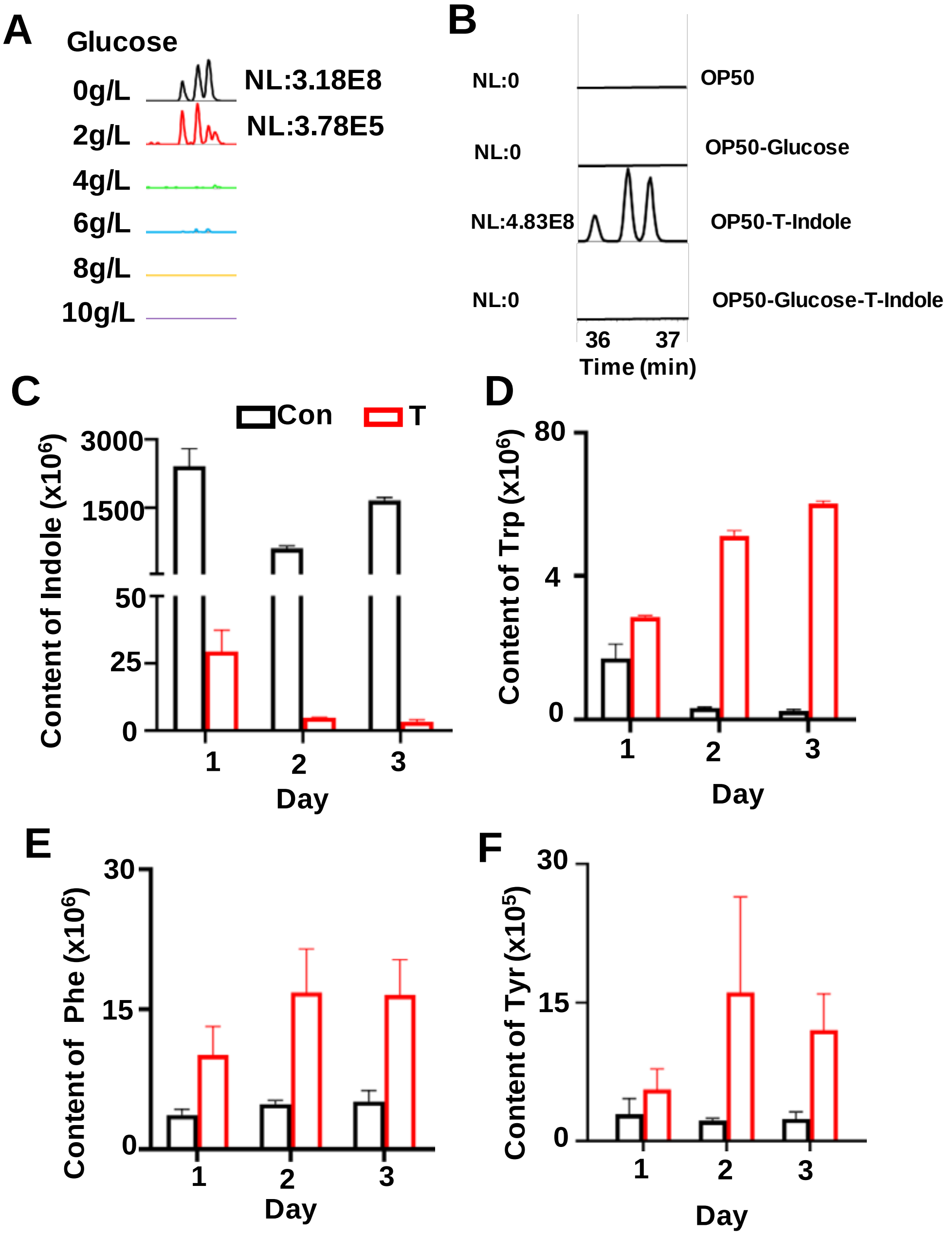


Figure S8. Glucose inhibited the synthesis of arthrocolins and degradation of aromatic amino acids in *E. coli* fed with toluquinol. T: toluquinol, Con: solvent, Trp: tryptophan, Phe: phenylalanine, and Tyr: tyrosine.

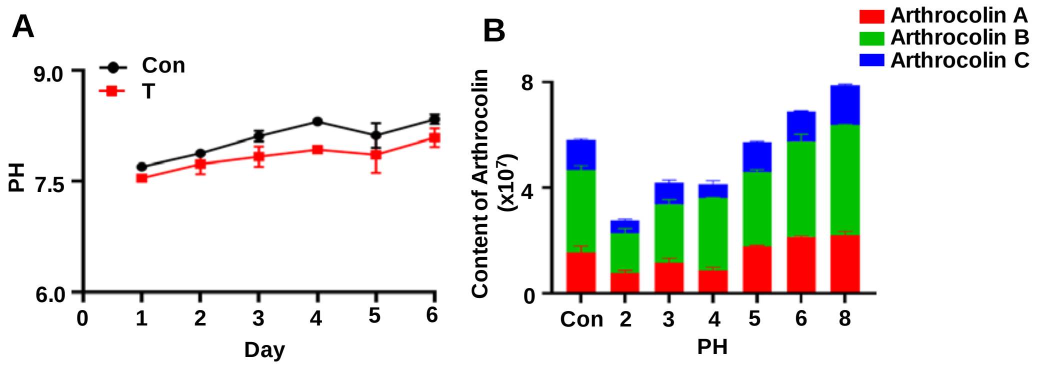


Figure S9. The effect Glucose inhibited the synthesis of arthrocolins and degradation of aromatic amino acids in *E. coli* fed with toluquinol. T: toluquinol, Con: solvent, Phe: phenylalanine, Tyr: tyrosine, and Trp: tryptophan.

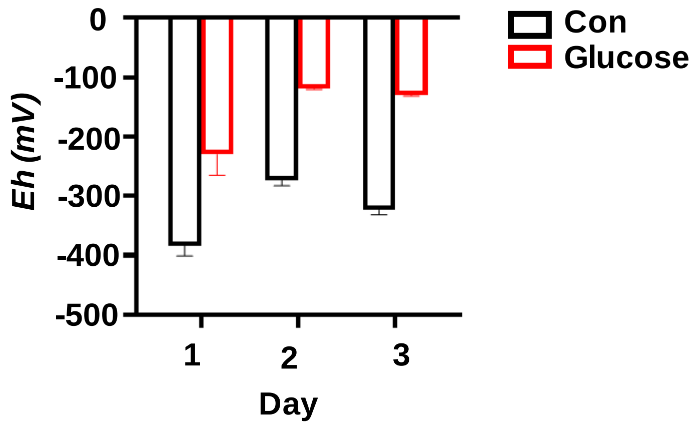


Figure S10. Glucose increased the redox reduction potentials in *E. coli*. Con: solvent.

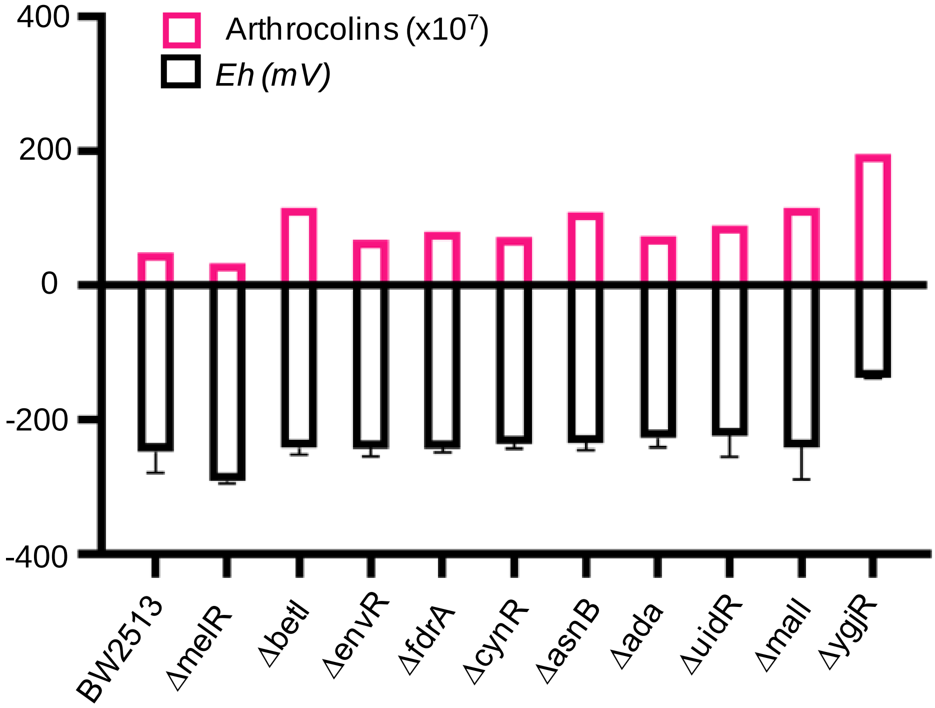


Figure S11. Comparison of redox reduction potentials and the arthrocolin contents in ten mutants fed with toluquinol indicated that the redox reduction potentials were not related to the arthrocolin contents in *E. coli*.

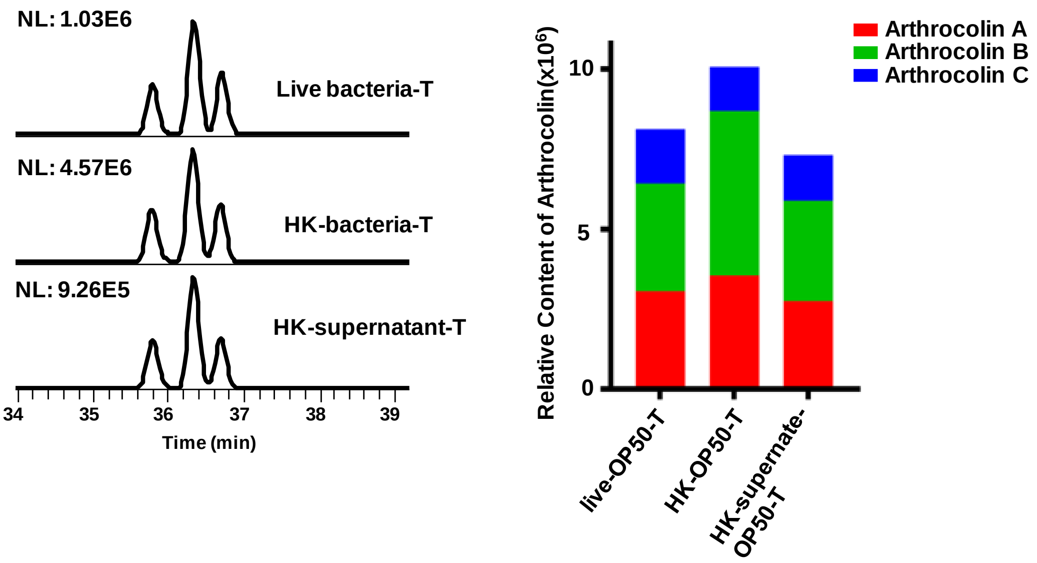


Figure S12. Comparison of the arthrocolins levels in the bacteria, cultural broths and supernates of *E. coli* fed with toluquinol under heat killing treatment. HK: Heat killing treatment.

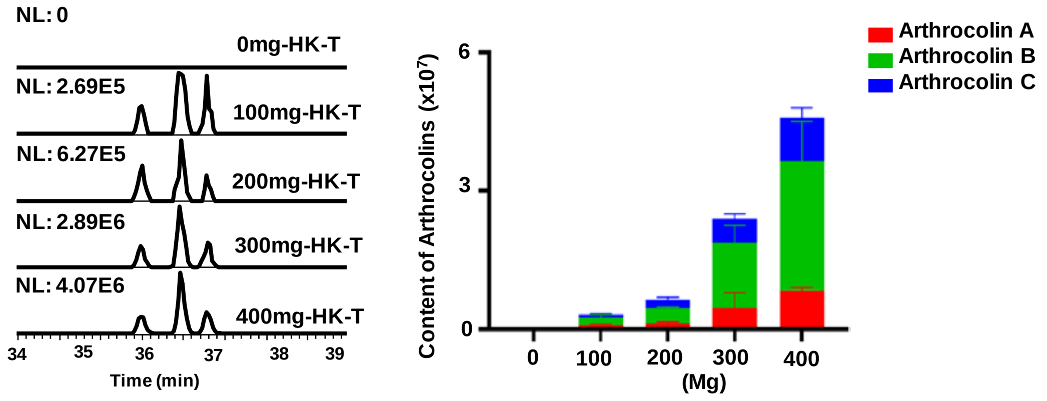


Figure S13. The arthrocolins levels were related to bacterial weights of *E. coli* fed with toluquinol under heat killing treatment. HK: heat killing treatment.

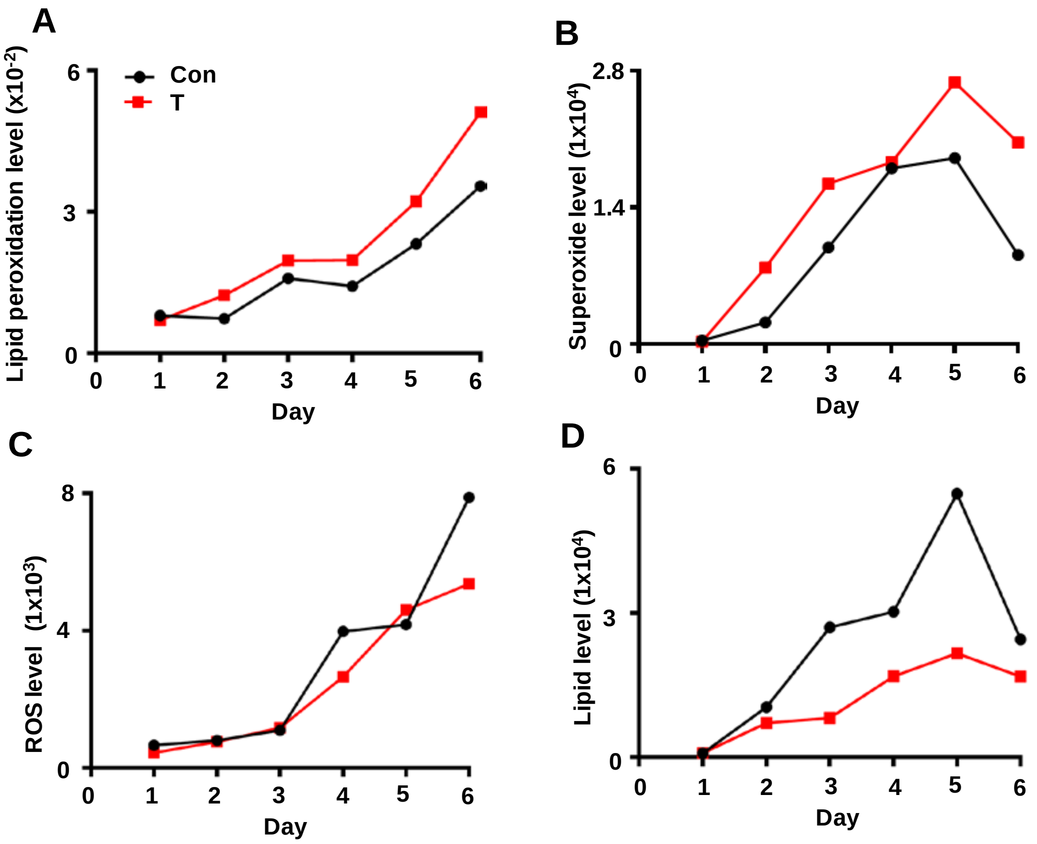


Figure S14. The effects of toluquinol on the levels of superoxide anions (A), lipid peroxidation (B), ROS (C) and lipid (D) in *E. coli*. T: Toluquinol.

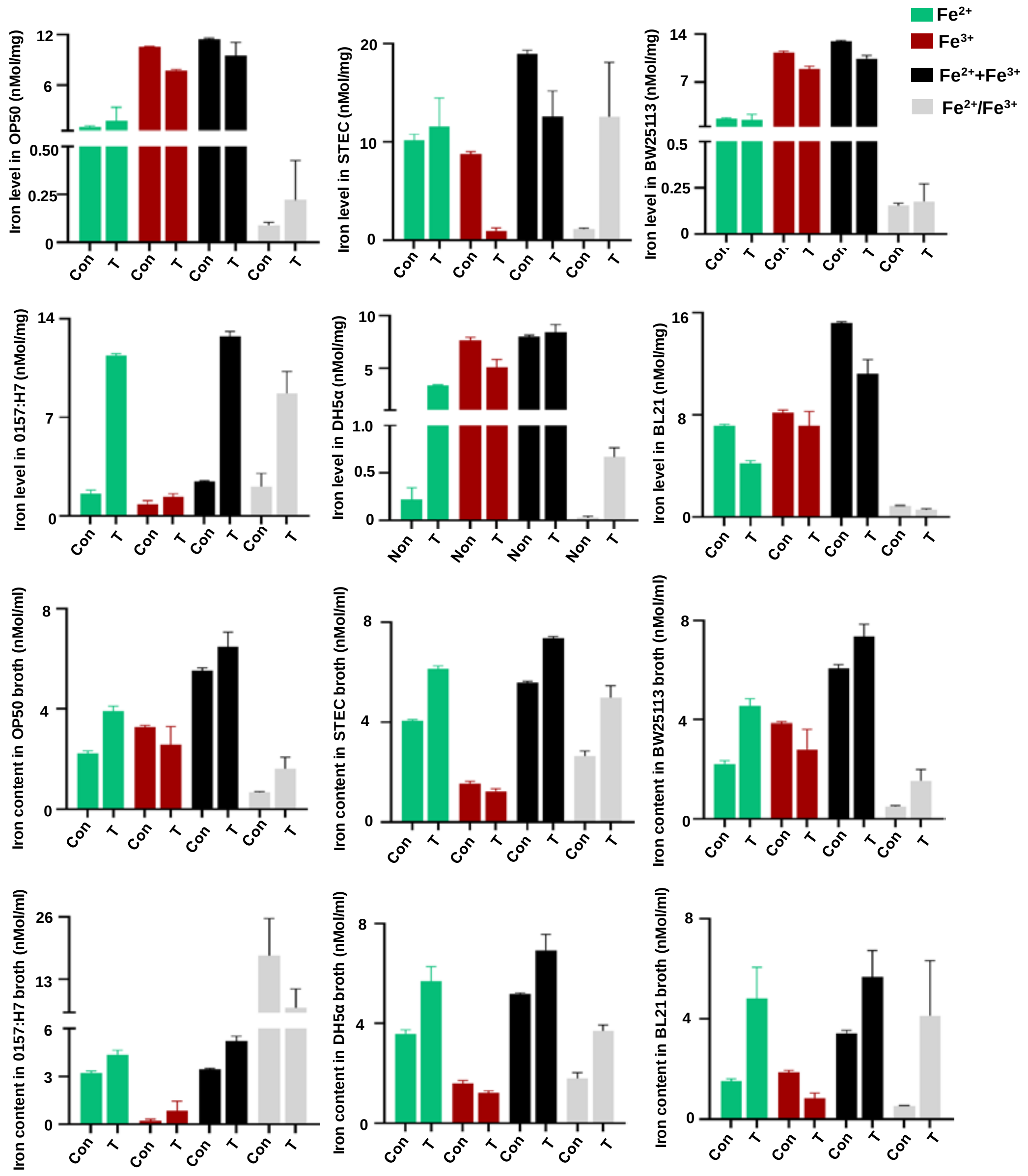


Figure S15. The effects of toluquinol on the levels of ferrous, ferric and total iron and iron ratios in and out of six *E. coli* strains. T: Toluquinol. Con: Solvent.

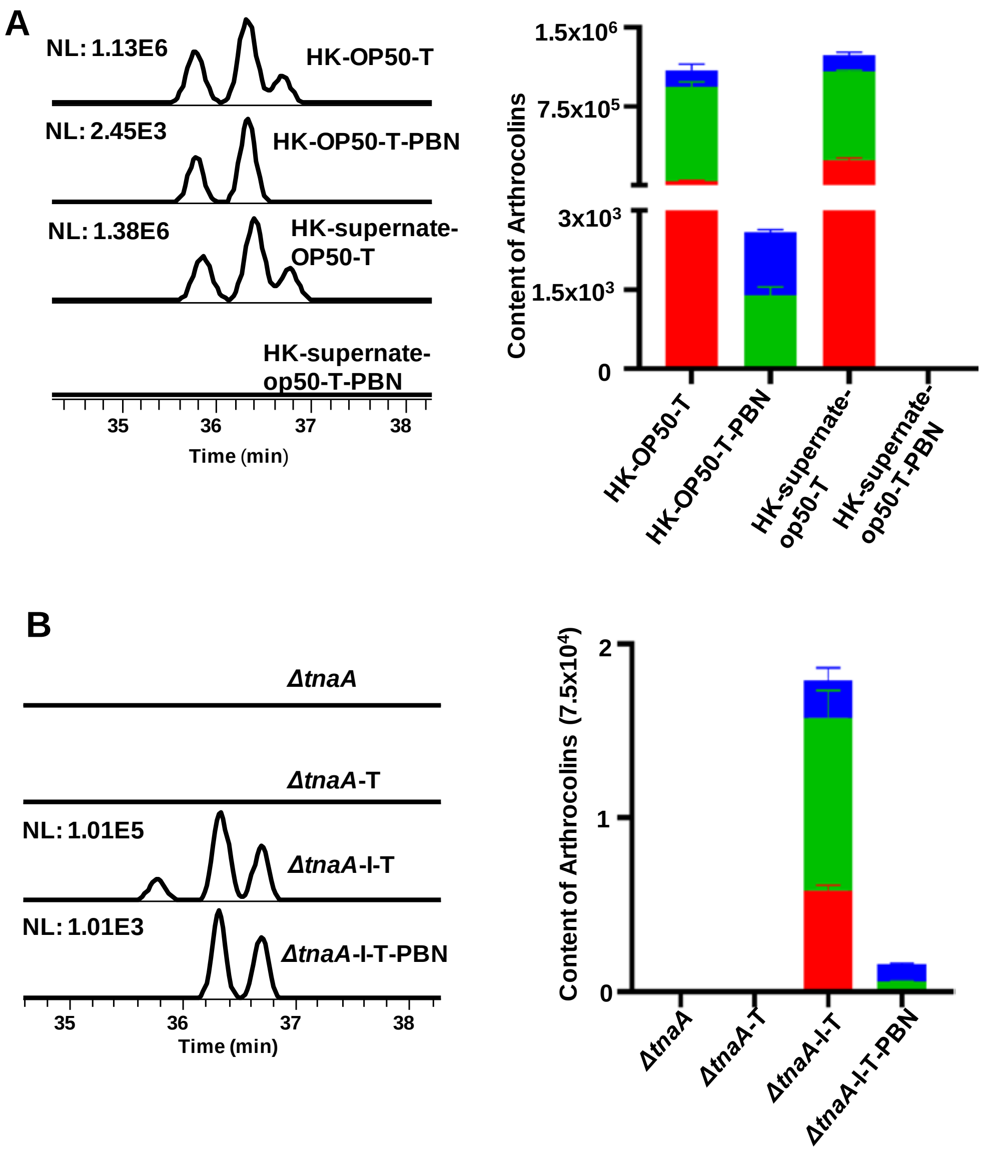


Figure S16. A free radical capturing agent α-phenyl-tert-butylnitrone (PBN) inhibited the synthesis of arthrocolins in *E. coli* WT (A) fed with toluquinol and Δ*tnaA* (B) treated with indole and toluquinol. T: Toluquinol, I: indole.

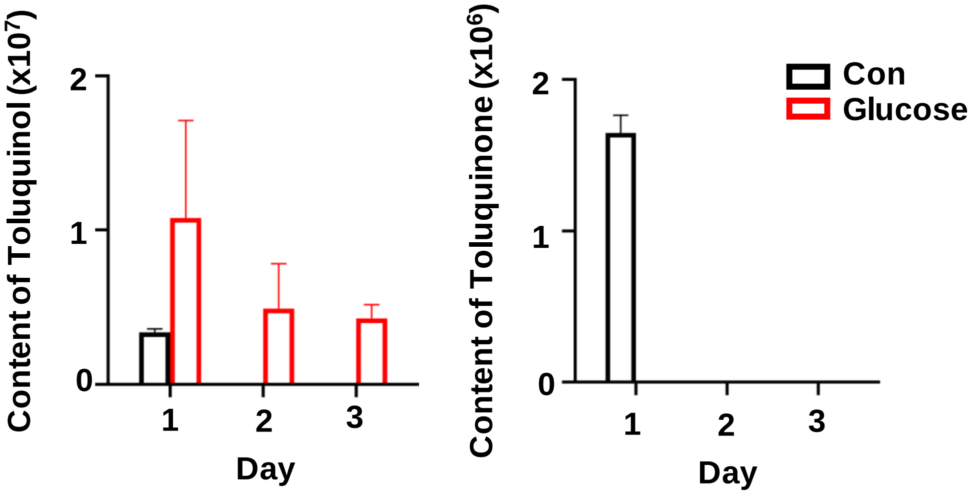


Figure S17. Glucose inhibited transformation of toluquinol to toluqunone. (Left) Comparison of toluquinol levels in *E. coli* fed with glucose and toluquinol. (Right) Comparison of toluquinone levels in *E. coli* fed with glucose and toluquinol. Con: solvent.

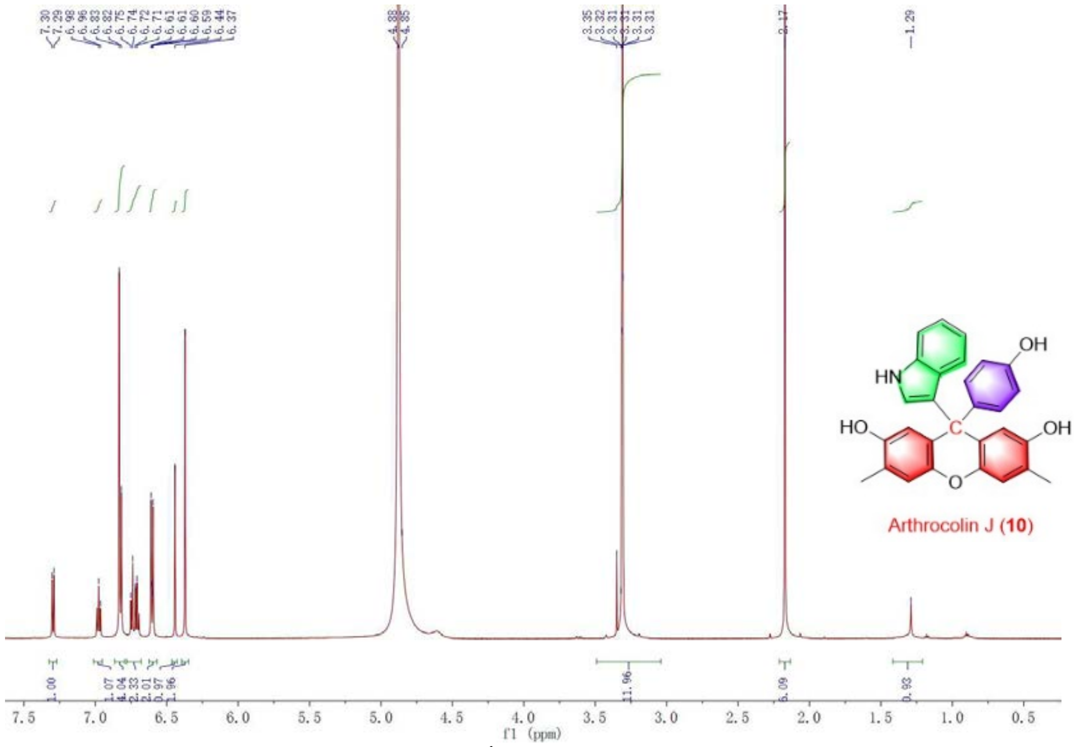


Figure S18. The $^{1}H$ NMR spectrum of **10.**

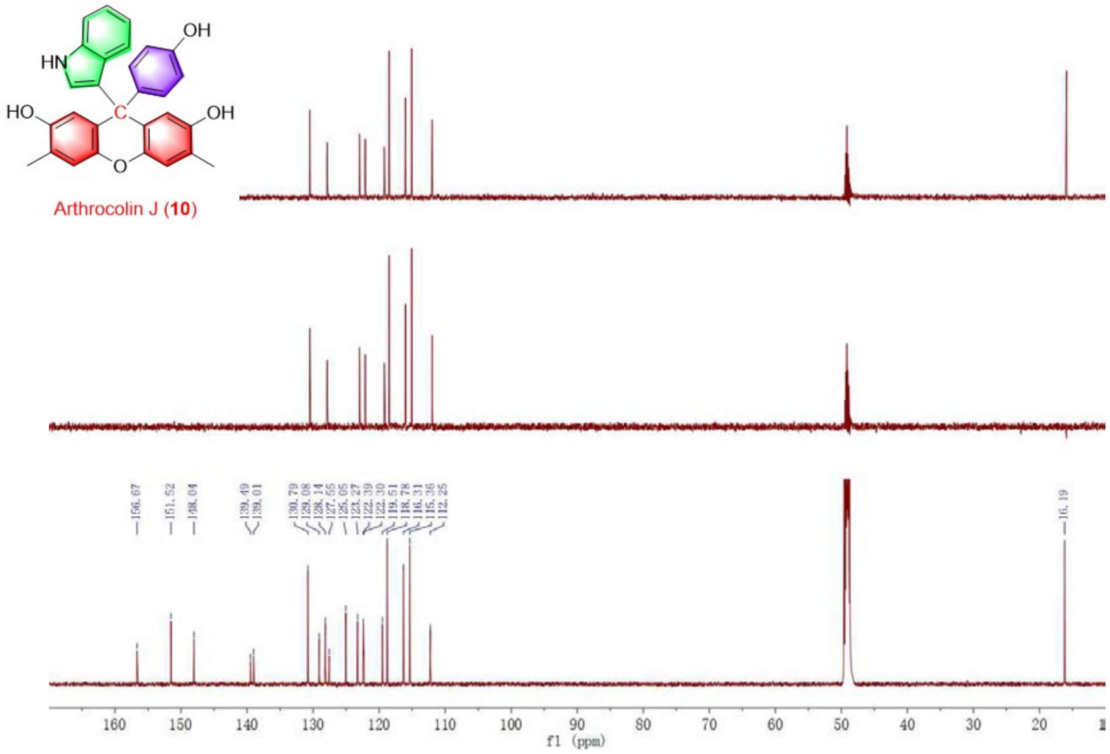


Figure S19. The $^{13}C$ NMR spectrum of **10.**

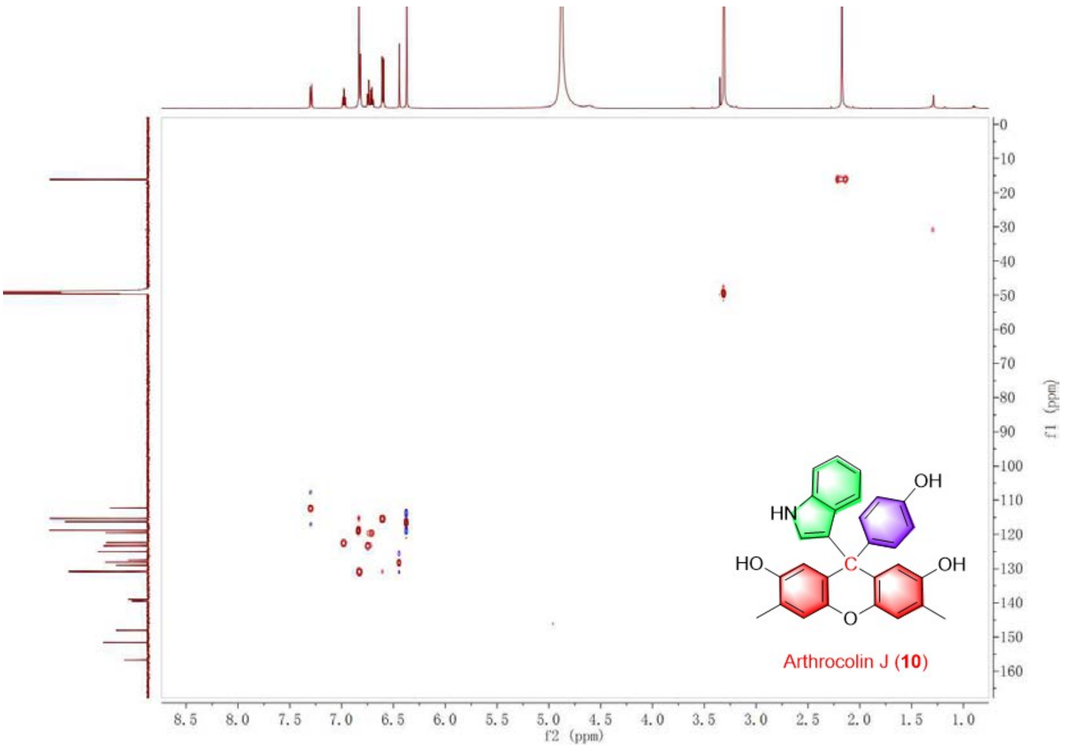


Figure S20. The HSQC spectrum of **10.**

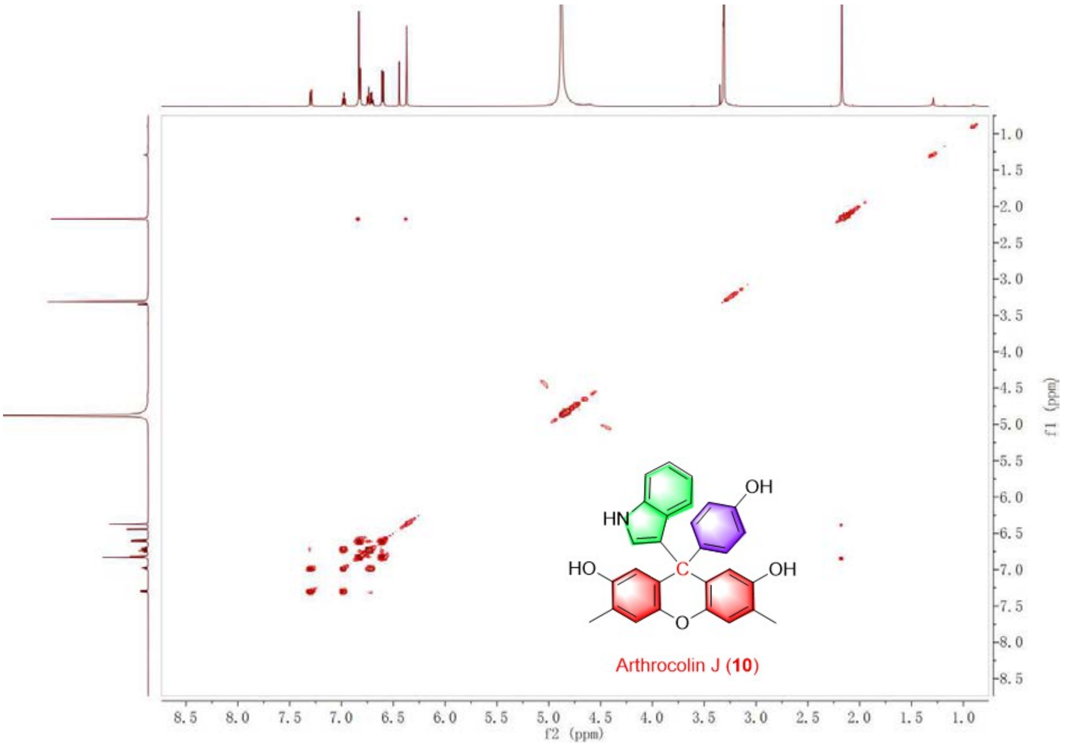


Figure S21. The $^{1}H$-$^{1}H$COSY spectrum of **10.**

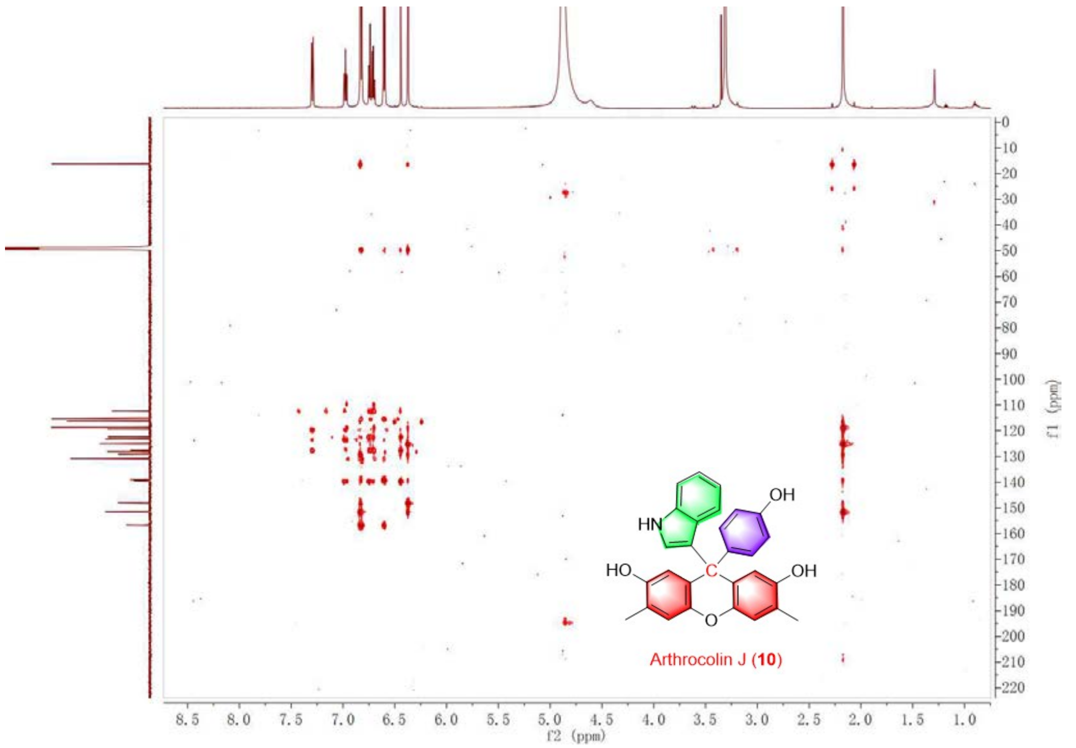


Figure S22. The HMBC spectrum of **10.**

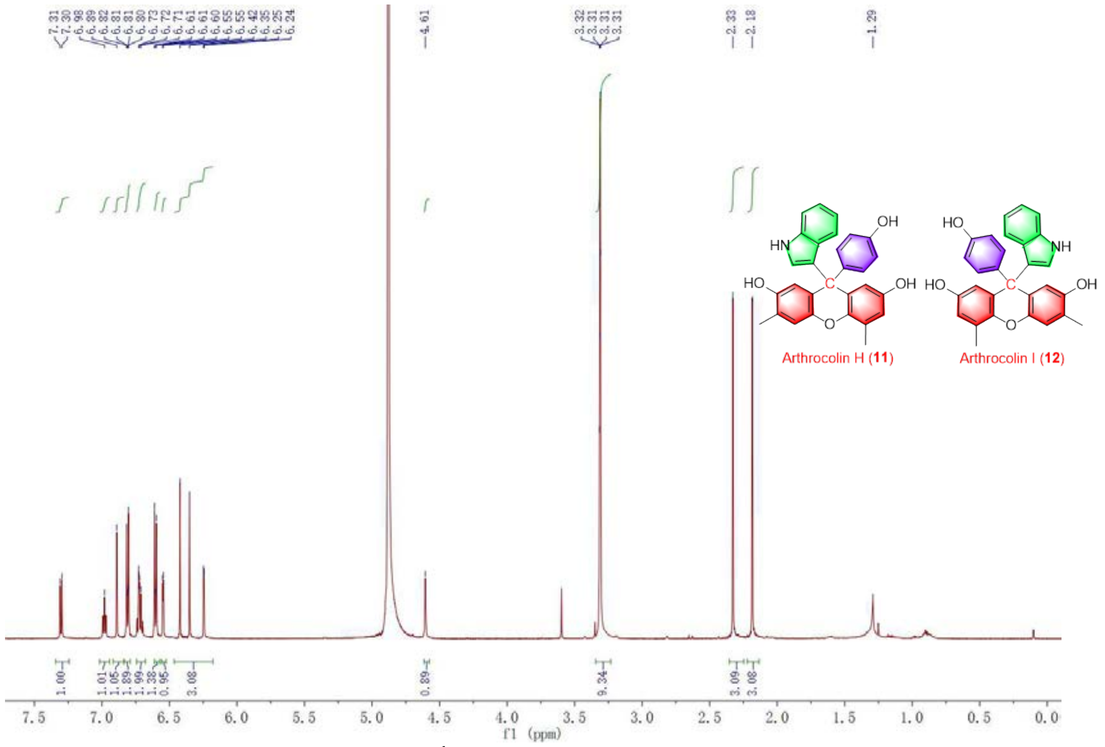


Figure S23. The $^{1}$H NMR spectrum of **11 and 12.**

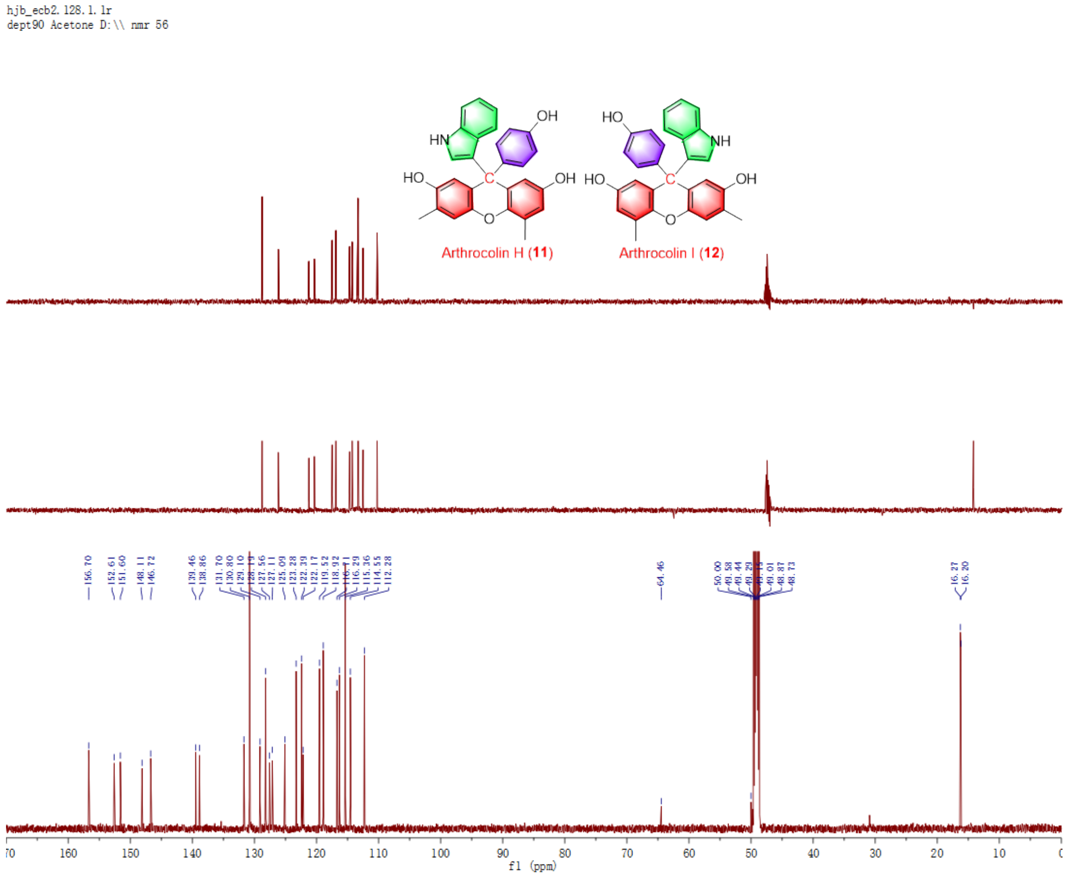


Figure S24. The $^{13}$C NMR spectrum of **11 and 12.**

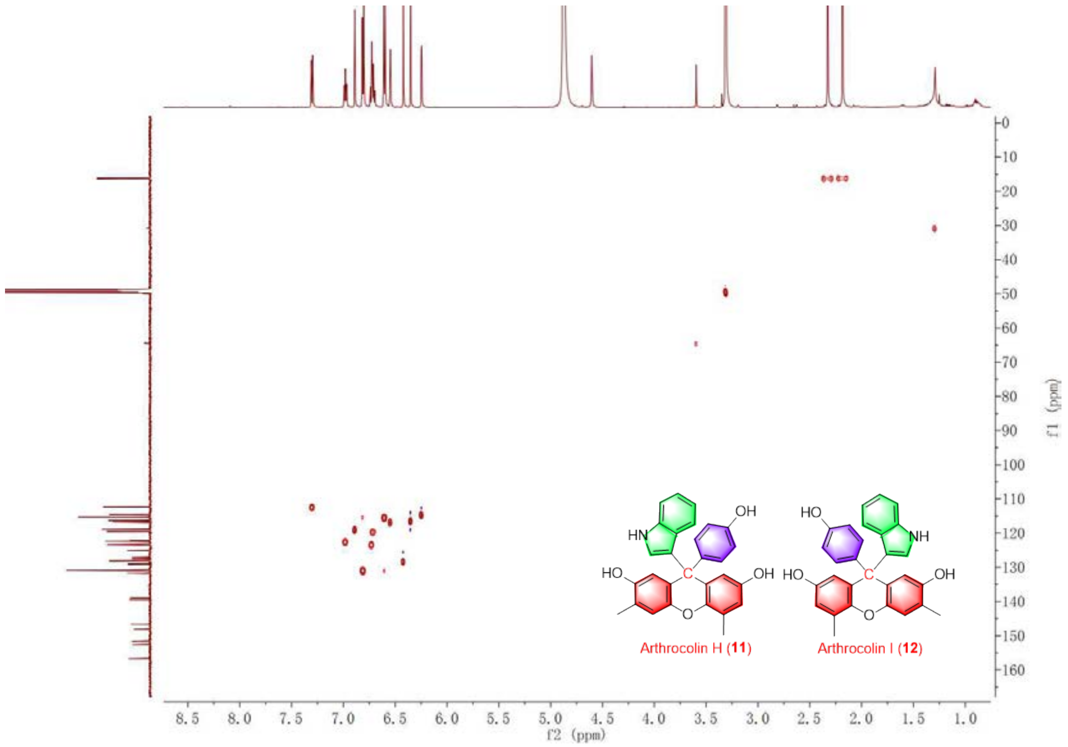


Figure S25. The HSQC spectrum of **11 and 12.**

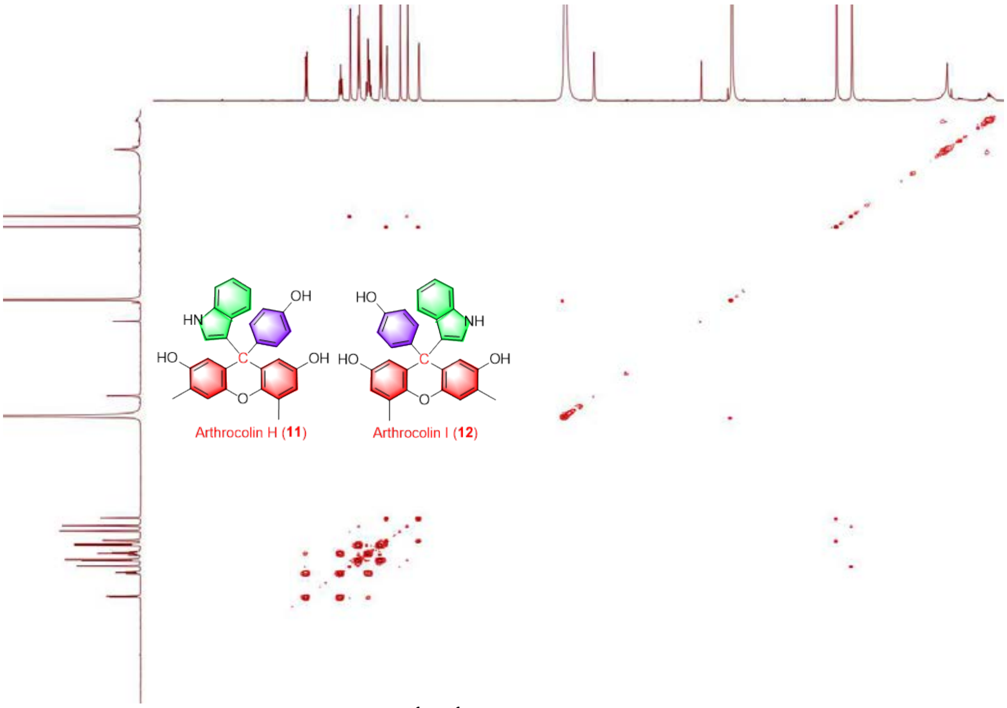


Figure S26. The $^{1}H$-$^{1}H$COSY spectrum of **11 and 12.**

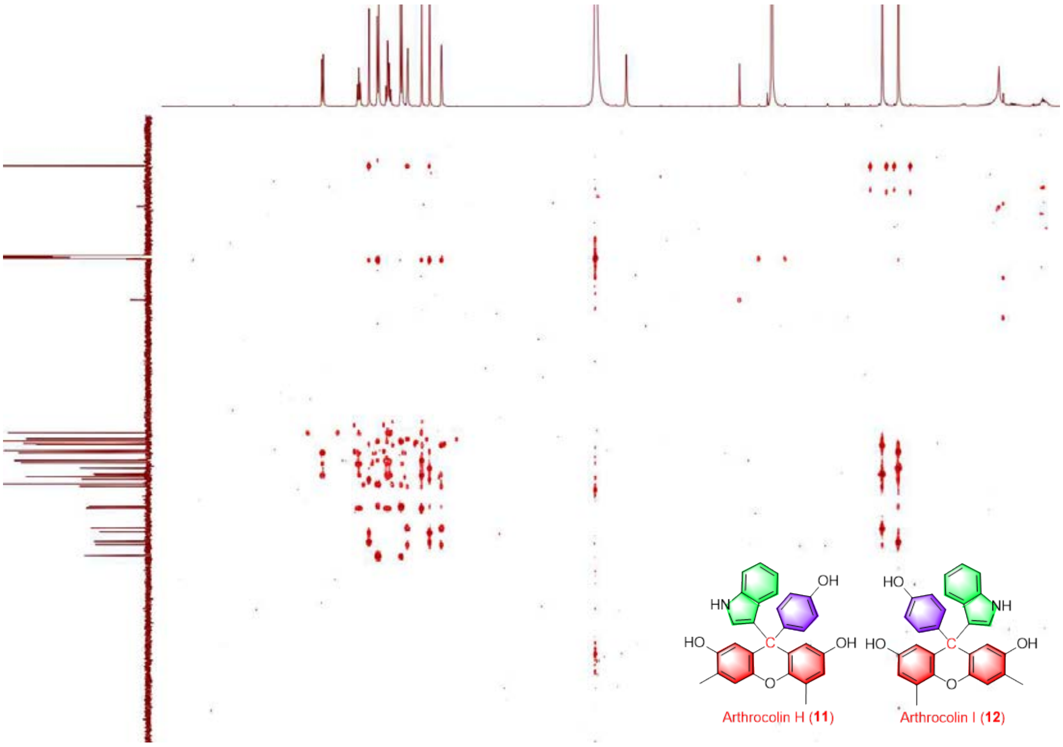


Figure S27. The HMBC spectrum of **11 and 12.**

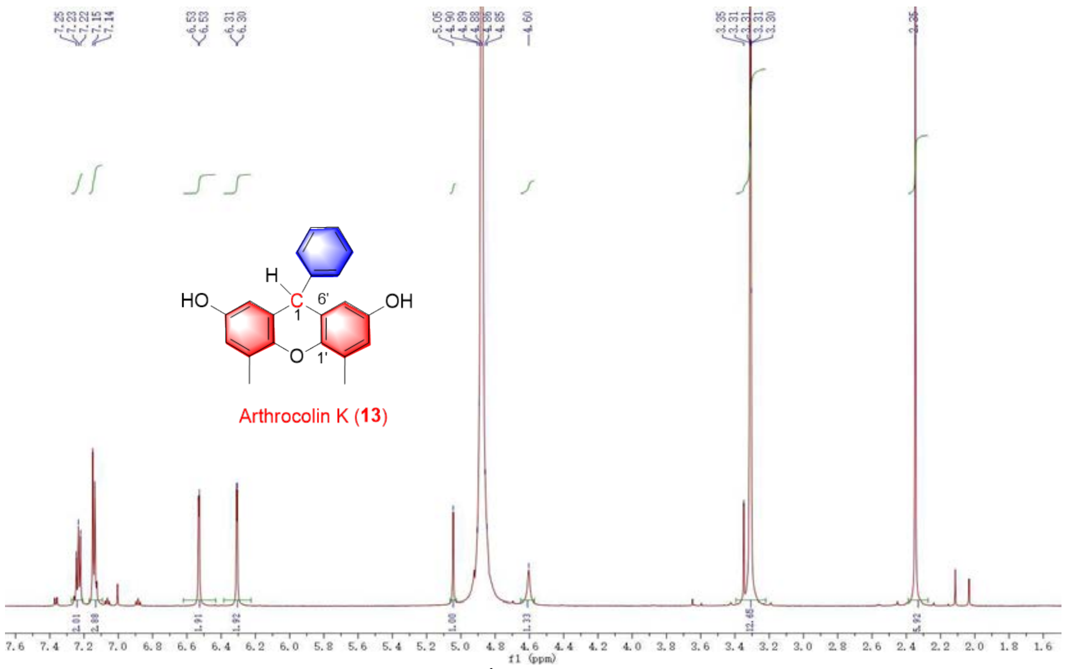


Figure S28. The $^{1}H$ NMR spectrum of **13.**

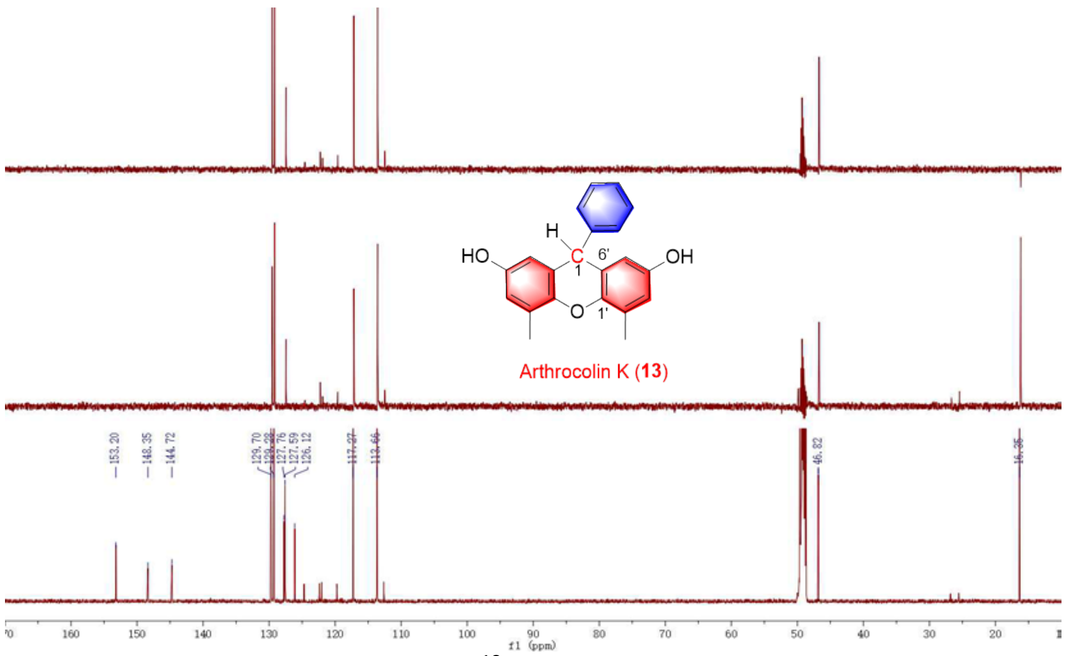


Figure S29. The $^{13}C$ NMR spectrum of **13.**

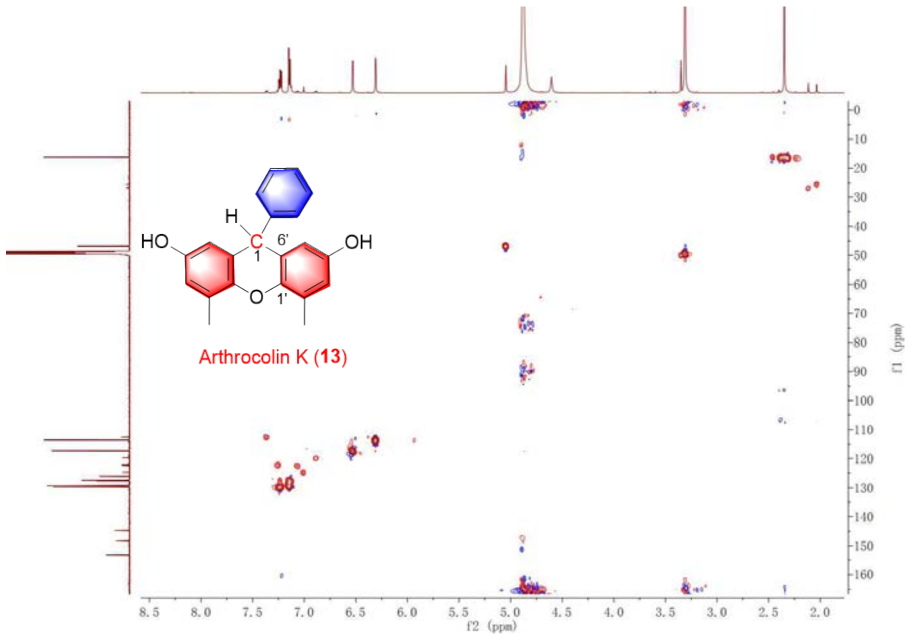


Figure S30. The HSQC spectrum of **13.**

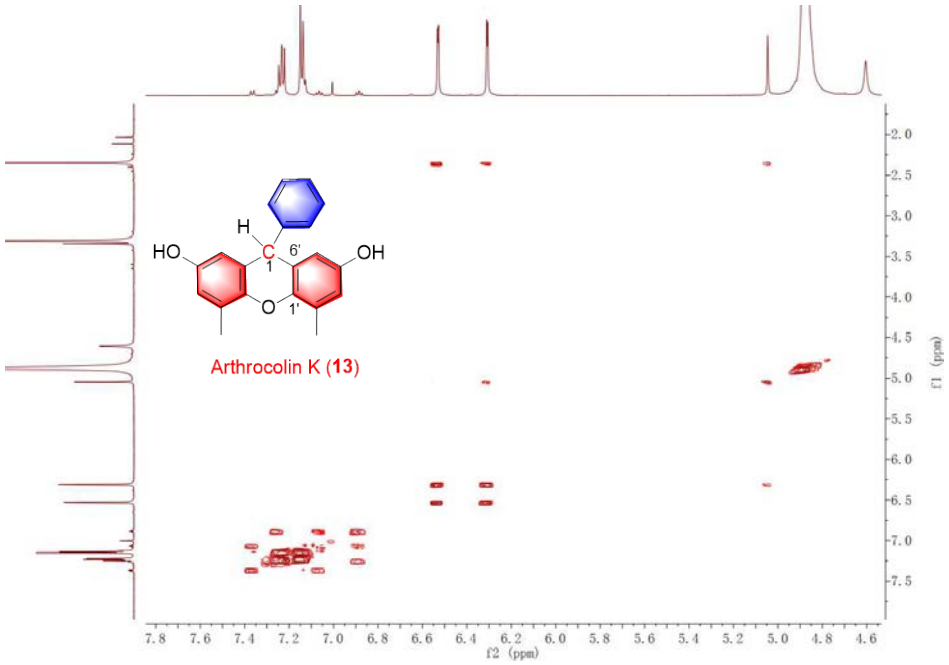


Figure S31. The $^{1}H$-$^{1}H$COSY spectrum of **13.**

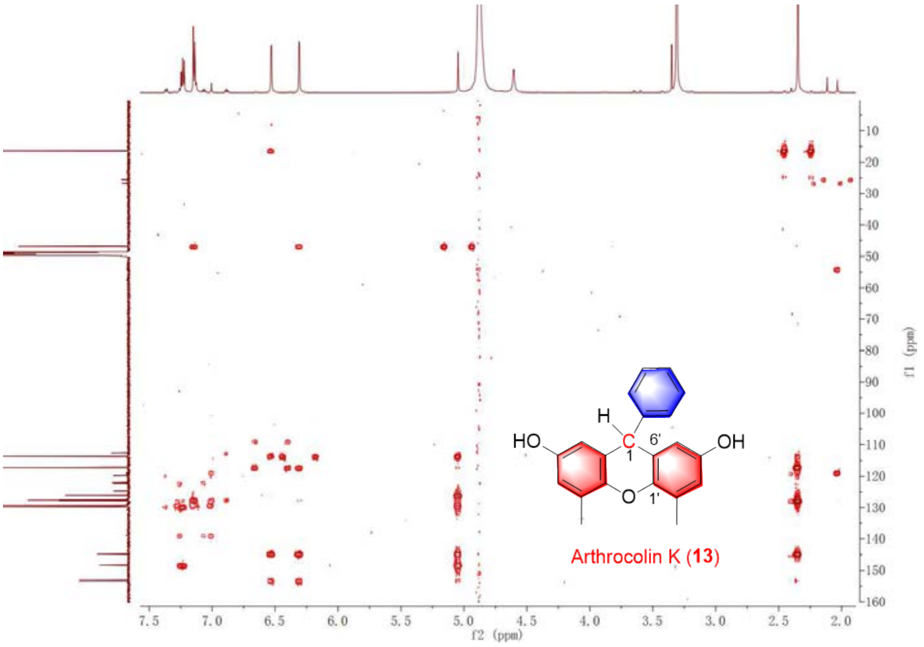


Figure S32. The HMBC spectrum of **13.**

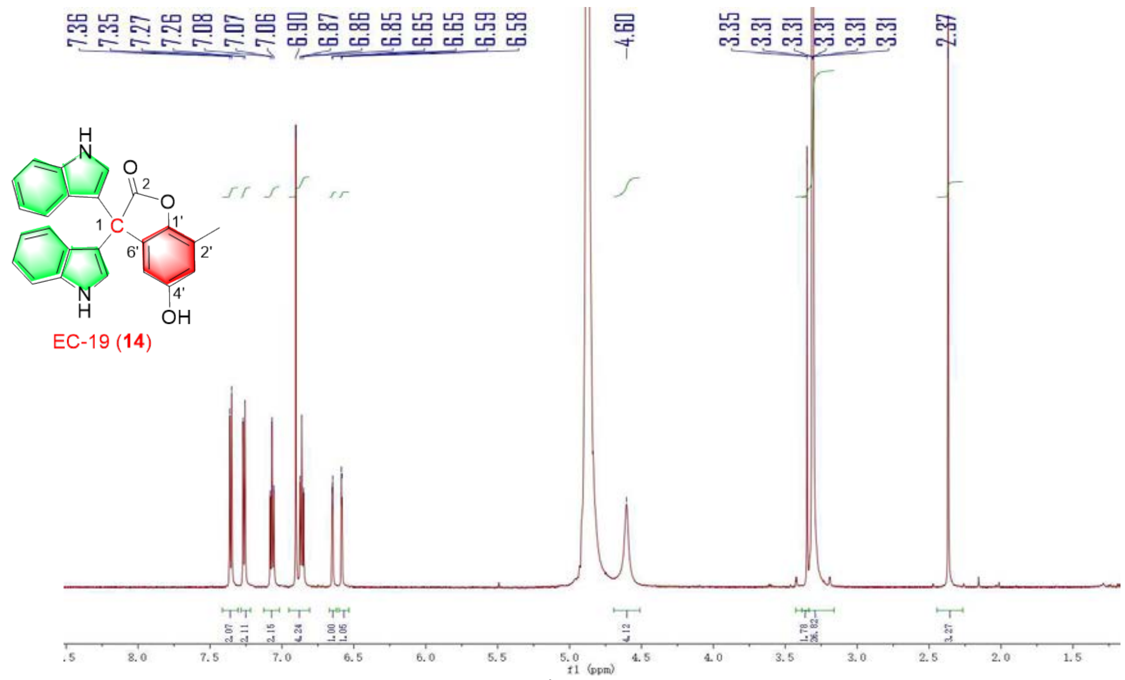


Figure S33. The $^{1}$H NMR spectrum of **14.**

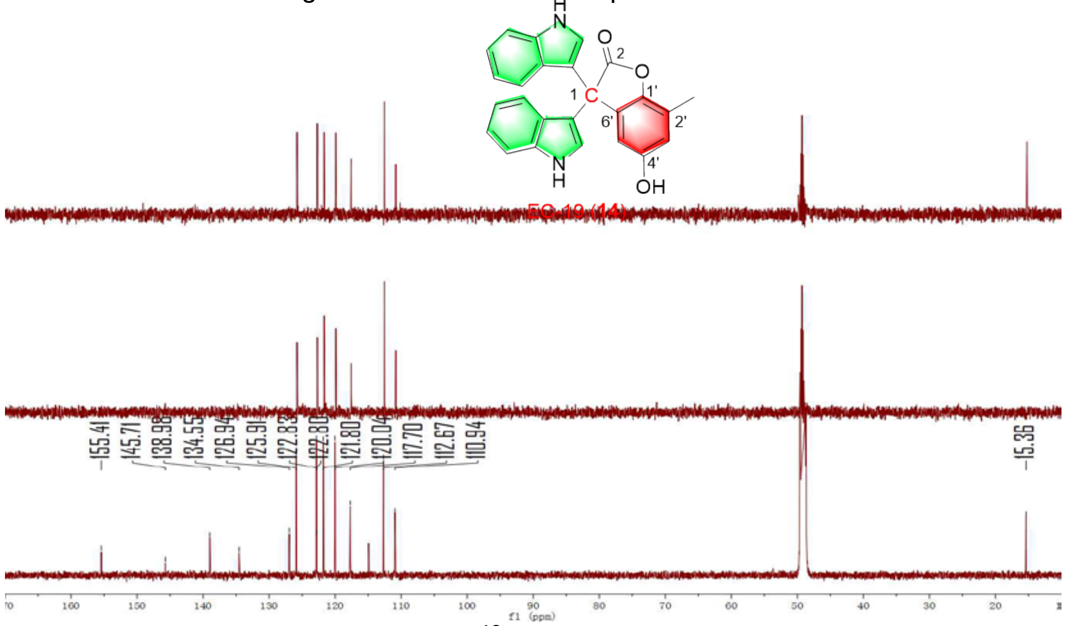


Figure S34. The $^{13}$C NMR spectrum of **14.**

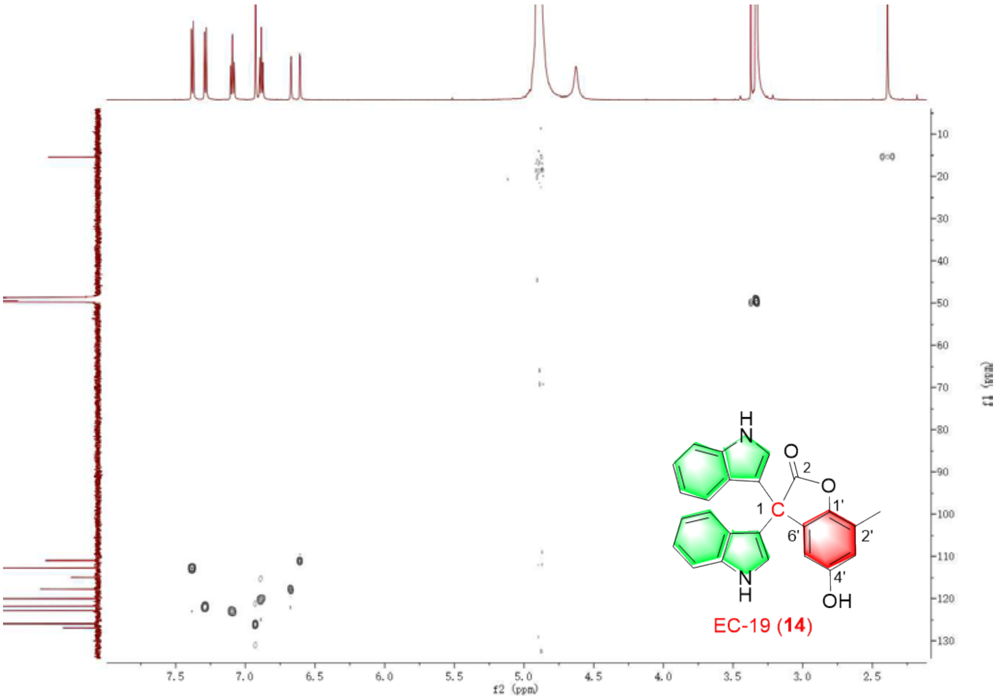


Figure S35. The HSQC spectrum of **14.**

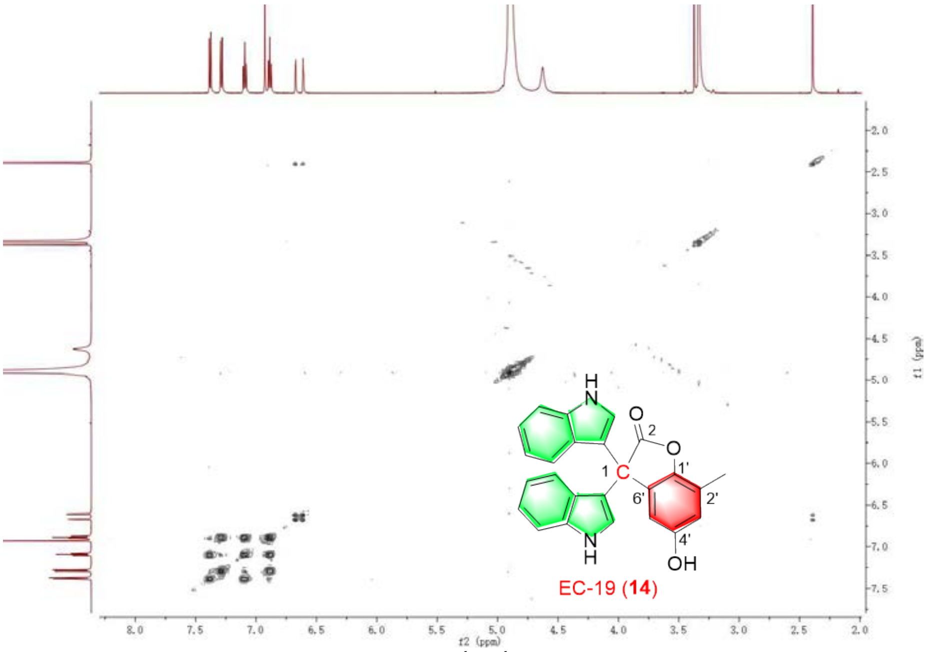


Figure S36. The $^{1}$H-$^{1}$HCOSY spectrum of **14.**

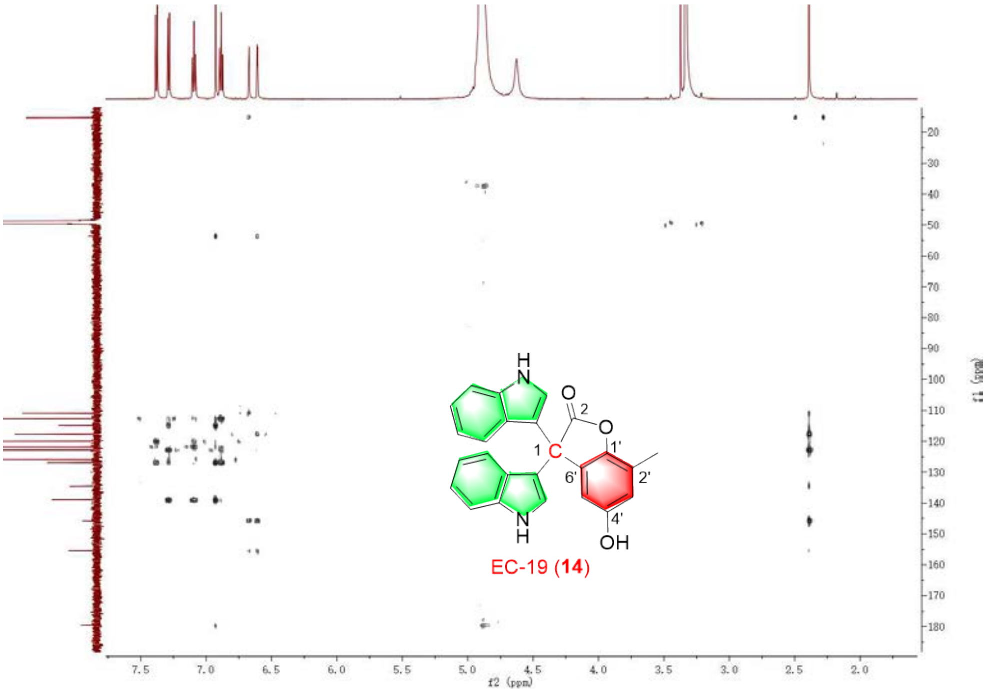


Figure S37. The HMBC spectrum of **14.**

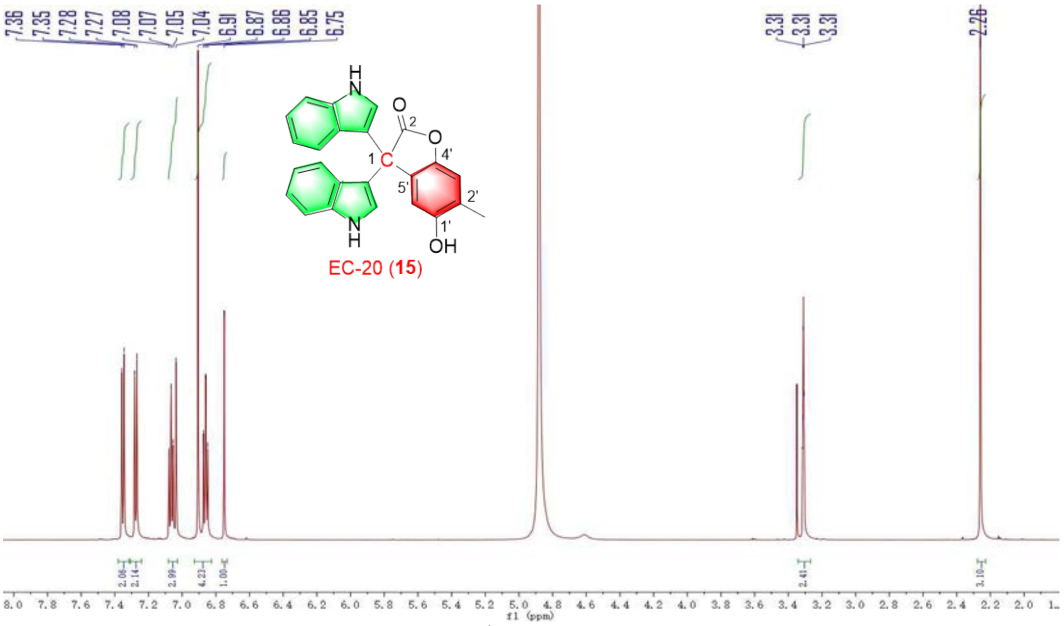


Figure S38. The $^{1}$H NMR spectrum of **15.**

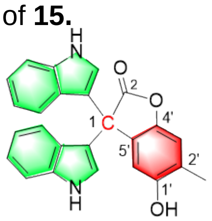

EC-20 (**15**)

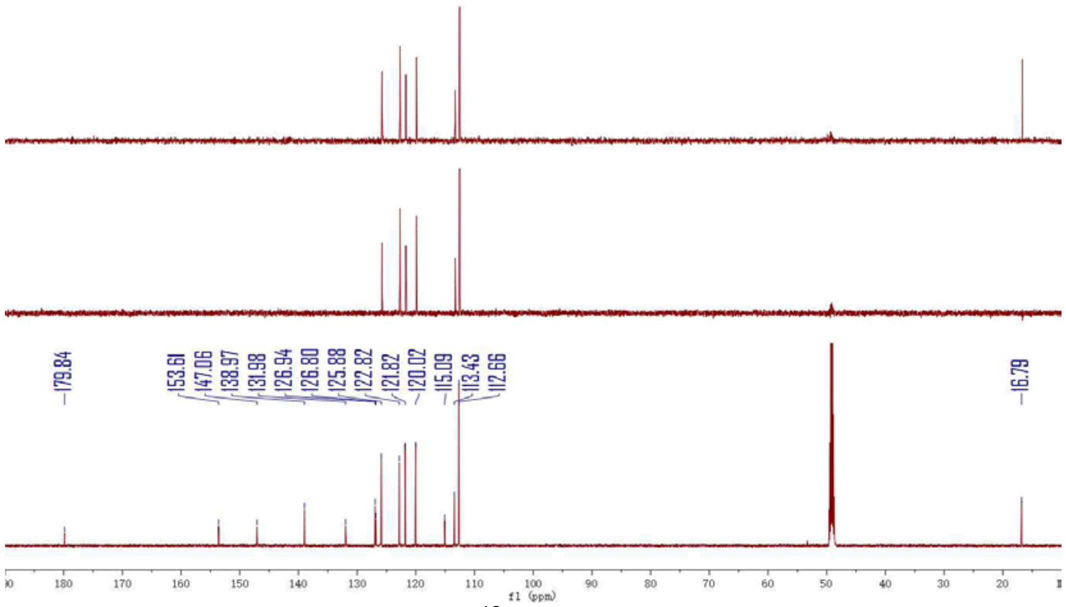


Figure S39. The $^{13}C$ NMR spectrum of **15.**

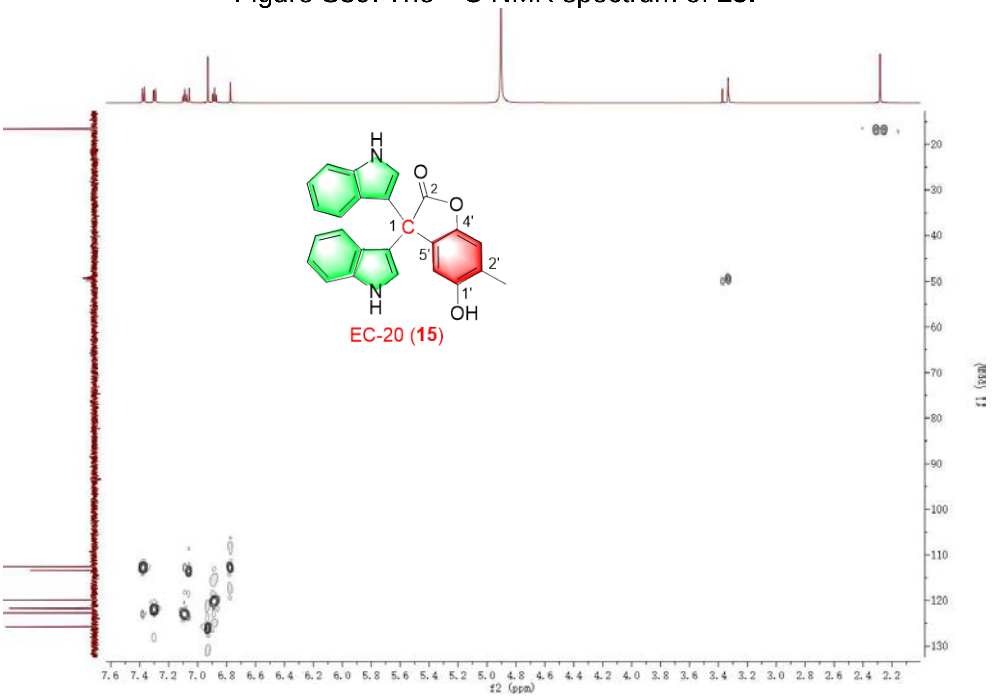


Figure S40. The HSQC spectrum of **15.**

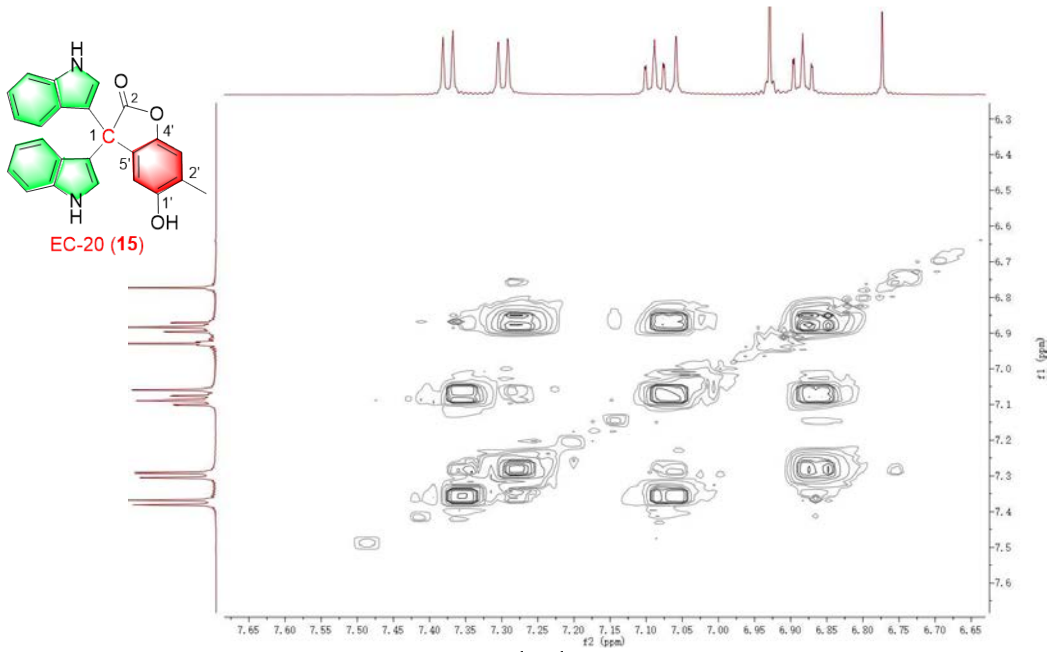

Figure S41. The $^{1}H$-$^{1}H$COSY spectrum of **15.**

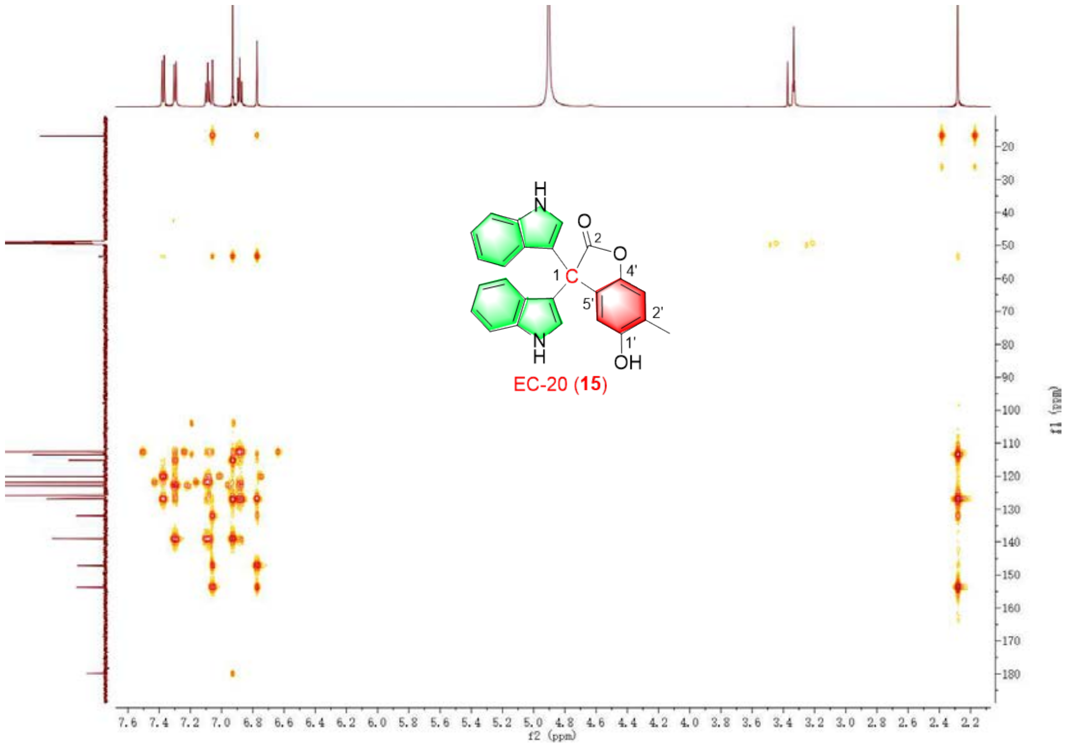

Figure S42. The HMBC spectrum of **15.**

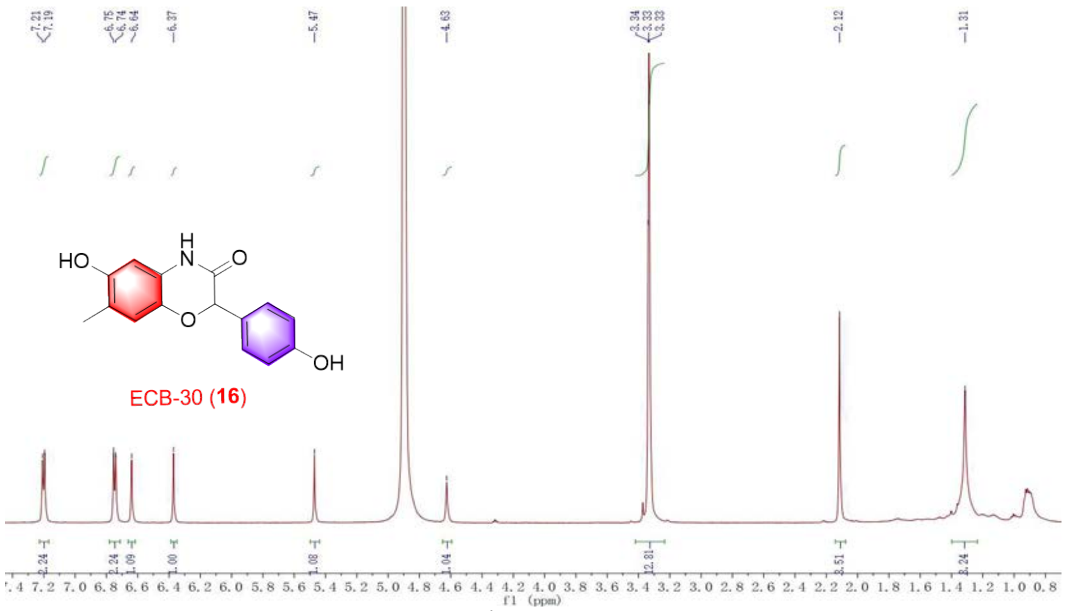


Figure S43. The $^{1}H$ NMR spectrum of **16.**

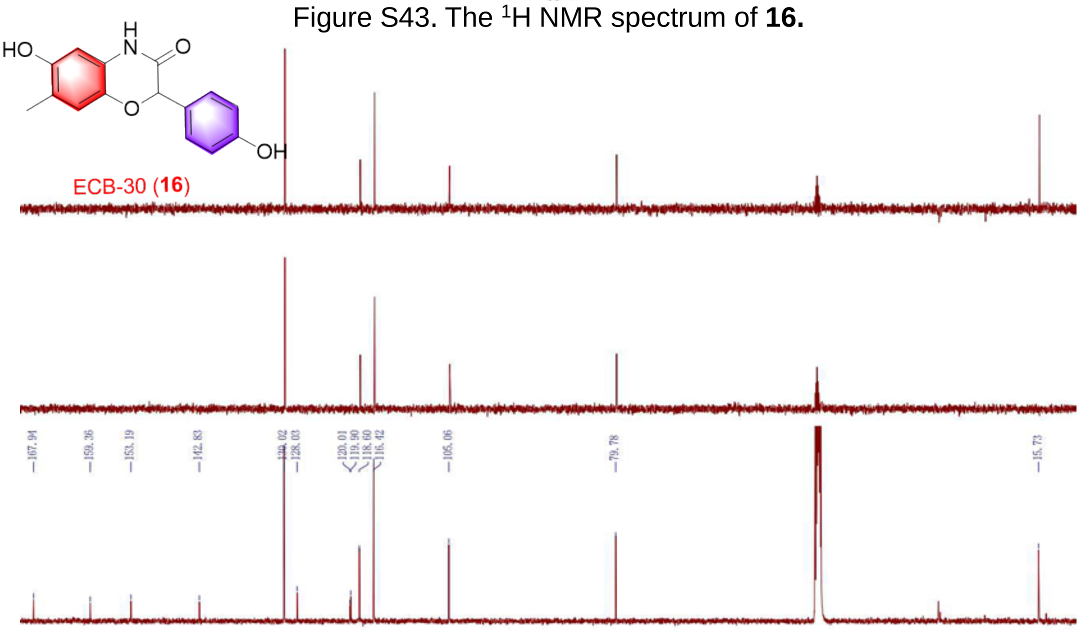


Figure S44. The $^{13}C$ NMR spectrum of **16.**

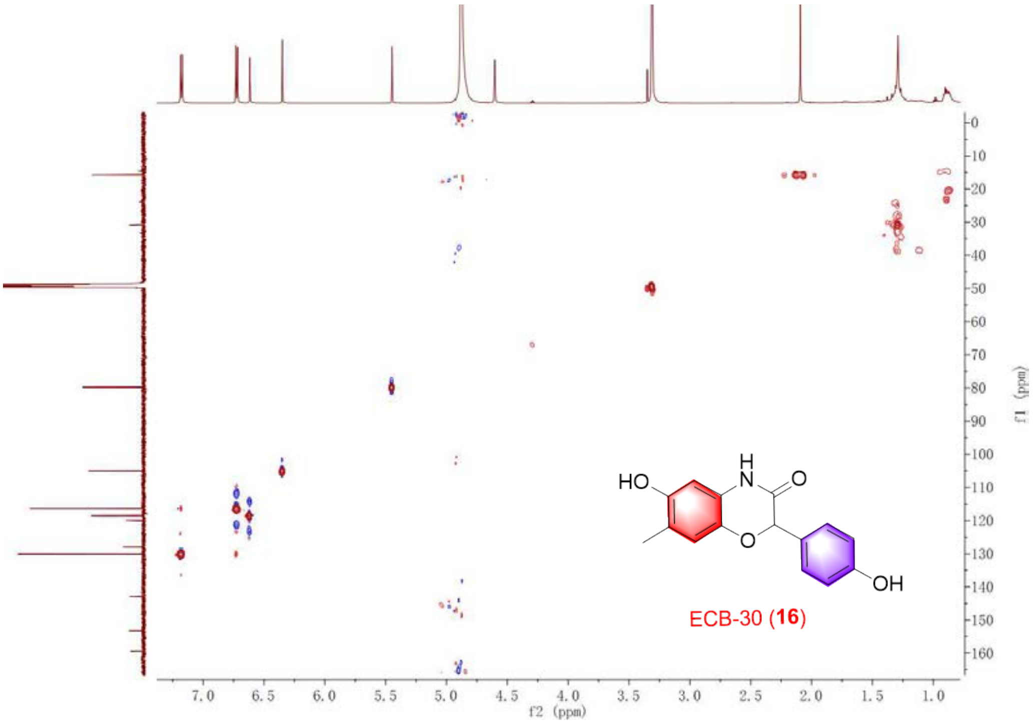

Figure S45. The HSQC spectrum of **16.**

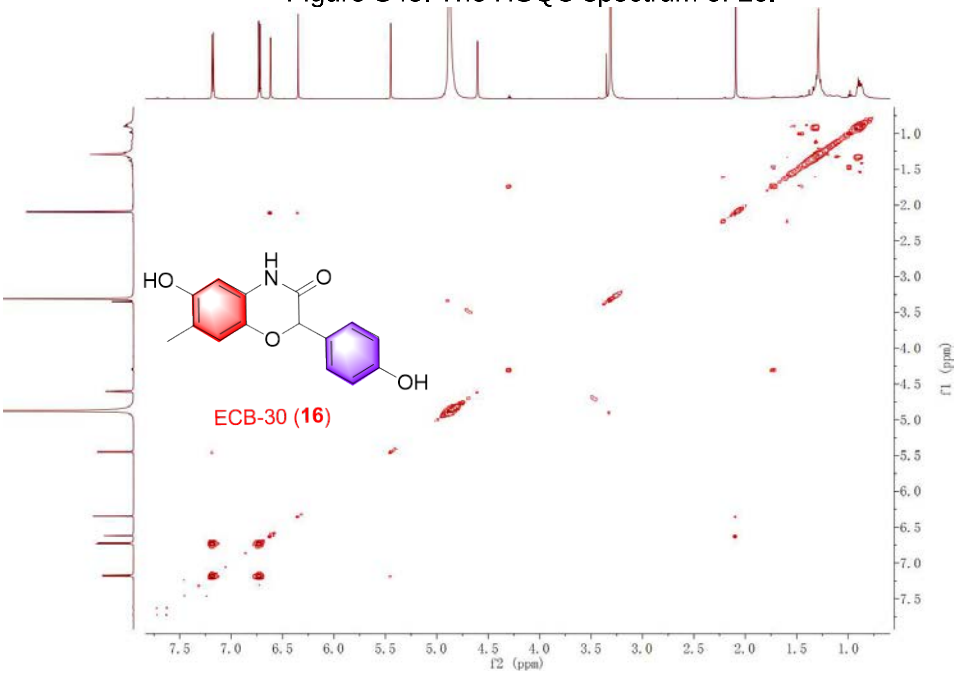

Figure S46. The $^{1}H$-$^{1}H$COSY spectrum of **16.**

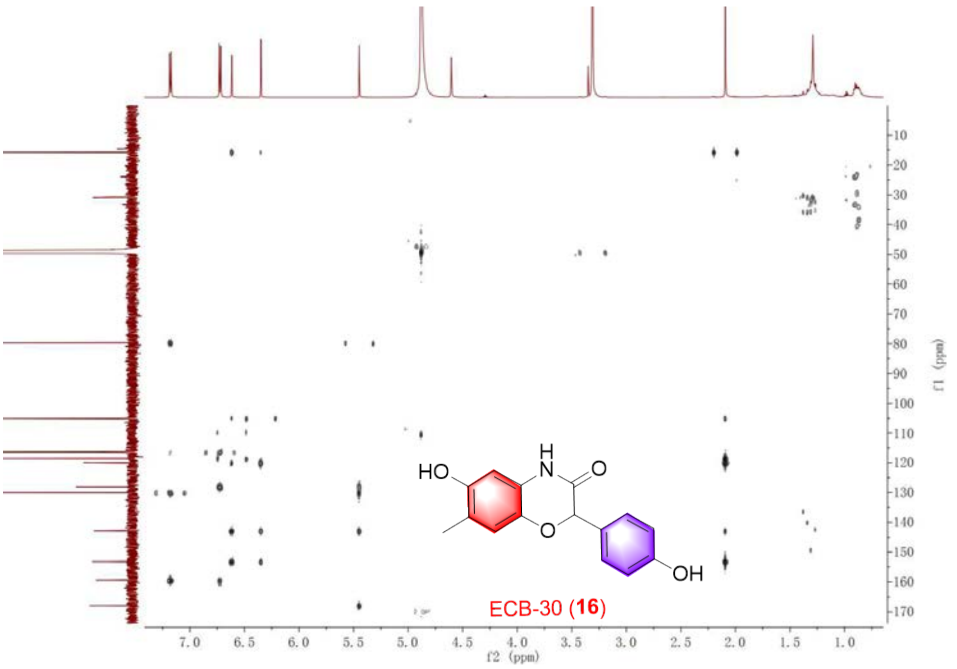


Figure S47. The HMBC spectrum of **16.**

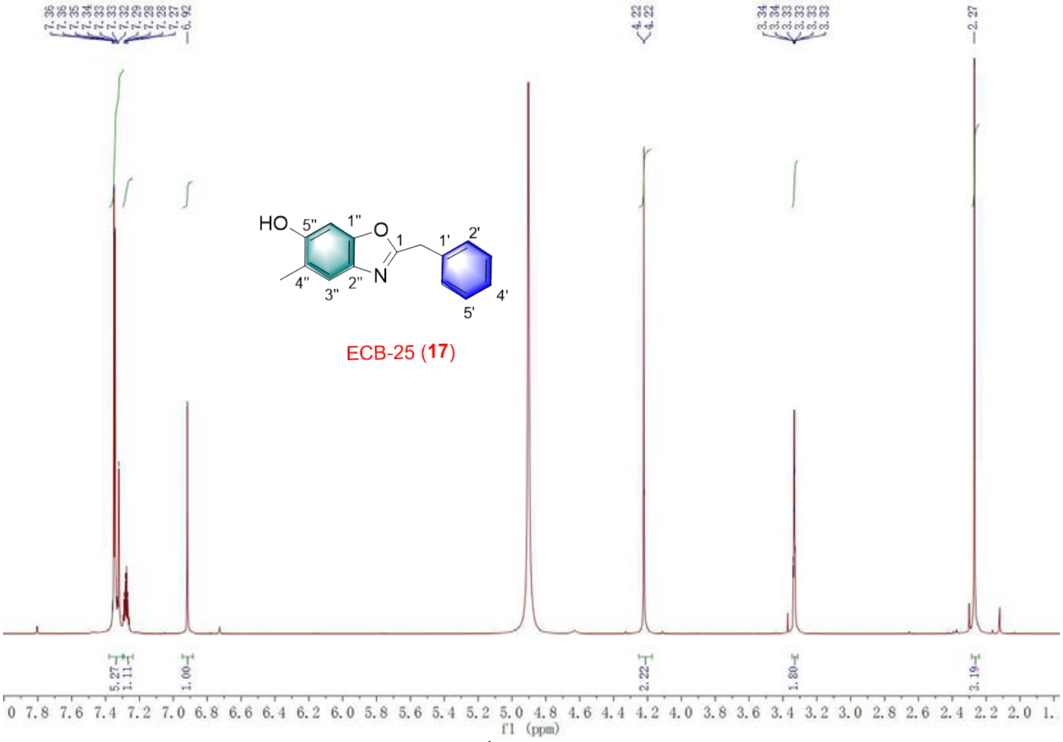


Figure S48. The $^{1}$H NMR spectrum of **17.**

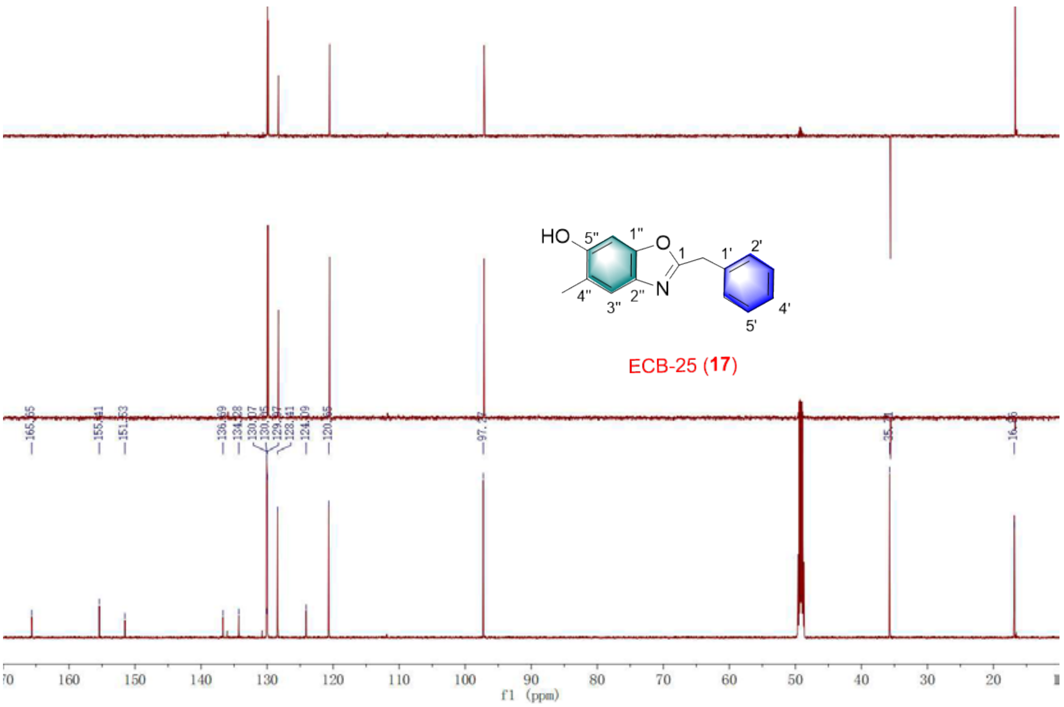


Figure S49. The $^{13}$C NMR spectrum of **17.**

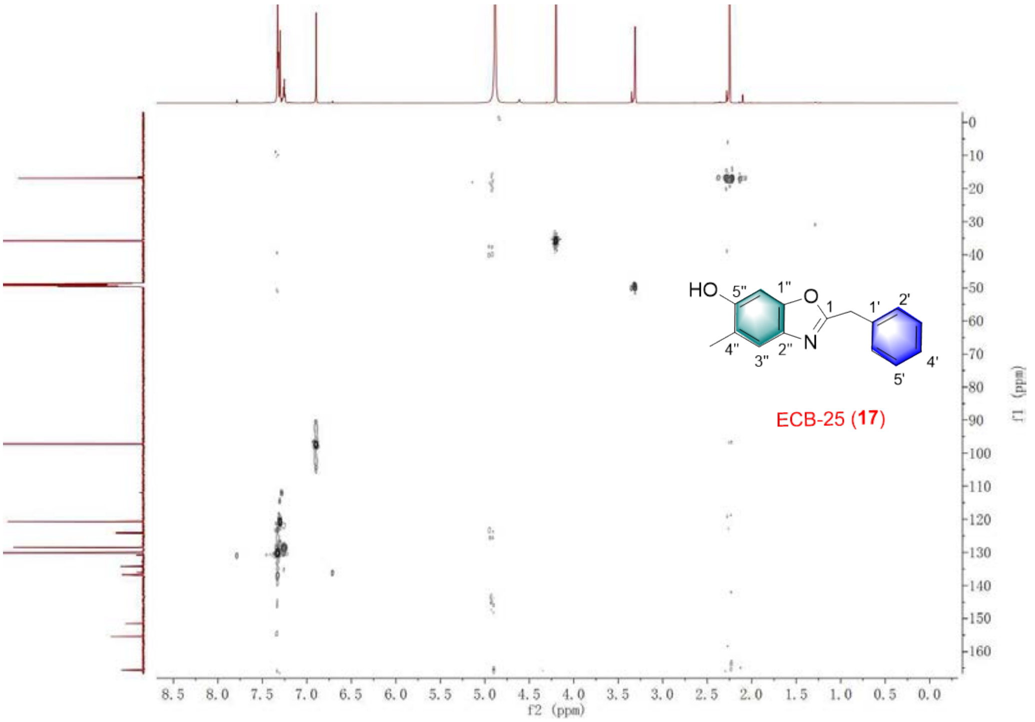


Figure S50. The HSQC spectrum of **17.**

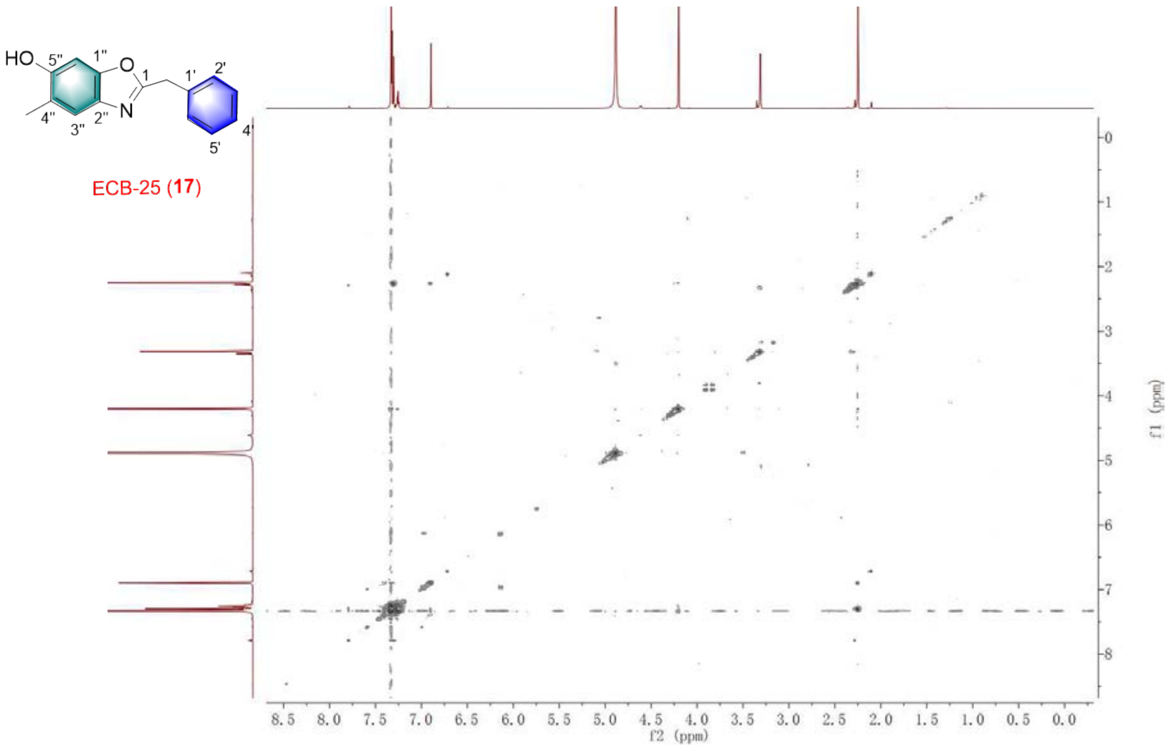


Figure S51. The $^{1}H$-$^{1}H$COSY spectrum of **17.**

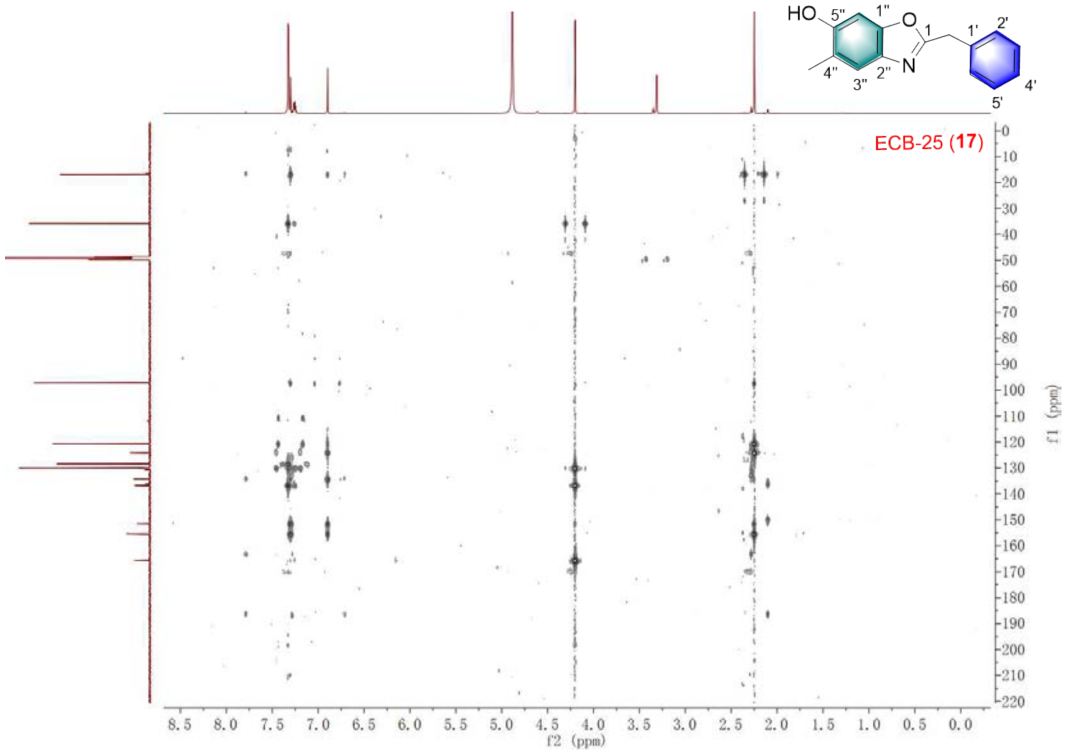


Figure S52. The HMBC spectrum of **17.**

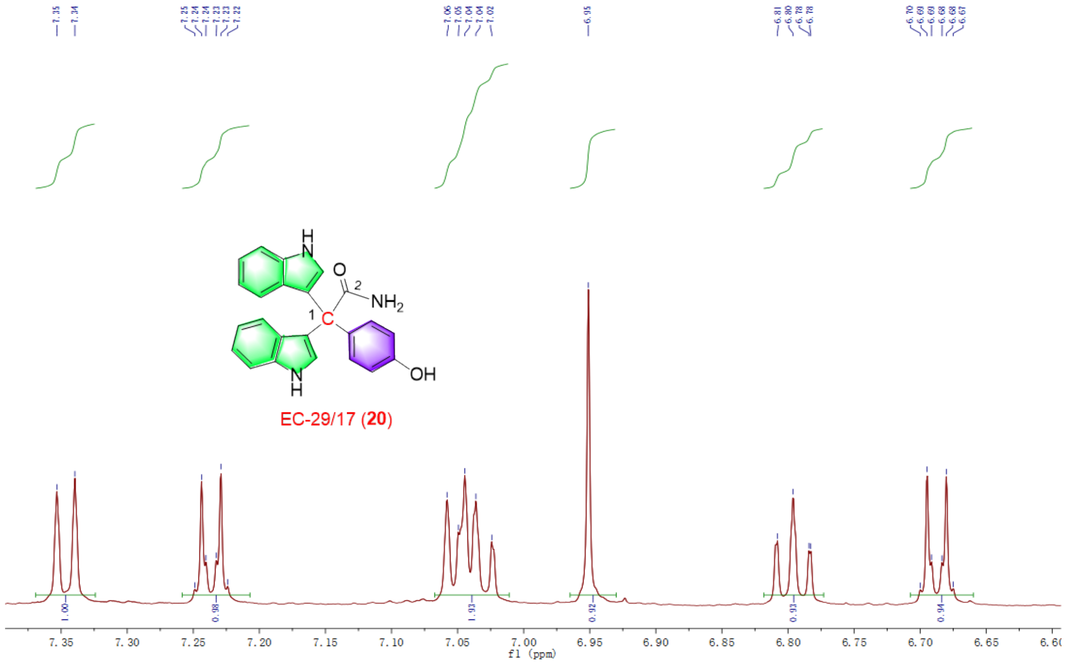


Figure S53. The $^{1}$H NMR spectrum of **20.**

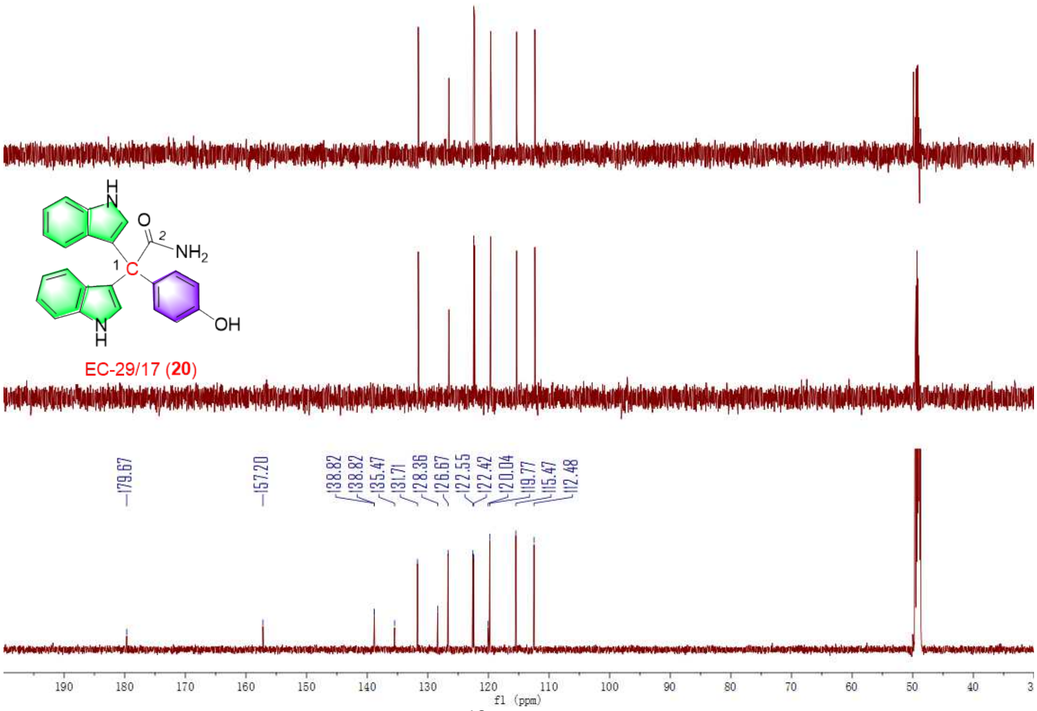


Figure S54. The $^{13}$C NMR spectrum of **20.**

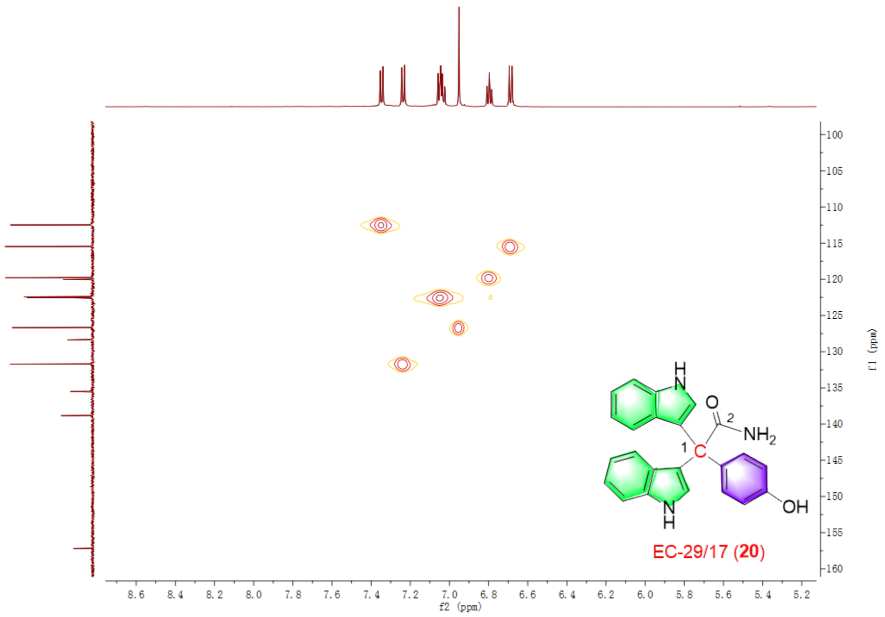


Figure S55. The HSQC spectrum of **20.**

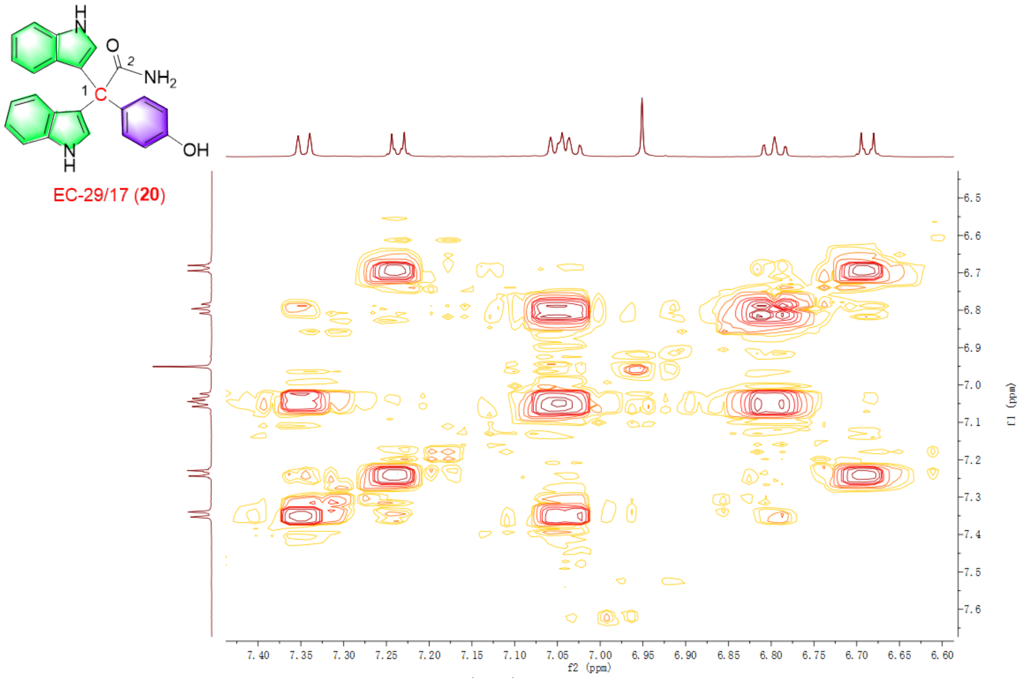


Figure S56. The $^{1}H$-$^{1}H$COSY spectrum of **20.**

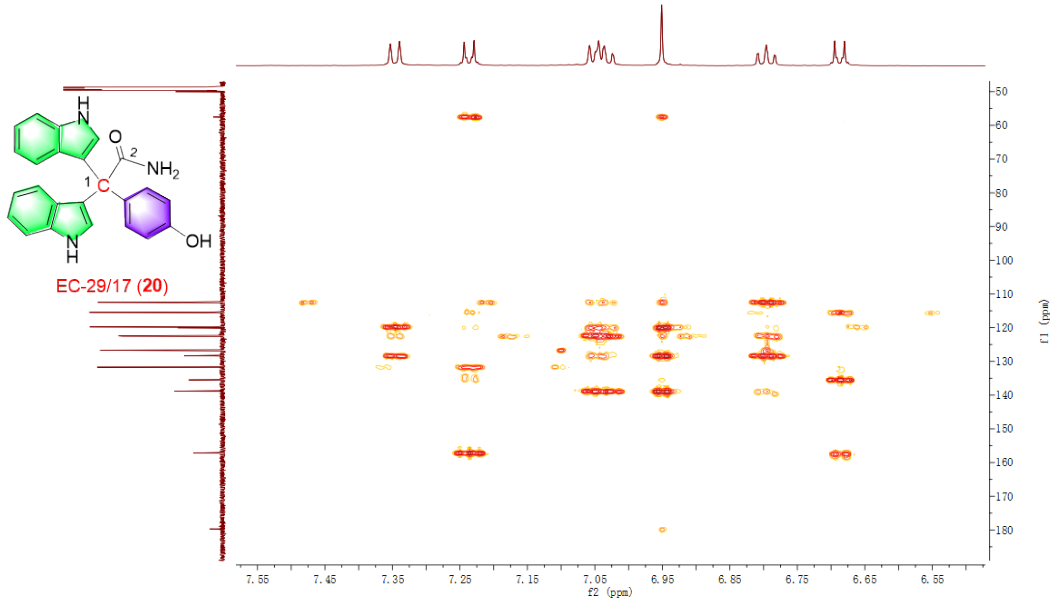


Figure S57. The HMBC spectrum of **20.**

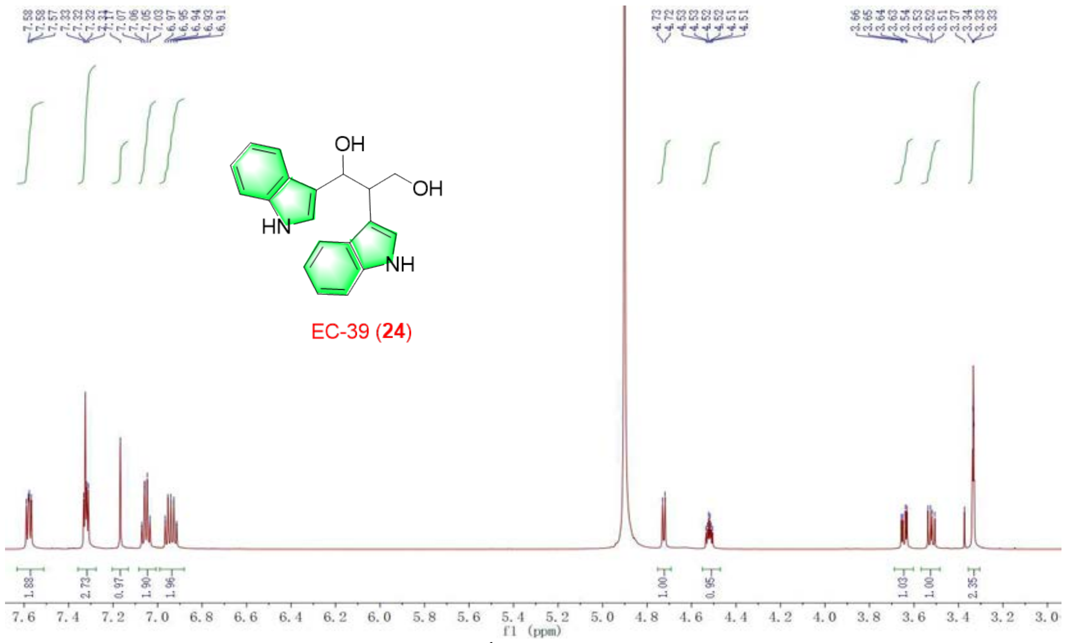


Figure S58. The $^{1}H$ NMR spectrum of **24.**

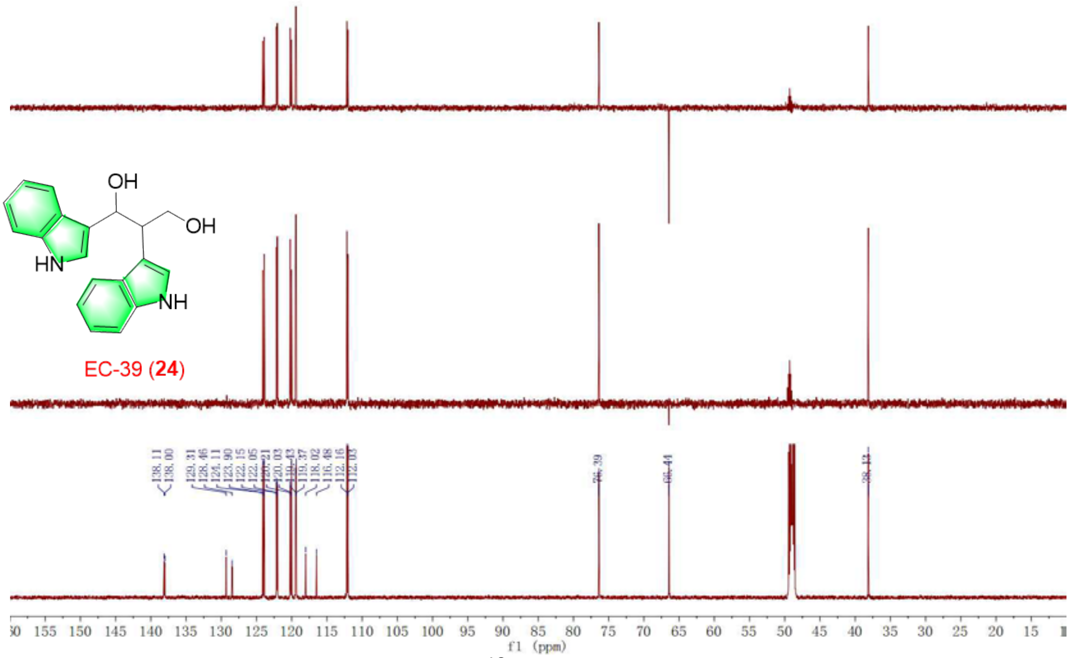

Figure S59. The $^{13}C$ NMR spectrum of **24.**

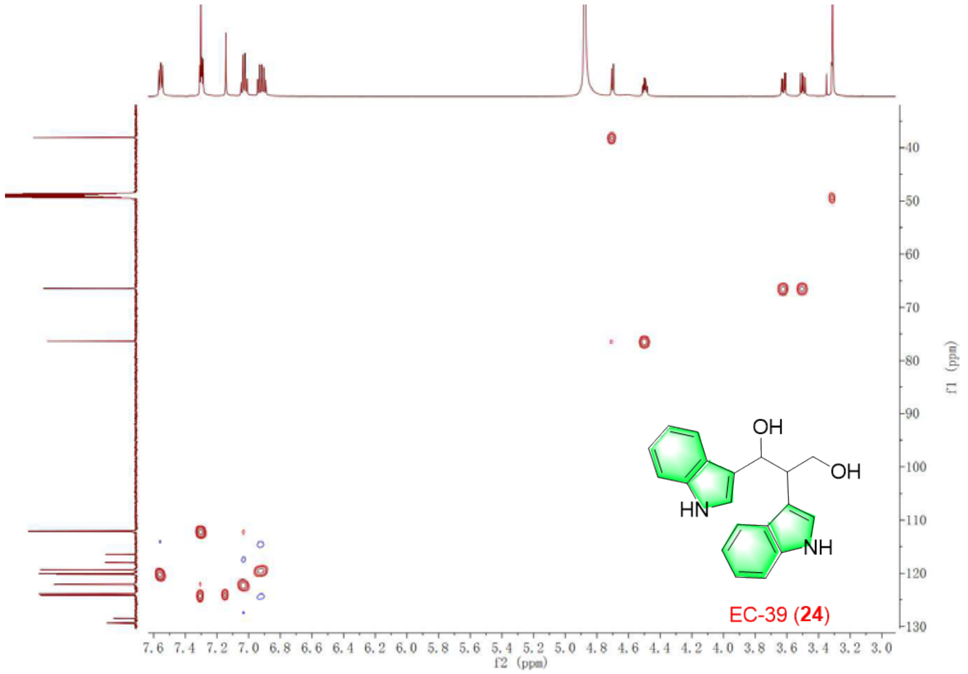

Figure S60. The HSQC spectrum of **24.**

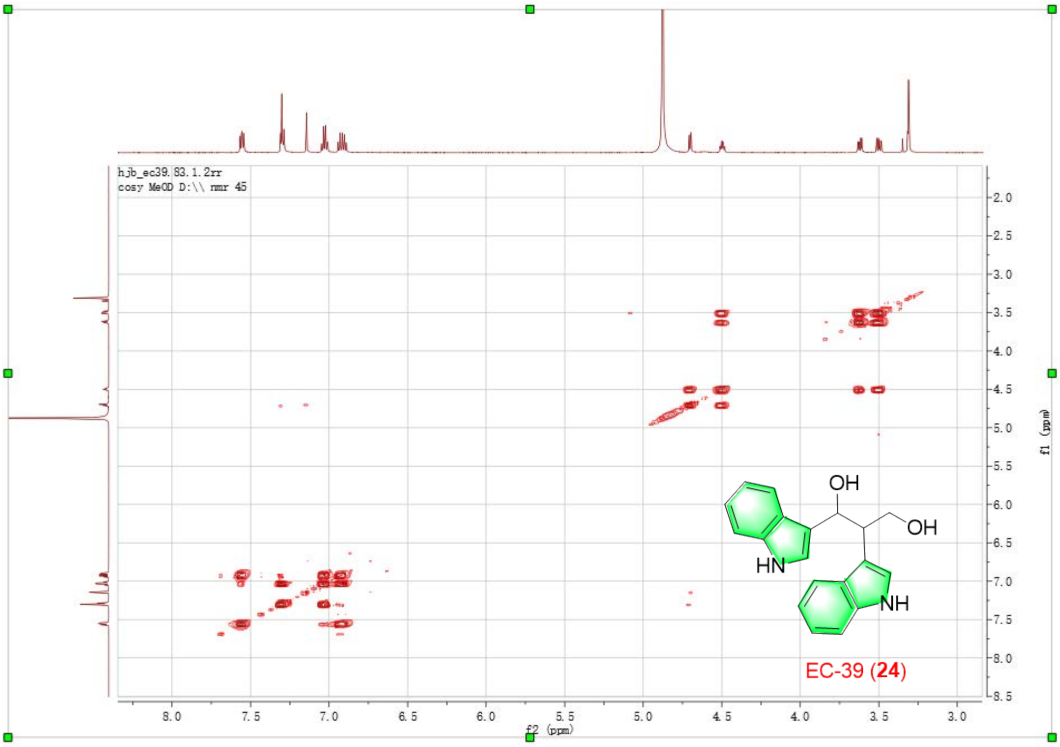


Figure S61. The $^{1}$H-$^{1}$HCOSY spectrum of **24.**

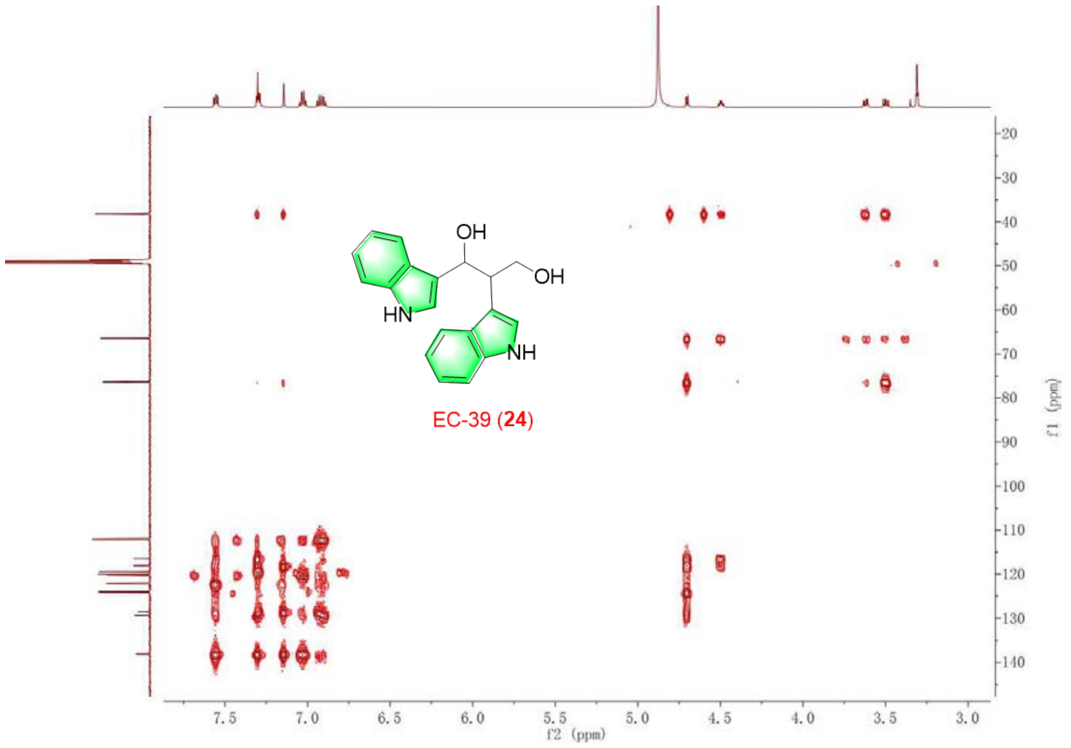


Figure S62. The HMBC spectrum of **24.**

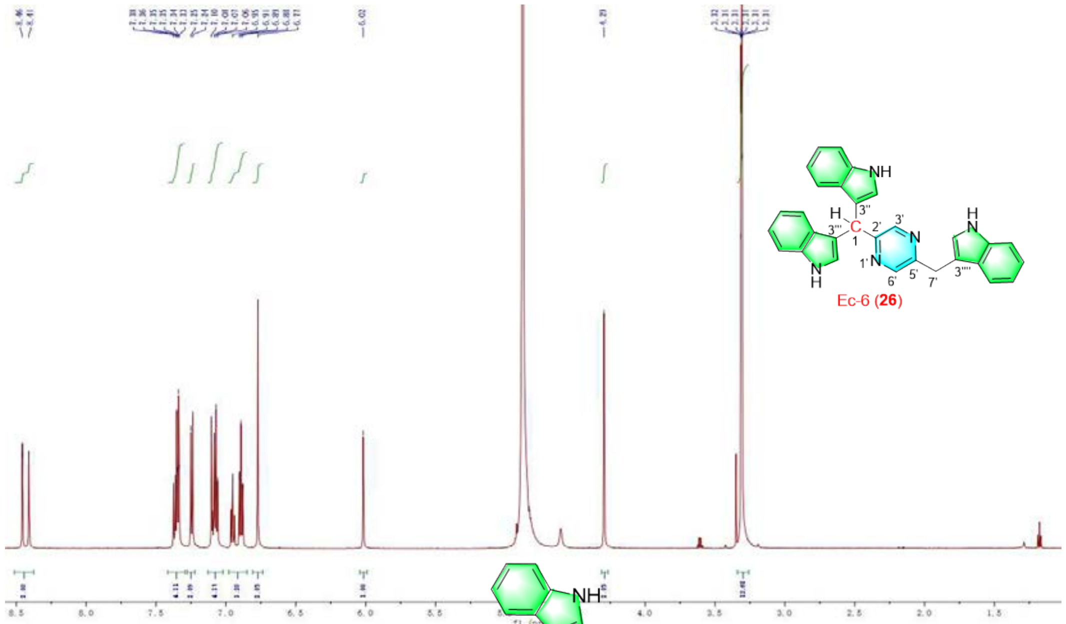


Figure S63. The $^{1}$H NMR spectrum of **26.**

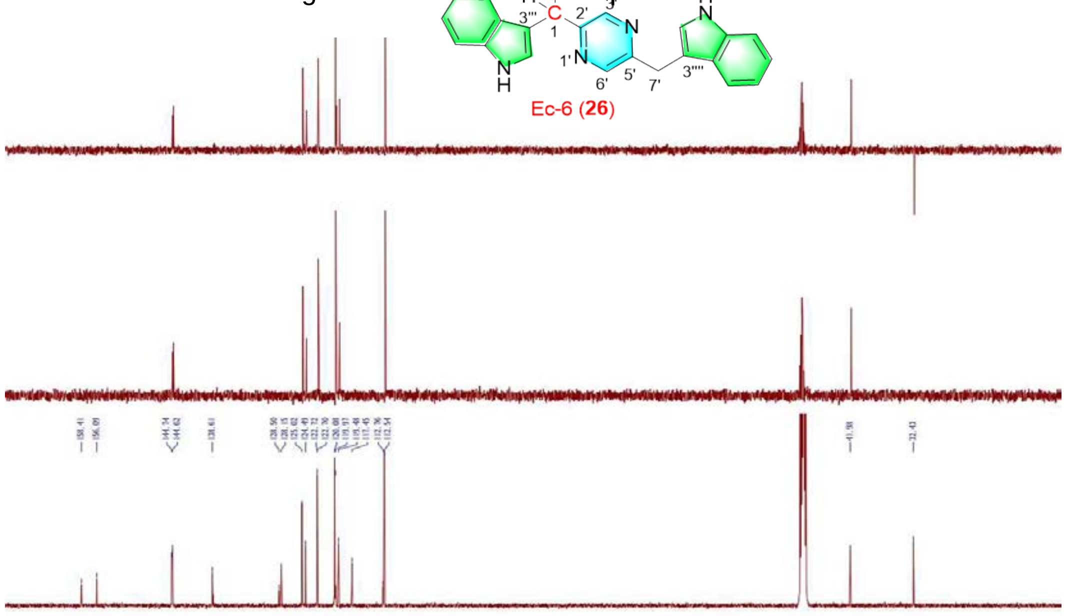


Figure S64. The $^{13}$C NMR spectrum of **26.**

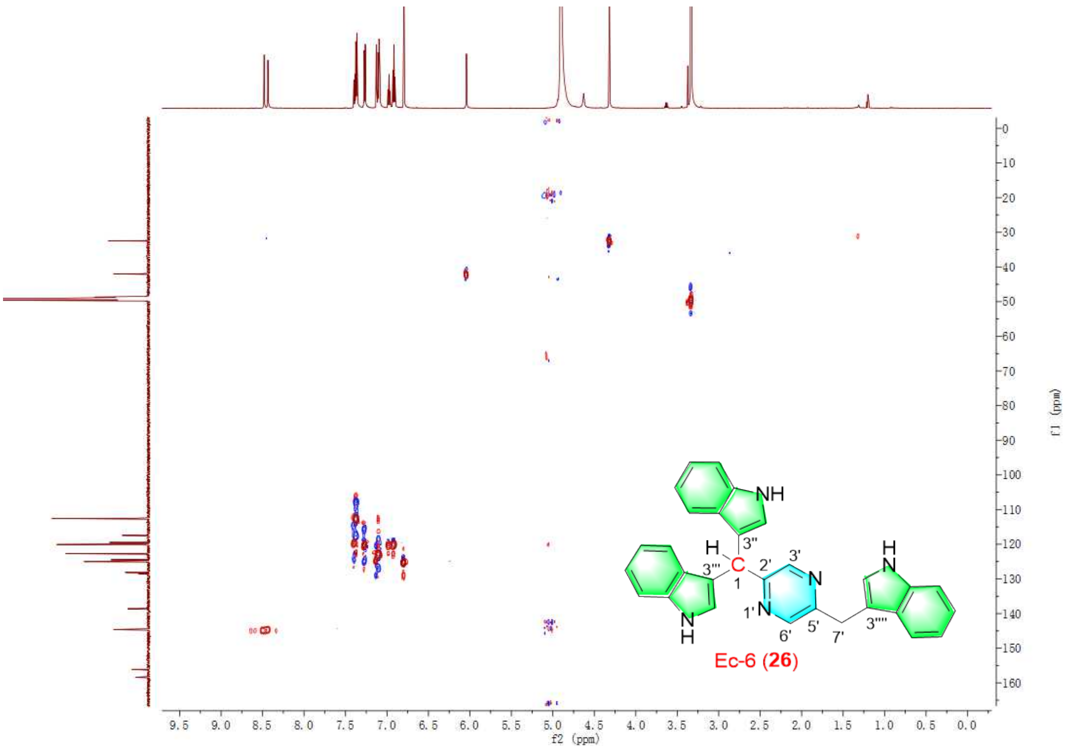


Figure S65. The HSQC spectrum of **26.**

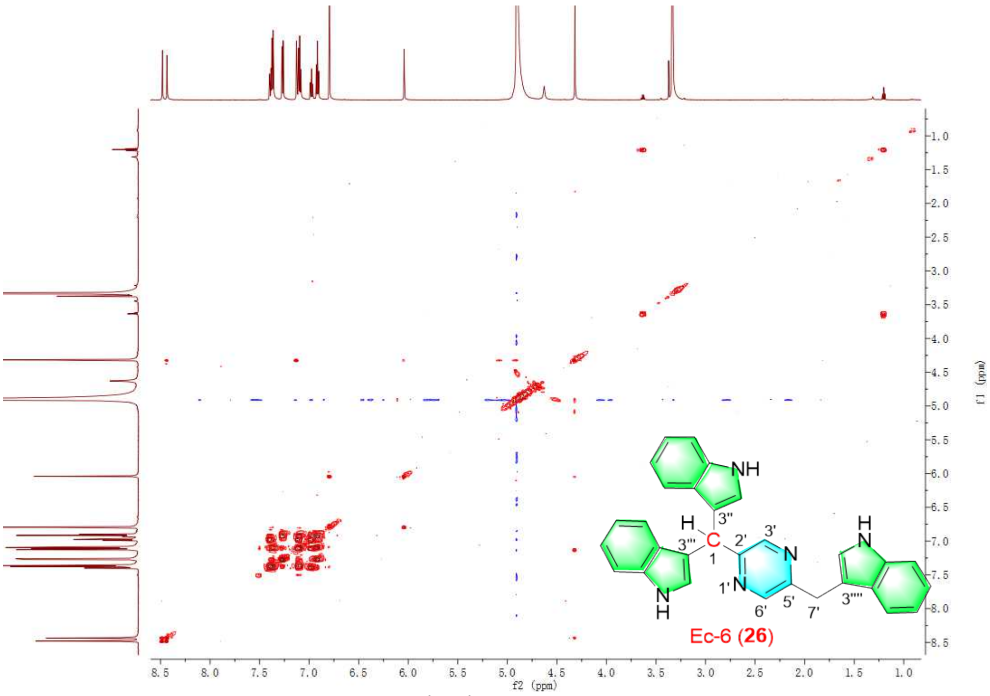


Figure S66. The $^{1}$H-$^{1}$HCOSY spectrum of **26.**

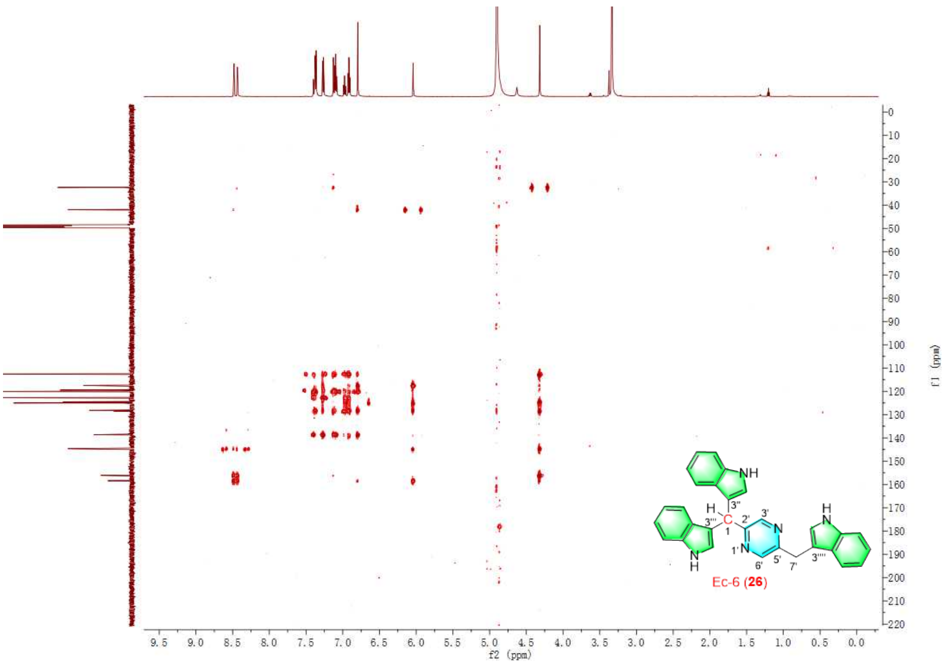

Figure S67. The HMBC spectrum of **26.**

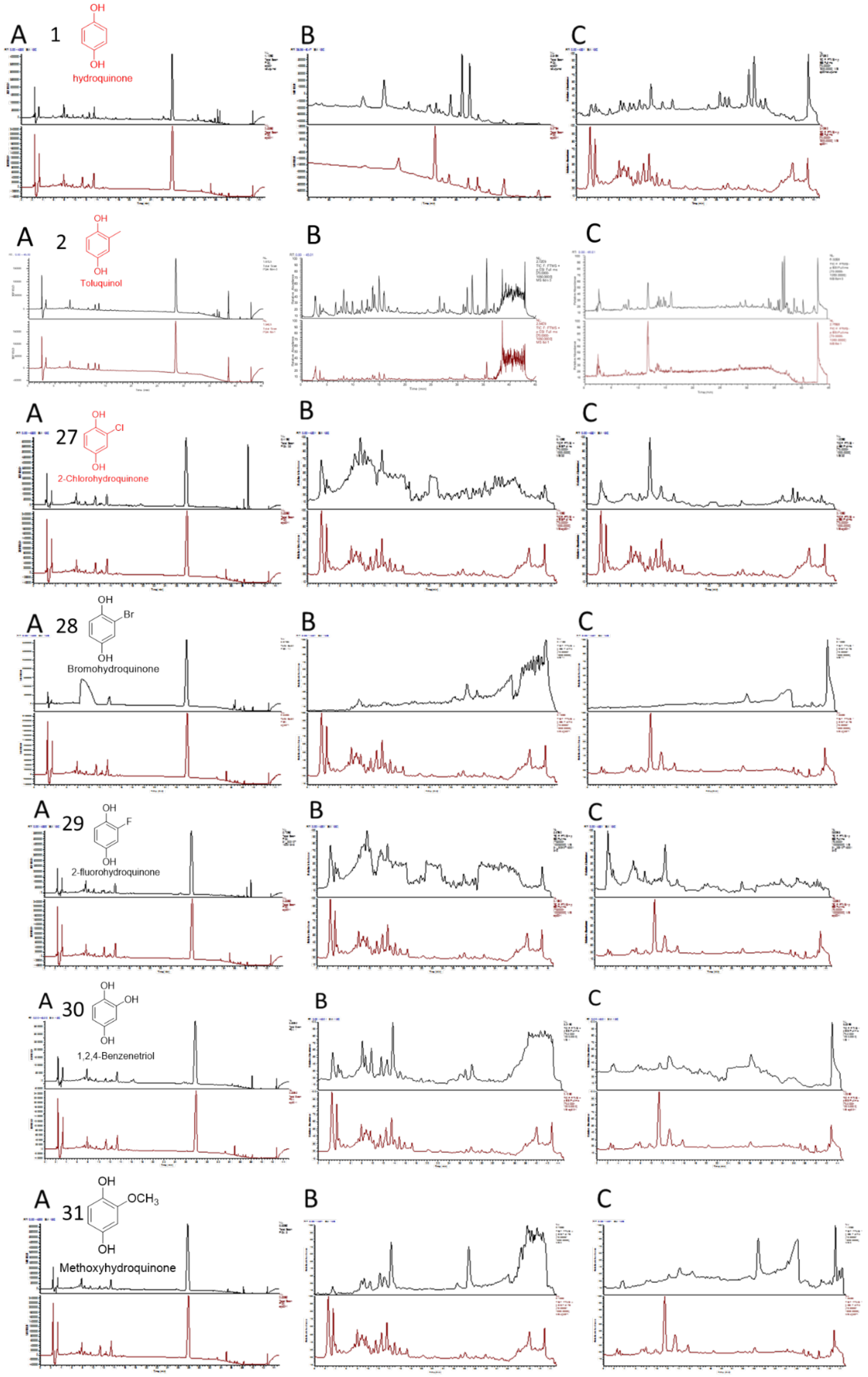


Figure S68. The LC-PDA/MS analysis of *E. coli* fed with compounds **1-2** and **27-31**. A: PDA analysis; B: Positive MS analysis; C: B: Negative MS analysis.

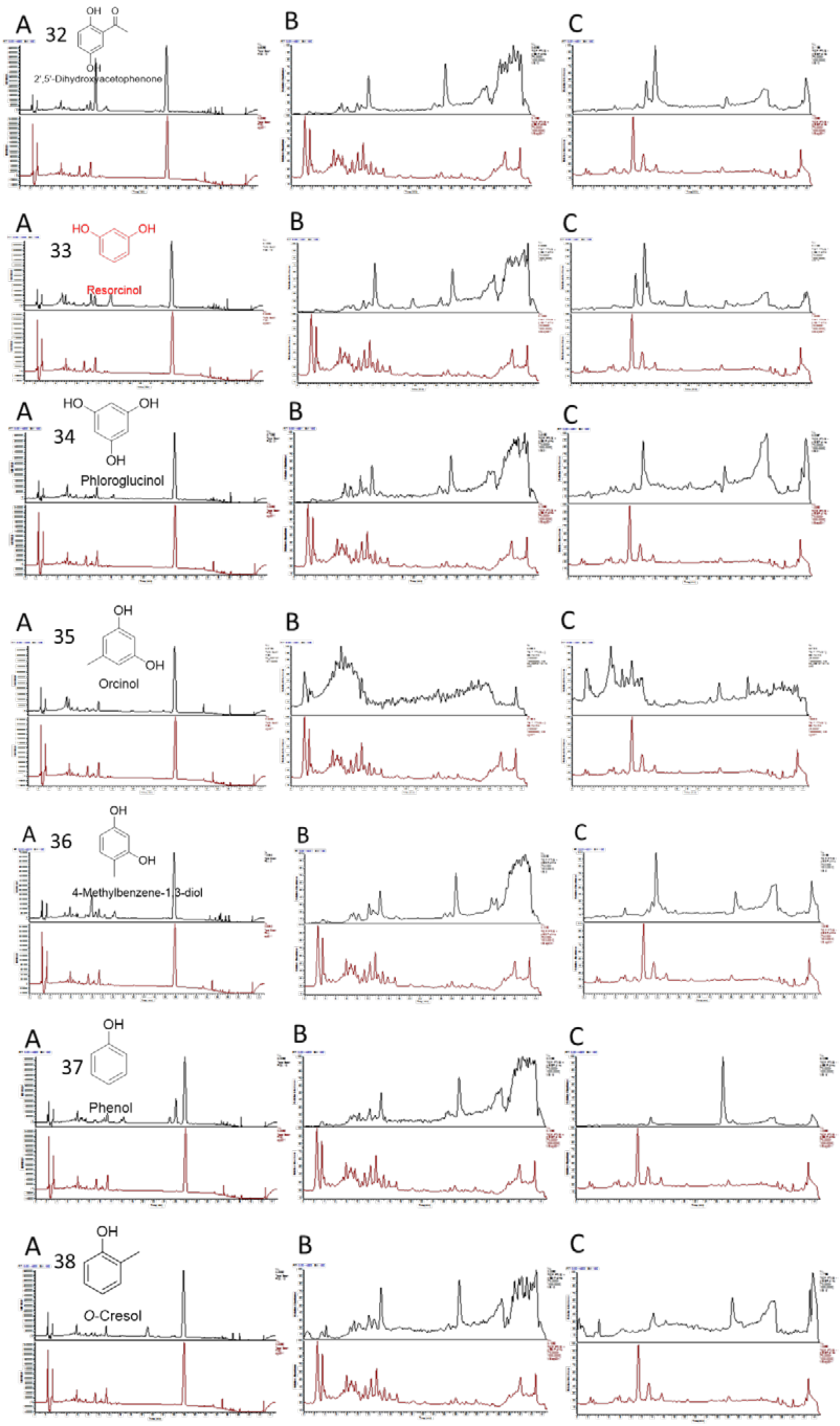


Figure S69. The LC-MS analysis of *E. coli* fed with compounds **32-38**. A: PDA analysis; B: Positive MS analysis; C: B: Negative MS analysis.

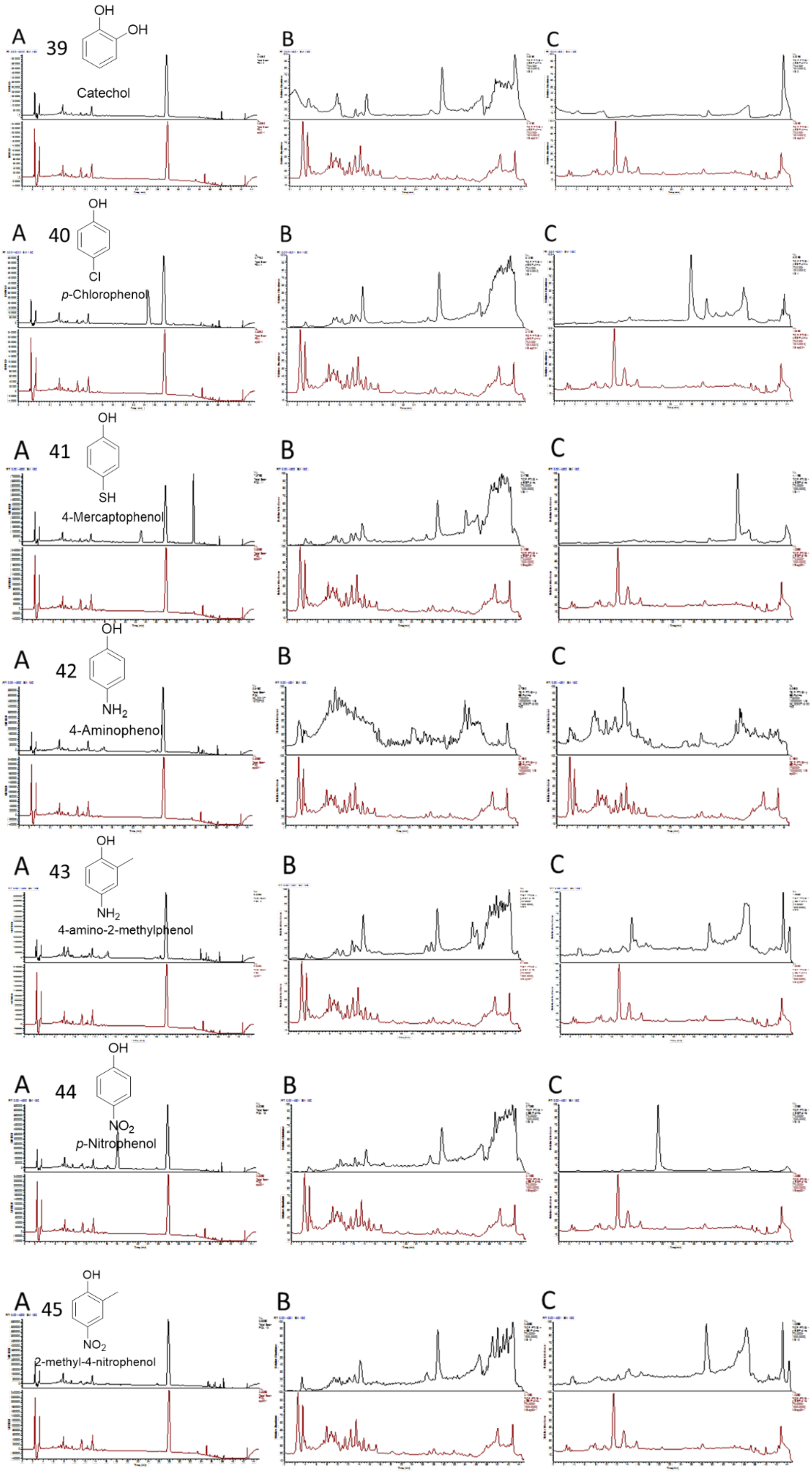


Figure S70. The LC-MS analysis of *E. coli* fed with compounds **39-45**. A: PDA analysis; B: Positive MS analysis; C: B: Negative MS analysis.

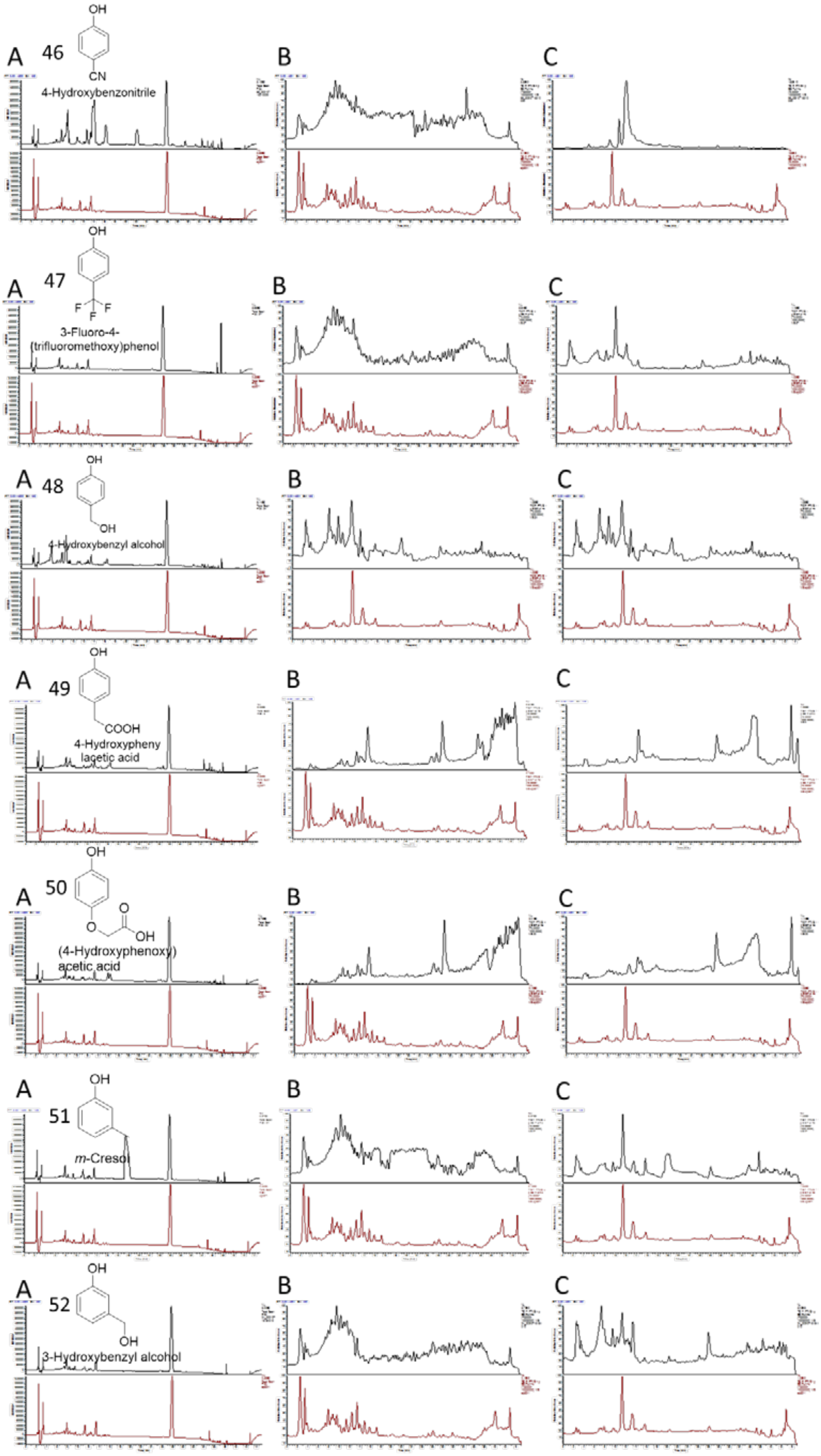


Figure S71. The LC-MS analysis of *E. coli* fed with compounds **46-52**. A: PDA analysis; B: Positive MS analysis; C: B: Negative MS analysis.

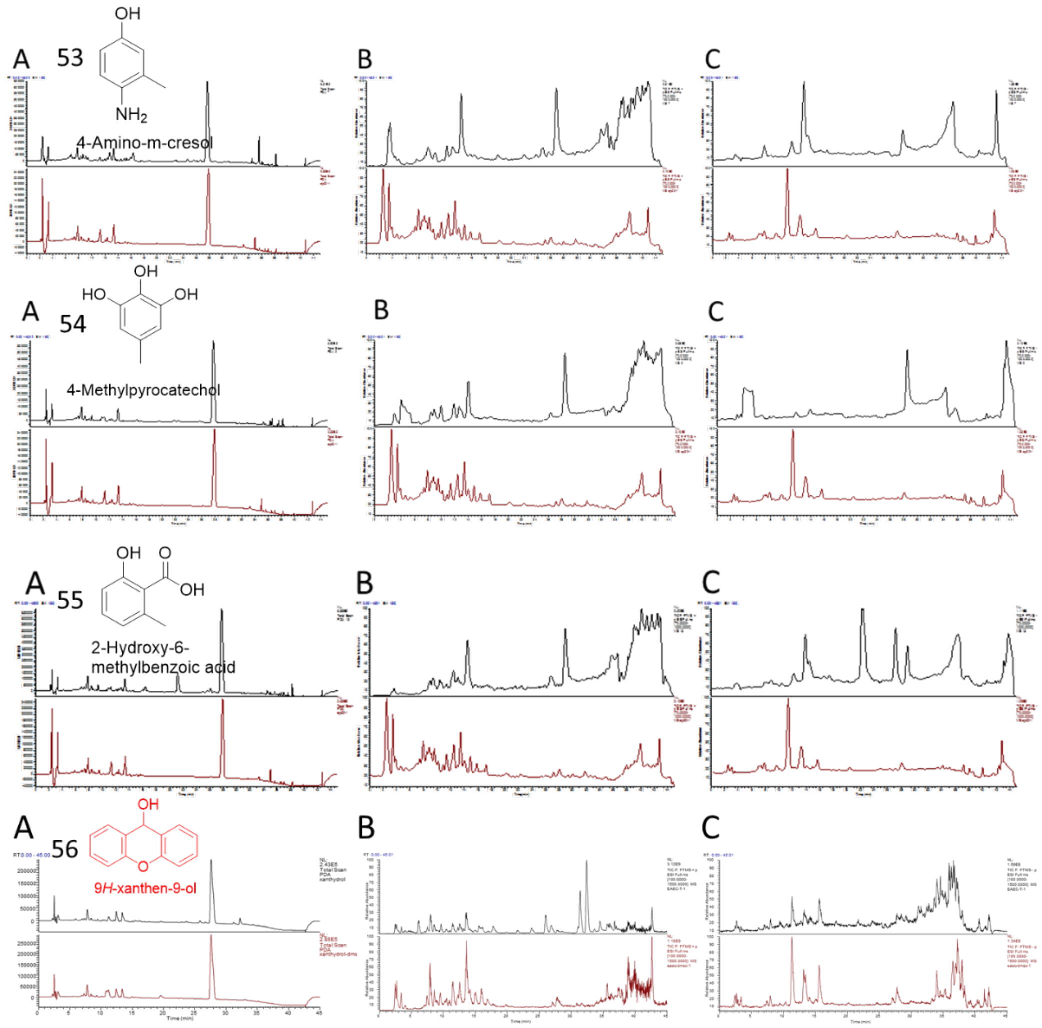


Figure S72. The LC-MS analysis of *E. coli* fed with compounds **53-56**. A: PDA analysis; B: Positive MS analysis; C: B: Negative MS analysis.

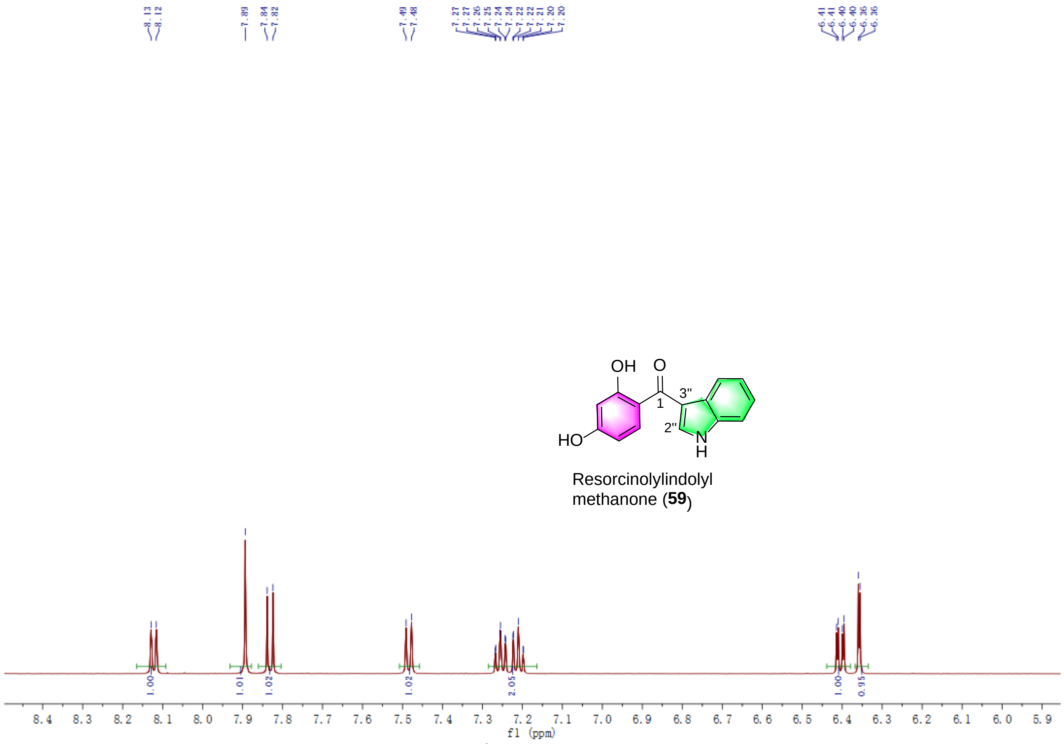


Figure S73. The $^{1}$H NMR spectrum of **59.**

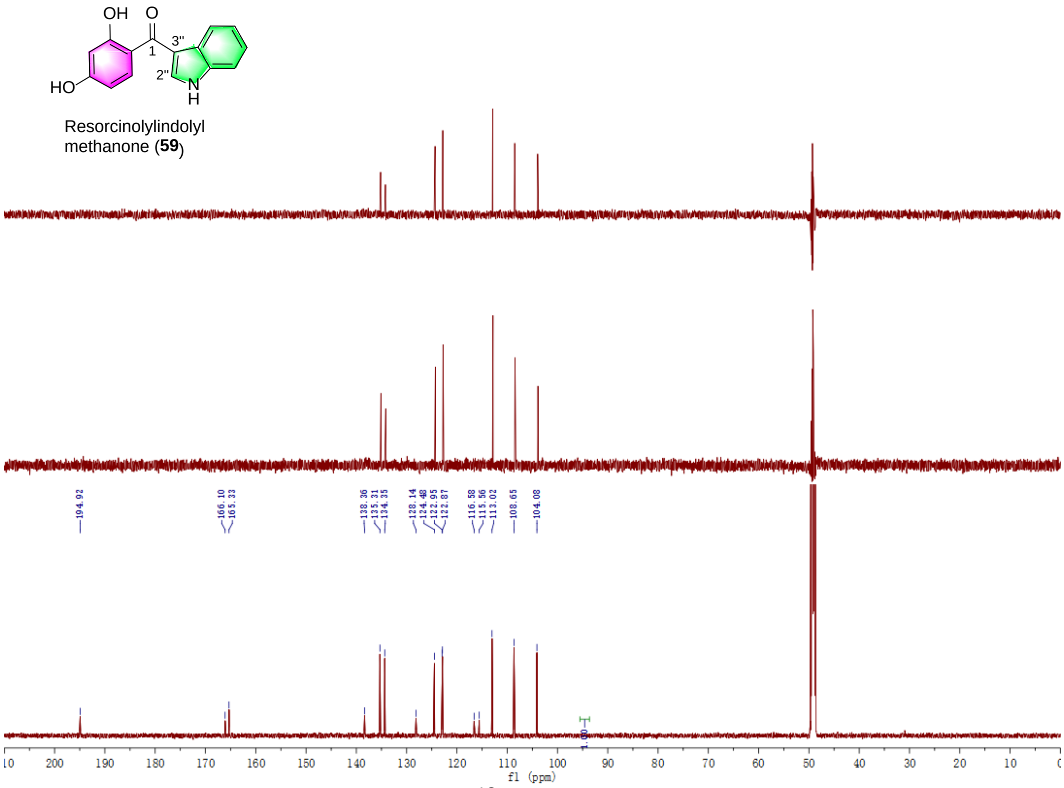


Figure S74. The $^{13}$C NMR spectrum of **59.**

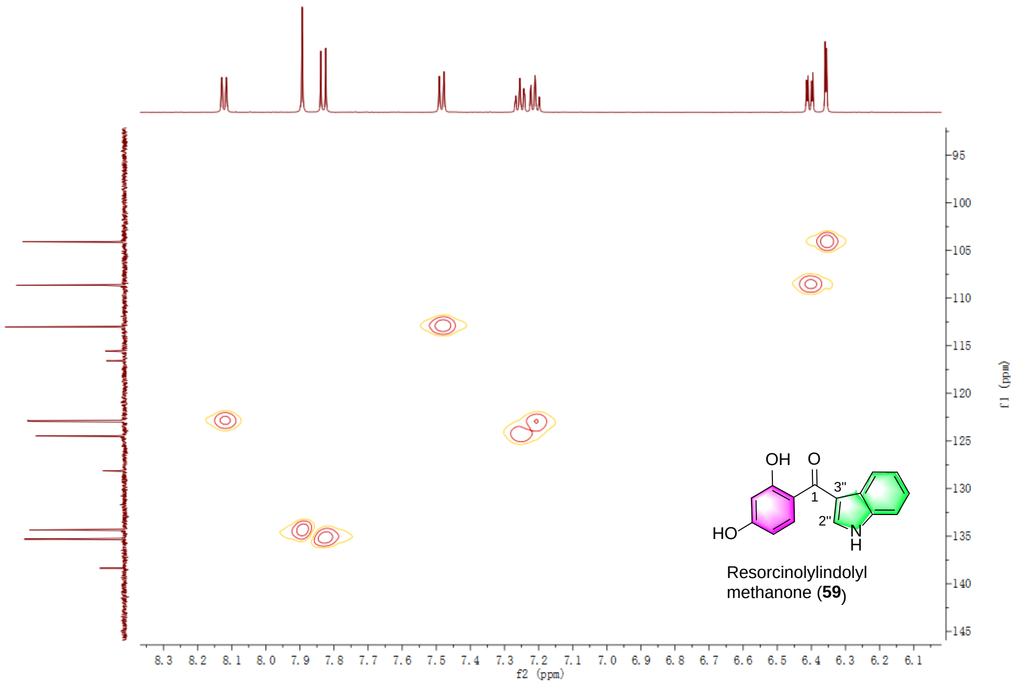


Figure S75. The HSQC spectrum of **59.**

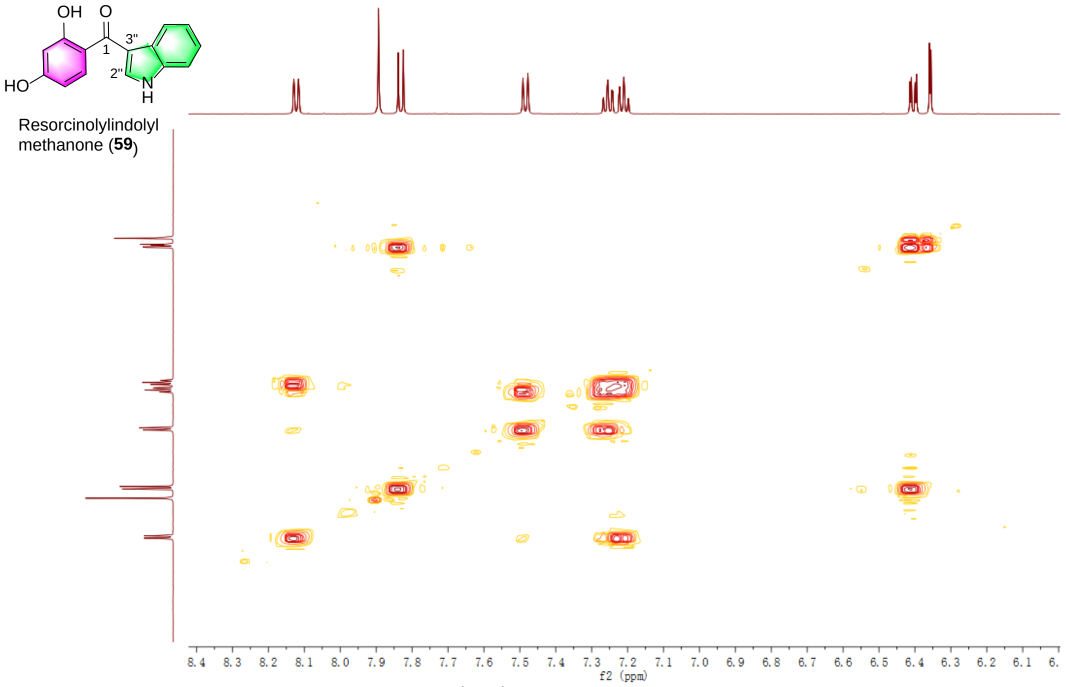


Figure S76. The $^{1}$H-$^{1}$HCOSY spectrum of **59.**

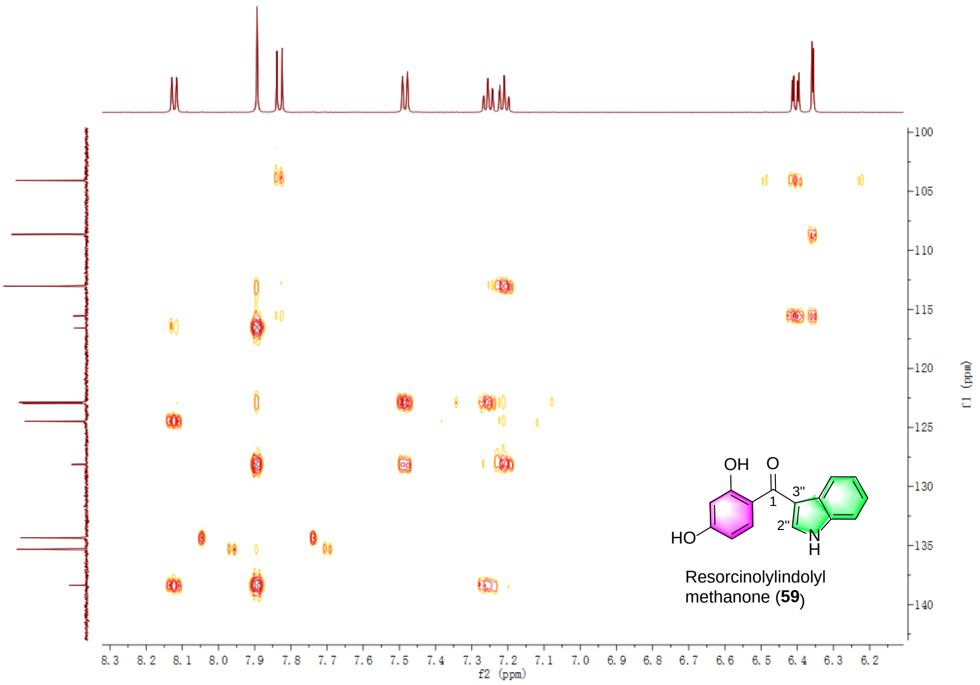


Figure S77. The HMBC spectrum of **59.**

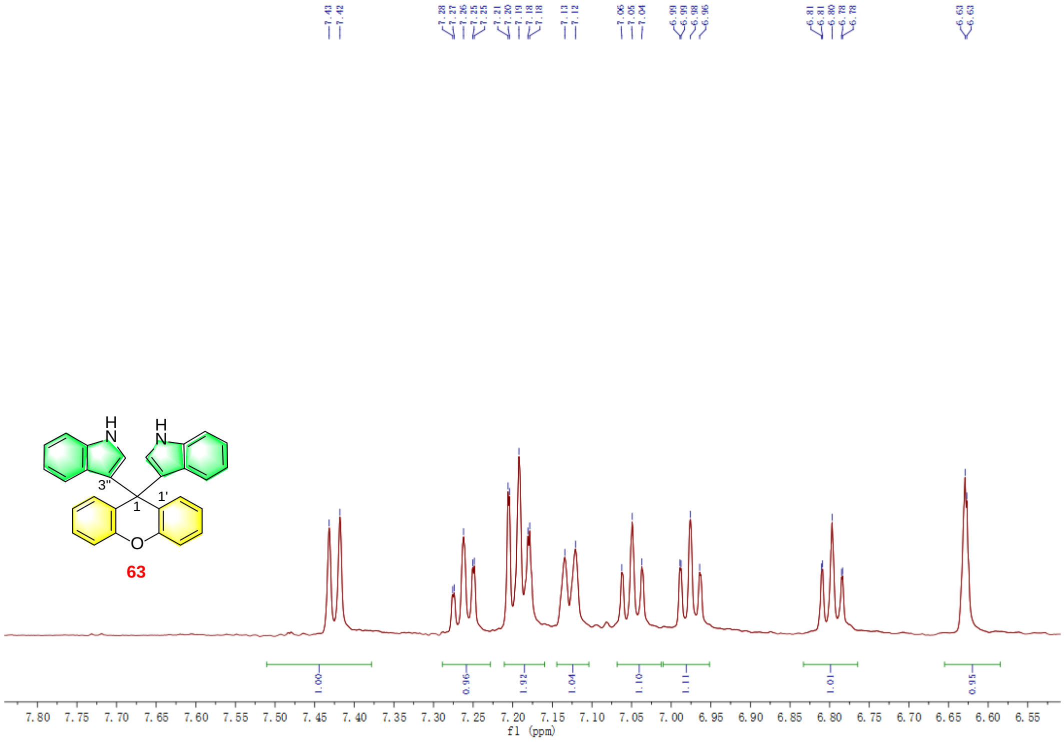


Figure S78. The $^{1}$H NMR spectrum of **63.**

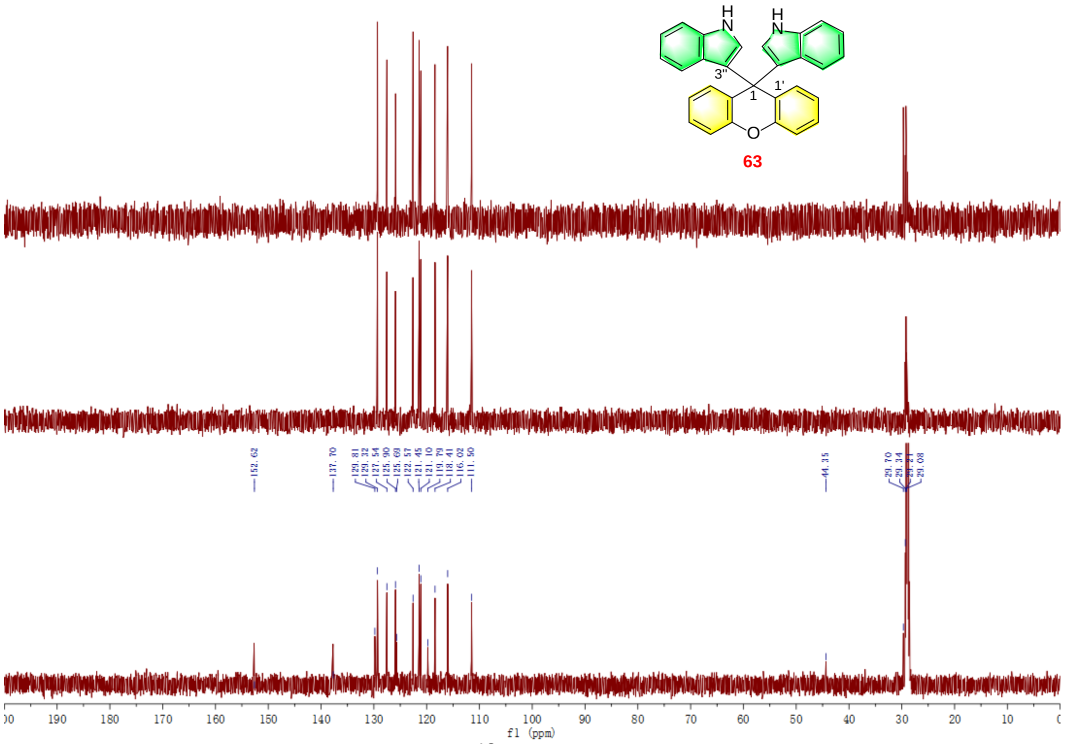

Figure S79. The $^{13}C$ NMR spectrum of **63.**

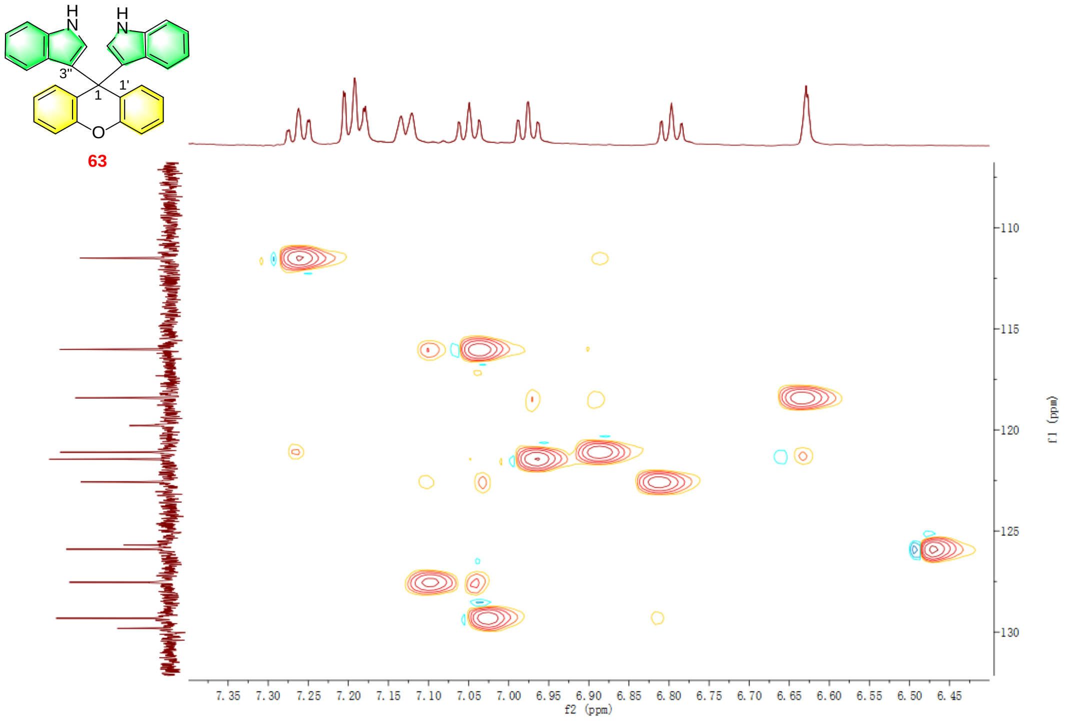

Figure S80. The HSQC spectrum of **63.**

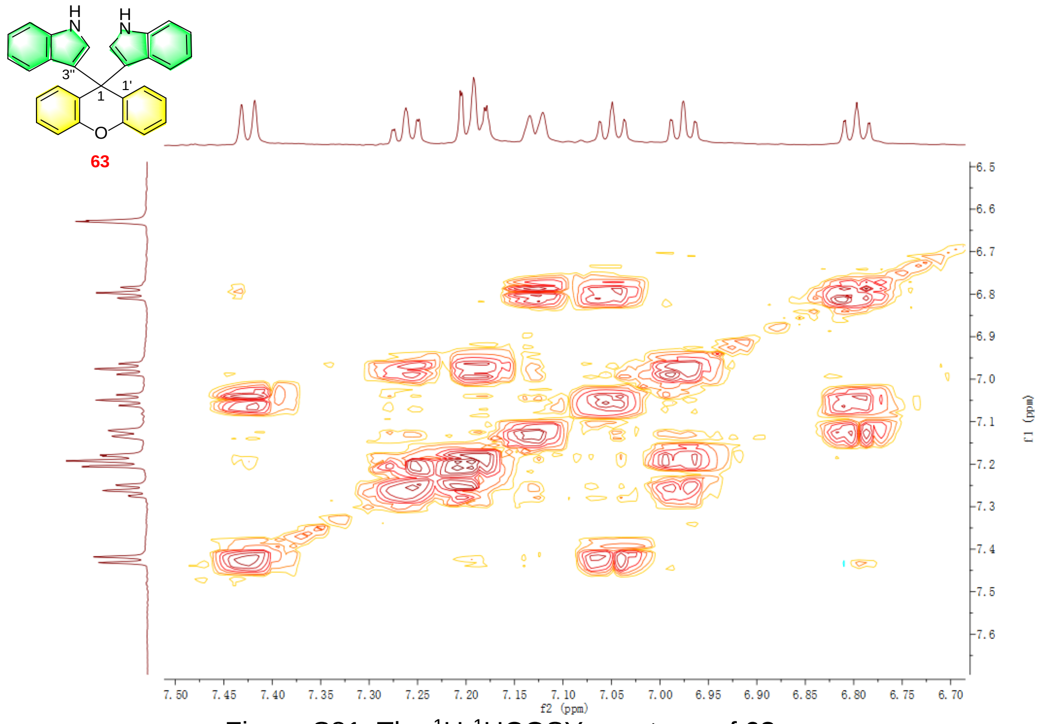


Figure S81. The $^1H$-$^1H$COSY spectrum of **63.**

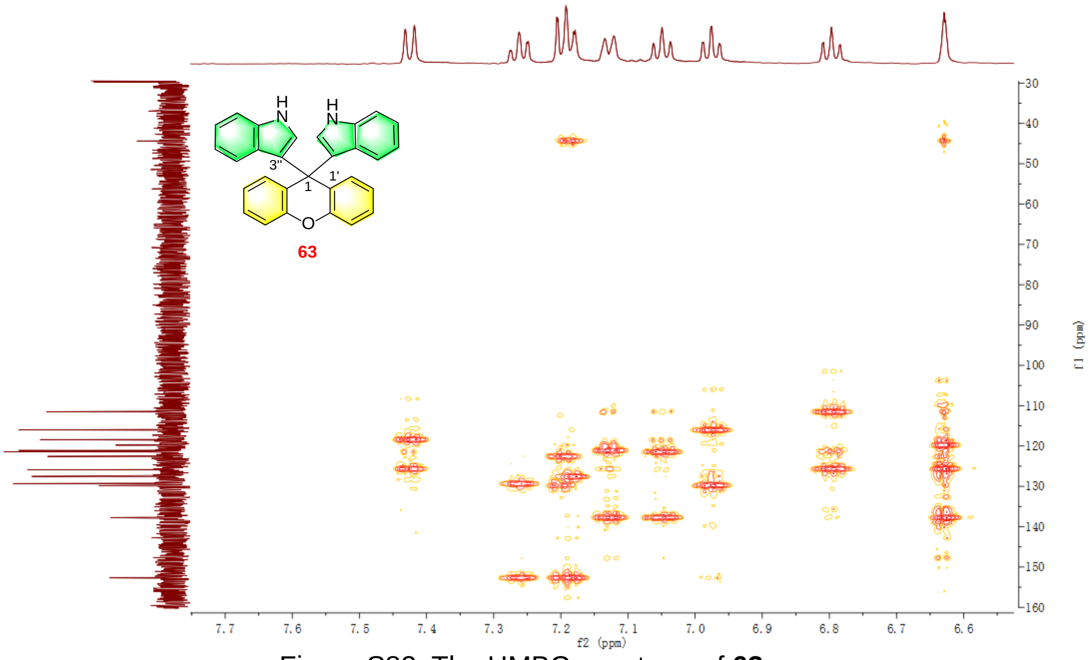


Figure S82. The HMBC spectrum of **63.**

Crystal data for gecb1 (Arthrocolin J, **10**): $C_{29}H_{23}NO_4 \bullet 2(CH_4O)$, $M$ = 513.57, $a$ = 9.0191(4) Å, $b$ = 20.9712(8) Å, $c$ = 13.7202(6) Å, $\alpha$ = 90°, $\beta$ = 96.485(2)°, $\gamma$ = 90°, $V$ = 2578.45(19) Å$^3$, $T$ = 100.(2) K, space group $P121/c1$, $Z$ = 4, $\mu$(Cu K$\alpha$) = 0.745 mm$^{-1}$, 74199 reflections measured, 5079 independent reflections ($R_{int}$ = 0.2421). The final $R_1$ values were 0.1083 ($I > 2\sigma(I)$). The final $wR(F^2)$ values were 0.3302 ($I > 2\sigma(I)$). The final $R_1$ values were 0.1357 (all data). The final $wR(F^2)$ values were 0.3469 (all data). The goodness of fit on $F^2$ was 1.319.

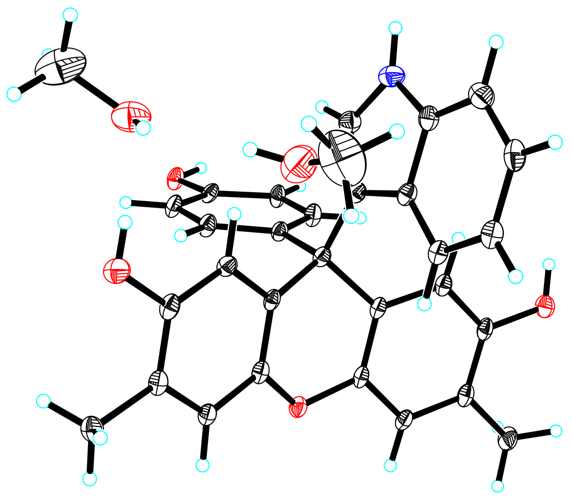

View of the molecules in an asymmetric unit.
Displacement ellipsoids are drawn at the 30% probability level.

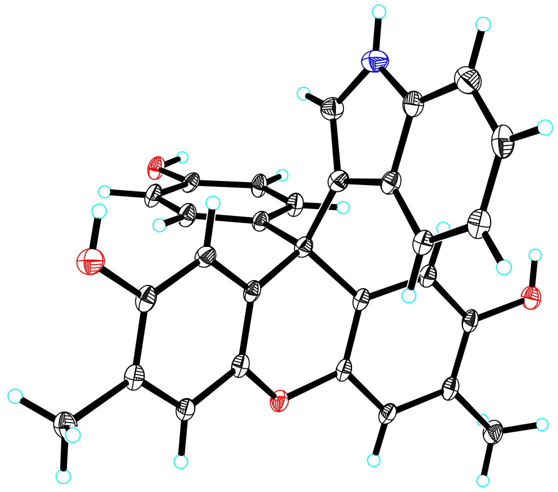

View of a molecule of gecb1 (Arthrocolin J, **10**) with the atom-labelling scheme.
Displacement ellipsoids are drawn at the 30% probability level.

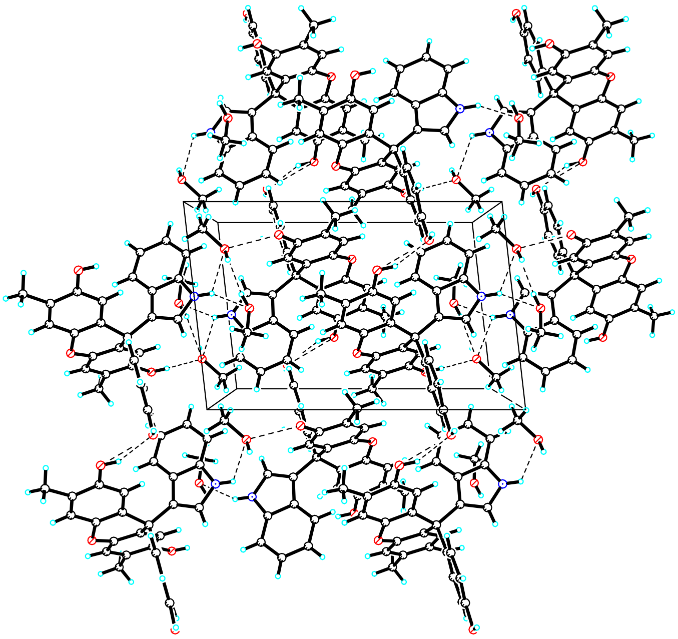

View of the pack drawing of gecb1 (Arthrocolin J, **10**).

Hydrogen-bonds are shown as dashed lines.

Crystal data for gecb2 (Arthrocolin H, **11**): $C_{29}H_{23}NO_4$, $M$ = 449.48, $a$ = 13.9126(10) Å, $b$ = 14.7966(11) Å, $c$ = 13.9592(14) Å, $\alpha$ = 90°, $\beta$ = 104.623(5)°, $\gamma$ = 90°, $V$ = 2780.5(4) Å$^3$, $T$ = 100.(2) K, space group $P$121/$c$1, $Z$ = 4, $\mu$(Cu Kα) = 0.577 mm$^{-1}$, 42430 reflections measured, 5284 independent reflections ($R_{int}$ = 0.3475). The final $R_1$ values were 0.1258 ($I > 2\sigma(I)$). The final $wR(F^2)$ values were 0.3260 ($I > 2\sigma(I)$). The final $R_1$ values were 0.2029 (all data). The final $wR(F^2)$ values were 0.3828 (all data). The goodness of fit on $F^2$ was 1.096.

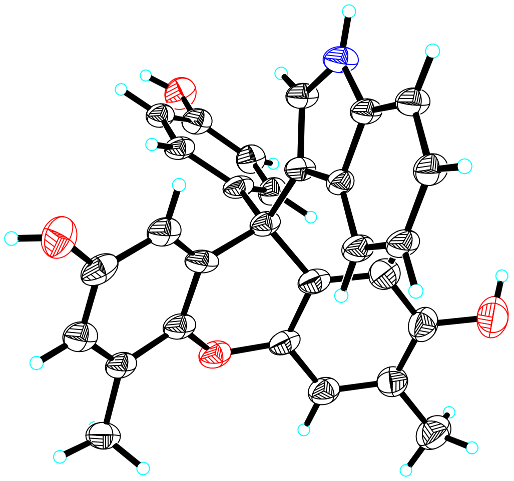

View of a molecule of gecb2 (Arthrocolin H, **11**) with the atom-labelling scheme. Displacement ellipsoids are drawn at the 30% probability level.

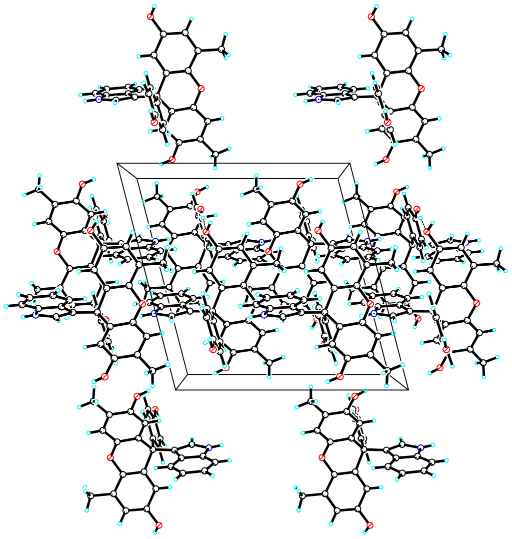

View of the pack drawing of gecb2 (Arthrocolin H, **11**). Hydrogen-bonds are shown as dashed lines.

Crystal data for gecb25 (**17**): $C_{15}H_{13}NO_2$, $M$ = 239.26, $a$ = 6.0106(3) Å, $b$ = 18.5798(9) Å, $c$ = 11.3684(5) Å, $\alpha$ = 90°, $\beta$ = 96.0690(10)°, $\gamma$ = 90°, $V$ = 1262.46(10) $Å^3$, $T$ = 102.(2) K, space group $P121/n1$, $Z$ = 4, $\mu$(Cu Kα) = 0.677 $mm^{-1}$, 19889 reflections measured, 2480 independent reflections ($R_{int}$ = 0.0696). The final $R_1$ values were 0.0477 ($I > 2\sigma(I)$). The final $wR(F^2)$ values were 0.1237 ($I > 2\sigma(I)$). The final $R_1$ values were 0.0496 (all data). The final $wR(F^2)$ values were 0.1259 (all data). The goodness of fit on $F^2$ was 1.046.

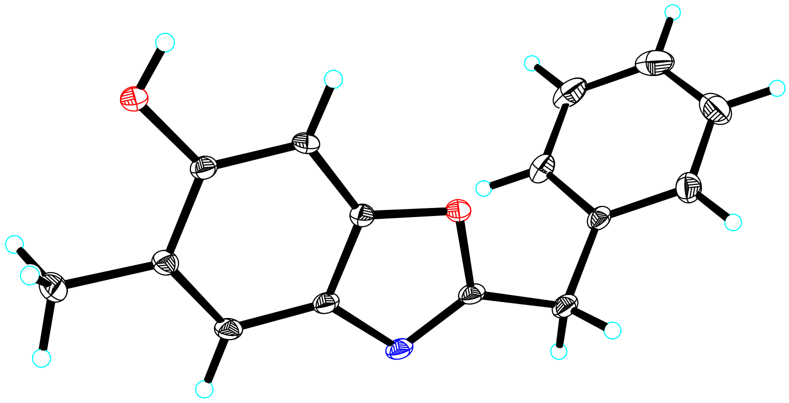

View of a molecule of gecb25 (**17**) with the atom-labelling scheme. Displacement ellipsoids are drawn at the 30% probability level.

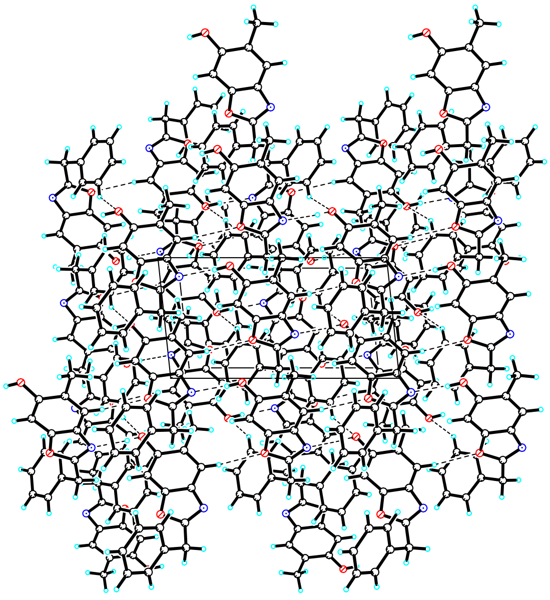

View of the pack drawing of gecb25 (**17**). Hydrogen-bonds are shown as dashed lines.
